# A diverse family of protein antibiotics inhibits the BAM complex by β-signal mimicry

**DOI:** 10.64898/2026.09.25.754295

**Authors:** Fabian Munder, Chunxiao Wang, Luis Jimenez, Matthew T. Doyle, James P. R. Connolly, Gavin J. Knott, Rhys Grinter

## Abstract

Lectin-like protein antibiotics (Llps) kill *Pseudomonas* by inhibiting BamA, the essential core of the β-barrel assembly machinery (BAM). Llps are composed of one or two β-lectin domains and a C-terminal peptide, with characterisation to date centred on the two-domain L-type pyocins of *P. aeruginosa*, which bind BamA extracellular loop 6 and inhibit by delivering the C-terminal peptide to BamA β-strand 1. Whether this mechanism is general, and how Llps contend with BamA sequence differences across species, remained unknown. We screened 238 *Pseudomonas* Llps against 101 genome-sequenced isolates, showing susceptibility across the family is set by BamA extracellular loop 6. Loop 6 length varies, and Llps fall into three targeting modes: short-loop and long-loop specialists and broad-range dual targeters, which have arisen repeatedly. We determine the 2.66 Å cryo-electron microscopy structure of a two-domain LlpA bound to BAM, showing recognition has been relocated relative to the L-type pyocins, with loop 6 bound by a single face of the N-terminal β-lectin domain rather than the inter-domain cleft. This explains how single-domain LlpBs retain BamA targeting and why the second lectin domain has been lost at least twice. We show that the BamA inhibition mechanism is shared across Llps, with a hypervariable C-terminal peptide binding BamA β-strand 1, inhibiting the BAM insertase by mimicking the β-signal of BamA substrates. Together, these data provide an atlas of Llp diversity and the mechanistic principles for engineering it, identifying Llps as a tuneable, multi-interface scaffold for BAM-directed precision antibacterials against pathogenic *Pseudomonas*, including economically important plant pathogens.

## Introduction

The rise of antimicrobial resistance among Gram-negative bacteria, in combination with a sparse antibiotic development pipeline, has generated significant interest in antibacterials that act through unexploited targets and mechanisms^1,2^. One such target is the Gram-negative outer membrane, which acts as a formidable permeability barrier that excludes many antibiotics before they can reach their cellular targets^3,4^. The outer membrane also presents essential, surface-exposed biogenesis machinery that can be inhibited from outside the cell, negating the outer membrane permeability barrier^5^. BamA, the conserved and essential core of the β-barrel assembly machinery (BAM) that folds and inserts outer-membrane proteins, has consequently become a high-interest antimicrobial target^6–10^. Diverse BamA-inhibiting antimicrobials have been discovered or developed, including Lectin-like protein antibiotics (Llps), the macrocyclic peptides darobactin and dynobactin, and antibodies, defining several distinct modes of inhibition. In some cases, atomic-resolution structures have directly guided the development of narrow-spectrum derivatives^5,8,10–13^.

Bacteria have independently evolved diverse precision protein antimicrobials, deployed largely against the closely related bacteria that are their direct competitors^14–16^. Llps are a prominent example of bacterially produced protein antimicrobials. The prototypical Llp, LlpA, was first described in a rhizosphere *Pseudomonas* isolate and comprises two monocot mannose-binding lectin (MMBL, or β-lectin) domains followed by an unstructured C-terminal extension^17–19^. The family now extends to the single-lectin LlpBs of *Pseudomonas*, the dual-lectin L-type pyocins of *P. aeruginosa*, and more distantly related dual-lectin Llps from *Burkholderia* and *Xanthomonas*^19–24^.

Our previous work on L-type pyocins indicates that Llp killing proceeds through a sequence of molecular recognition steps, each performed by a distinct region of the protein. The C-terminal β-lectin domain first binds the extracellular lipopolysaccharide (LPS) O-antigen, concentrating and positioning the toxin at the cell surface^23^. The β-lectin domains then bind the polymorphic extracellular loop 6 of BamA. Finally, the C-terminal extension enters the BamA lumen and delivers a short C-terminal peptide that binds BamA β-strand 1, competitively inhibiting outer-membrane protein assembly by BAM^24^. BamA therefore serves as both the antibiotic target and a primary specificity determinant for Llps. Spontaneous Llp resistance maps to O-antigen loss and substitutions in the surface-exposed loop 6, and polymorphism of this loop tracks closely with susceptibility^24,25^. How BamA loop 6 contributes to the killing spectrum of individual Llps, alongside O-antigen recognition and other susceptibility determinants, remains unclear.

The modular structure of Llps makes them compelling starting points for the development of precision antibacterials. Their activity is potent and genus-restricted, inhibiting multidrug-resistant *P. aeruginosa* and plant-pathogenic pseudomonads at nanomolar concentrations^23^. Llps are effective for combatting *Pseudomonas* infections *in vivo*, with L-type pyocins able to clear *P. aeruginosa* lung infection in mice, and transgenic expression of LlpA confers resistance to *Pseudomonas syringae* in plants^26,27^. As narrow-spectrum agents, Llps have the potential to eliminate a defined pathogen while sparing the surrounding microbial community, an increasingly attractive property both in the clinic and in agriculture, where reliance on copper formulations and broad biocides drives resistance and collateral ecological harm^28,29^.

Despite this, our understanding of the family remains narrow. Functional characterisation has centred on a handful of *P. aeruginosa* L-type pyocins, and a few prototypical LlpAs and LlpBs, with structural information on Llp-BAM interactions available only for two L-type pyocins^21,24^. Llp genes are present in ∼10% of sequenced *Pseudomonas* and are enriched in plant-associated and soil-dwelling species, implying an ecological role in intra-genus competition that remains poorly defined^21^. Initial sequence analyses place LlpAs from environmental pseudomonads in phylogenetic clades distinct from the L-type pyocins of *P. aeruginosa*, and the single-lectin LlpBs widen the family further beyond the canonical dual-lectin architecture^21^. The evolutionary relationships between these types, and the rules linking their sequences and structure to target spectrum, are largely unexplored.

Here we determine the structural basis for BamA binding and inhibition by *Pseudomonas* LlpAs and LlpBs, placing it in the context of a genus-wide survey of the family. We combine phylogenomic analysis with a susceptibility screen of 238 Llps against 101 Pseudomonas strains, and integrate cryo-EM, structural modelling and BamA loop-variant analysis. Together, these data define the diversity and evolutionary relationships within the family and connect activity profiles to polymorphism in the BamA target, thereby establishing how LlpA/B and BamA sequences together contribute to spectrum, potency, and specificity. The result is an atlas of Llp diversity and a set of mechanistic principles for engineering, identifying Llps as a tuneable, multi-interface scaffold for the precision control of pathogenic *Pseudomonas*, including the economically significant plant pathogens of this genus.

## Main

### Lectin-like protein antibiotics are diverse and widespread in *Pseudomonas*

To identify lectin-like bacteriocins across the genus *Pseudomonas*, we searched the NCBI non-redundant protein database using characterised Llp sequences as queries and assembled a curated, dereplicated set of 238 Llps (Supplementary Data 1). Pairwise amino acid sequence identity ranged from 15 to 98%, capturing both closely related variants and deep sequence divergence across the family. To place these sequences within the wider Llp family and resolve their evolutionary relationships, we aligned the 238 sequences together with previously characterised *P. aeruginosa* L-type pyocins and representative characterised or putative Llps from *Burkholderia* (n = 14) and *Xanthomonas* (n = 7) species and inferred a maximum-likelihood phylogeny (Fig. 1a, Supplementary Data 1)^18,24,25,30,31^. The phylogeny resolved the *Pseudomonas* LlpA and LlpB sequences as a single large radiation, separated from the L-type pyocins and from the *Xanthomonas* and *Burkholderia* sequences, which each formed a monophyletic clade. Phylogenetic placement classified four of the Llps we identified with the L-type pyocins, forming a distinct 11-member clade, together with the seven previously characterised L-type pyocins. Our curated Llp set therefore comprises 234 LlpA/B sequences originating from non-*P. aeruginosa Pseudomonas* species and 4 L-type pyocins (Fig. 1a, Supplementary Data 1).

**Figure 1.**
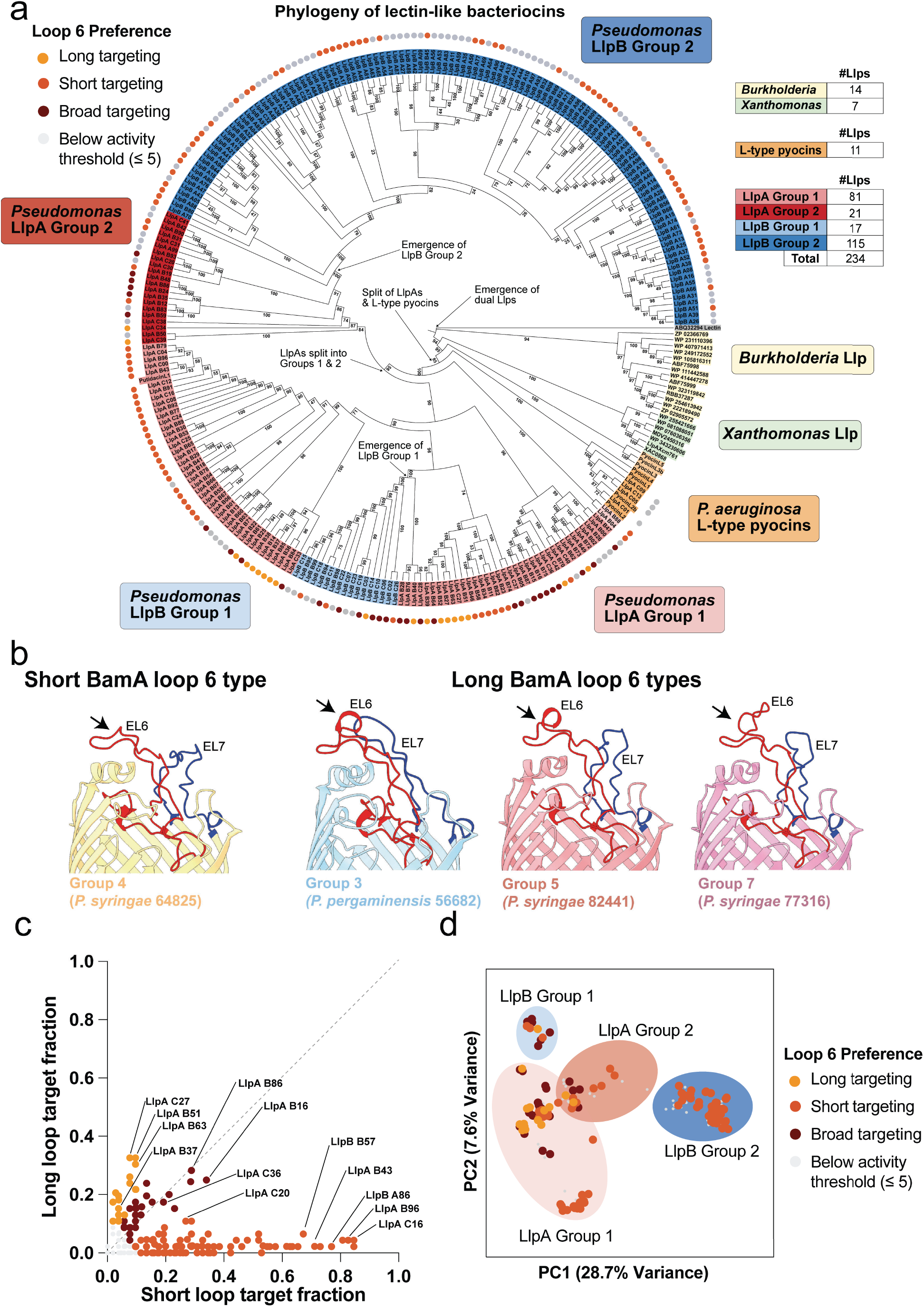
Diverse *Pseudomonas* LlpA/Bs show strong BamA loop 6-targeting preference. (a) A maximum-likelihood (ML) phylogeny of Llp proteins identified in the genomes of *Pseudomonas*, *Burkholderia*, and *Xanthomonas* species. *Pseudomonas* Llp groups (LlpA, LlpB, L-type pyocin) are shown, as well as the colony-screen-derived loop 6-targeting preference of Llps targeting diverse *Pseudomonas* strains, indicated as coloured circles. (b) A cartoon view of a selection of AlphaFold3 models of BamA, from the panel of *Pseudomonas* isolates, showing the predicted structure and topology of extracellular loop 6 and 7. The insertion location of long loop 6 variants is noted with an arrow. See Fig. S4 for models for each BamA loop 6 group identified in the isolate panel. (c) A scatter plot showing the fraction of long and short BamA loop 6-containing *Pseudomonas* strains targeted by the LlpA/Bs in our binary activity screening assay. (d) A principal component analysis (PCA) plot of LlpA/B sequence diversity, annotated with phylogenetic group and BamA loop 6-targeting preference. LlpA/B sequences are colour-coded based on BamA loop 6-targeting preference, or activity below the sensitivity threshold for the binary screening assay.

Several features of this phylogeny are worth emphasising. The 238 sequences in our dataset greatly outnumber the previously characterised Llps (4 sequences) and L-type pyocins (7 sequences), indicating that work to date has focused on a limited number of examples of a much larger and largely unexplored family^21,24,25^. The L-type pyocins emerge as a sister lineage to the larger LlpA/B radiation, consistent with their shared origin in *Pseudomonas*. *Burkholderia* and *Xanthomonas* sequences are further separated and monophyletic, consistent with Llp families having co-radiated within their host genera (Fig. 1a).

Within the *Pseudomonas* Llp radiation, the dual-lectin LlpA sequences resolved into two principal groups (LlpA Group 1 and LlpA Group 2) together with a two-member, basal LlpA lineage (Group 0). The single-lectin LlpBs form two groups (LlpB Group 1 and LlpB Group 2), with LlpB Group 2 by far the largest group in the dataset (n = 115) (Fig. 1a). A striking feature of the tree is the presence of two separate LlpB clades, each nested within, and arising from, dual-lectin LlpA ancestry. Because the dual-lectin architecture is ancestral to the whole Llp radiation, this topology indicates that single-domain LlpBs have arisen on at least two independent occasions through the loss of one β-lectin domain. Multiple-sequence alignment shows that the lost module is the C-terminal β-lectin domain, which predominantly mediates recognition of the LPS O-antigen in dual-lectin Llps (Fig. S1)^23^. Despite this architectural simplification, the size of LlpB Group 2 indicates that C-terminal β-lectin domain loss is a successful evolutionary innovation in *Pseudomonas* Llps. Whether LlpBs engage LPS O-antigen in the absence of the canonical C-terminal LPS-binding β-lectin domain is unclear.

### Broad screening of LlpA/Bs against diverse *Pseudomonas* isolates

To assess the activity of the 238 Llps against diverse hosts, we assembled a panel of 111 *Pseudomonas* isolates from the Australian Plant Pathology & Mycology Herbarium, chosen to maximise diversity of assigned species, host of origin, and geographic origin within Australia. Each isolate was genome-sequenced and draft-assembled, and new species assignments were made with GTDB-Tk (Supplementary Data 2).

Genes encoding all 238 Llps were synthesised and cloned for expression in *Escherichia coli*. To establish their inhibition profile, each Llp-expressing *E. coli* strain was arrayed as a discrete colony on nutrient agar in microplate (96-colony) format across three plates (Supplementary Data 3). Colonies were grown and lysed *in situ* to release the expressed Llp, after which the array was overlaid with *Pseudomonas-*seeded agar using isolates from our collection. Following incubation to establish confluent growth, zones of inhibition surrounding a colony were recorded as a binary measure of activity for the Llp against that strain (Fig. S2). Repeating this across the strain panel assembled an Llp × strain susceptibility matrix (Supplementary Data 4). The assay was noisy. False negatives likely occurred when Llp expression in *E. coli* was at low levels; false positives likely occurred due to inconsistent growth of the strain tested. In addition, large inhibition zones of one Llp at times encroached on neighbouring colonies, meaning not every Llp could be scored against every strain. Based on the screening results, 10 of the 111 strains were excluded due to poor growth or widespread, non-specific inhibition. The remaining 101 strains nonetheless yielded a large, analysable dataset of 23,406 scored Llp-strain observations, which we used to dissect the molecular determinants of Llp killing across the strain library (Supplementary Data 5).

### BamA loop 6 variation is a key determinant of Llp susceptibility

We previously demonstrated that BamA is the molecular target of L-type pyocins^24^. Given their shared ancestry and architecture, we hypothesised that LlpA/Bs also target BamA and, like L-type pyocins, initially bind to extracellular loop 6 of BamA before delivering a C-terminal peptide to inhibit the BAM complex^24^. In support of this hypothesis, previous evidence indicates that BamA loop 6 sequence contributes to susceptibility across the wider Llp family^24,25^. To further test this relationship, we analysed our screen data to determine how BamA loop 6 sequence correlates with Llp susceptibility across our panel of *Pseudomonas* isolates. We retrieved the BamA sequence from the genome of each *Pseudomonas* isolate in our screen and constructed a multiple-sequence alignment (Supplementary Data 6). Sequence variability was largely confined to the extracellular loop 6 and loop 7 regions, which were hypervariable (Fig. S3, Supplementary Data 6). Quantified across the 111 isolate BamA sequences, BamA is otherwise highly conserved, with a mean per-position identity of 95.5% outside the loop 6 and loop 7 segments, whereas the fifteen loop 6 residues (residues 666-680) average only 21.2% identity (Supplementary Data 6). We clustered the loop-6 sequences, allowing us to assign BamA sequences to 15 loop types comprising 41 unique loop 6 sequences (Fig. S3). These loop 6 sequence groups fall into two broad classes, which we classified as ‘long’ and ‘short’, with long loops distinguished by a 4-7 amino-acid insertion of variable sequence in the extracellular portion of the loop. AlphaFold3 modelling of BamA from strains with long and short loop types indicates a clear difference in loop 6 structure based on this insertion (Fig. 1b, Fig. S4, Supplementary Data 7)^32^. Across the 101 strains assessed for Llp susceptibility, these classes were near-evenly represented, with 53 short, 45 long, and 3 outgroup strains. The three outgroup strains belonged to *P. aeruginosa* or *Pseudomonas oryzihabitans* and had significantly more divergent BamA loop 6 sequences (Fig. S3, Supplementary Data 4 and 6). Mapping the loop types onto a GTDB-Tk-generated phylogeny of the isolates showed that loop type is broadly ancestral, with some switching of long and short loops within clades (Fig. S5)^33^.

To characterise loop 6 preference despite noise in individual inhibition calls, we excluded 90 low-activity Llps and classified the remaining 148 by their relative activity against long-and short-loop strains, retaining a broad-targeting category for Llps active against both classes (Methods; Supplementary Data 5 and 8). Of the 148 confidently active Llps, 99 were short loop 6-targeting, whereas 19 were long loop 6-targeting and 30 had broad targeting across both loop types (Fig. 1a,c,d). Short loop-targeting is therefore the dominant mode of activity in the family, with the variable 4-7-residue-long loop insertion possibly acting as a barrier to loop 6 interaction that comparatively few Llps overcome.

The difference in breadth of activity between these classes likely reflects the architecture of their loop 6 target. Short-preferring Llps recognise loop 6 with no insertion, resulting in a loop structure common to all short-loop strains, inhibiting a median of 14 strains, and up to 44 (83%) of short-loop strains (Fig. 1c). Long-preferring Llps face a more difficult recognition problem because the long loop 6 insertion varies in both length and sequence, meaning it is unlikely that a single Llp can bind every long-loop variant (Fig. 1b, Fig. S4, Supplementary Data 7). Consistent with this, long loop 6 targeting Llps were narrower in spectrum (median 11 strains) and were themselves sub-specialised; none was active against all seven long loop 6 groups, and their activity fell largely along two insertion families, one comprising the large group-03 insertion and the other the related group-05/09/10 insertions. Across all active Llps that engage long loop 6 types (long and broad targeting groups), these two insertion families were strongly anti-correlated (Spearman r = −0.67, P < 0.001, n = 47), indicating that recognition of one family rarely accompanies recognition of the other (Fig. S6, Supplementary Data 8). Even the most active long-targeting LlpA/Bs (e.g. LlpA-C27, LlpA-B51 and LlpA-B63) achieved their breadth of activity by targeting strains within the group-05/09/10 cluster. The 30 broad-range Llps killed strains of both loop classes (median 11.5 strains) and thus represent a distinct dual-targeting mode. The most active broad-range Llps (LlpA-B86 and LlpA-B16) killed close to 28 strains spanning long-and short-loop types, indicating a flexibility in loop 6 targeting not observed in long-or short-loop 6 targeting Llps.

Loop 6 specialisation mapped cleanly onto Llp phylogeny (Fig. 1a). The largest group, LlpB Group 2, appears to be an exclusively short-loop-targeting lineage (57 short-preferring and 58 below-threshold members). LlpA Group 1, by contrast, contained most of the long loop 6-preferring Llps (16 of 19) and broadly active Llps (18 of 30) in addition to a clade of highly active dedicated short loop 6 specialists. The smaller LlpA Group 2 and LlpB Group 1 both contained all three loop 6 targeting groups (Fig. 1a). All four L-type pyocins in our panel fell below the activity threshold, suggesting they are inactive across our environmental *Pseudomonas* isolates, consistent with the *P. aeruginosa*-restricted activity of this Llp group (Supplementary Data 5 and 8).

Finally, because loop 7 lies adjacent to loop 6 and is also hypervariable, we asked whether it also contributes to Llp susceptibility (Fig. S3, Supplementary Data 6). Mantel tests showed that variation in susceptibility was correlated with sequence variation in both loop 6 (r = 0.48) and loop 7 (r = 0.32; both P < 0.001). However, the two loops are themselves strongly correlated in sequence (r = 0.63), reflecting their structural adjacency and shared inheritance, so we assessed each association while controlling for variation in the other loop (partial Mantel tests; Methods). Controlling for loop-6 sequence abolished the loop 7 correlation entirely (r = 0.03, P = 0.51), whereas loop 6 retained its correlation when controlling for loop 7 (r = 0.38, P < 0.001) (Table S1). Loop 7 variation therefore appears to carry no association with susceptibility independent of loop 6 in our screen.

### Purified LlpA/B activity confirms BamA loop 6-targeting

Next, we sought to confirm the activity spectra of LlpA/Bs as purified proteins. Selected LlpA/Bs were expressed with an N-terminal 6×His affinity tag and affinity purified. We tested 21 LlpA/Bs, 20 from our collection together with the previously characterised Putidacin L1, against ten *Pseudomonas* isolates spanning both short and long loop 6 types, eight of which were also present in the colony screen. Representatives of each LlpA/B activity class were selected from the screen, including two of the most active short loop 6-targeting (LlpA-C16 and LlpB-A86), two broad loop 6-targeting (LlpA-B16 and LlpA-C36), and two long loop 6-targeting (LlpA-C27, LlpA-B51) LlpA/Bs (Fig. 1c). Purified Llps were assayed by soft agar overlay, reading zones of inhibition after 24 h. Each zone was scored visually for size and clarity on a scale from 0 (no inhibition) to 5 (large, clear zone; Fig. 2a, Fig. S7). The assay was performed in three independent repeats, with good agreement between repeats (Supplementary Data 9).

**Figure 2.**
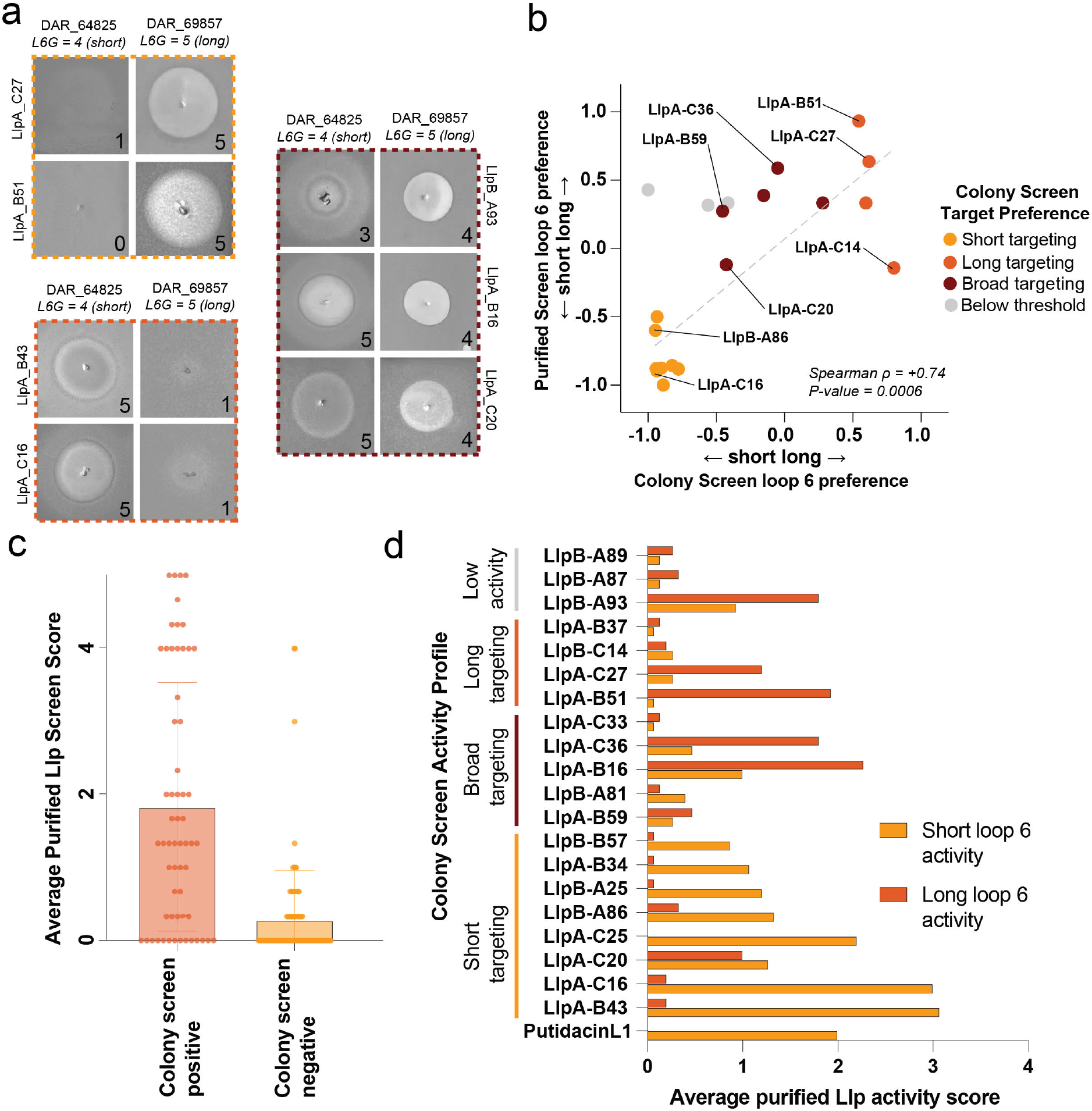
Purified LlpA/B activity validates loop-6 preferences inferred from the colony screen. (a) Representative inhibition zones from the plate-based activity assay of purified LlpA/Bs screened against long and short BamA loop 6-containing *Pseudomonas* isolates. This experiment was repeated three times (n = 3, biological replicates), with a single representative image shown. See Fig. S7 for representative images of a repeat of the full screen. (b) A scatter plot showing the correlation between the BamA loop 6 preference index (PI) of each Llp in the colony-based and purified protein screens, with dots colour-coded for BamA loop 6-targeting type in the colony screen. (c) A box-and-whisker plot showing the average activity of LlpA/Bs tested in the purified activity screen, separated into proteins that were active or inactive in the colony screen. (d) The average activity of the purified LlpA/Bs towards long and short loop 6 sequences, in comparison to their loop 6-targeting preference identified in the broad screen. PI values for representative Llps (purified-protein assay versus colony screen) were: LlpA-C16, −0.87 versus −0.90; LlpA-C25, −1.00 versus −0.89; LlpA-B43, −0.88 versus −0.94; LlpB-A86, −0.60 versus −0.95; LlpA-C27, +0.64 versus +0.62; LlpA-B51, +0.93 versus +0.54; LlpA-B16, +0.39 versus −0.15; and LlpA-C36, +0.59 versus −0.05. LlpA-B16 produced individual inhibition-zone scores of 4–5 against strains in both loop classes (Supplementary Data 10).

Agreement between the purified-protein assay and the original colony-based screen was good. Across the 158 Llp-strain pairs scored in both assays, 71% of binary inhibition calls were concordant (Cohen’s kappa = 0.50; ROC-AUC 0.78; Spearman ρ = 0.55 between consensus purified potency and binary colony screen call, P < 1 × 10^−1^^3^; Fig. 2c). The dominant class of disagreement was purified-positive/colony-negative (32 pairs, 28 of them with an average purified potency ≤ 1), consistent with weak inhibition falling below the detection threshold of the colony screen. LlpB-A93 is a clear example; it had strong and reproducible activity as a purified protein yet scored below threshold against these same strains in the colony screen, likely due to poor expression in this format (Fig. 2d). The smaller class of purified-negative/colony-positive observations (14 pairs) likely represent screen false positives.

The short, long and broad loop 6 preferences assigned from the colony screen were largely recapitulated by the purified proteins (Fig. 2b,d). We compared the assays using a normalised preference index (PI), ranging from −1 for exclusive short-loop activity to +1 for exclusive long-loop activity (Methods). Every screen-defined short loop 6 specialist retained this preference as a purified protein. LlpA-C27 and LlpA-B51 preferentially inhibited long-loop strains, while LlpA-B16 and LlpA-C36 retained activity against both loop classes despite shifts towards long-loop preference in the purified-protein assay (Fig. 2b,d; Supplementary Data 10).

Across the seventeen Llps active in the colony screen, PI correlated well between the two assays (Spearman ρ = +0.74, P = 0.0006; Fig. 2b), and for the twelve Llps with a clear preference in both assays (Methods), the direction agreed in every case. LlpB-C14 was an exception to the overall agreement in targeting class: it was a long loop 6 targeter in the colony screen but had a PI of ∼0 as a purified protein, indicating no strong loop preference. The remaining five Llps were broad targeters in the colony screen and retained robust activity against short and long loop 6 strains as purified proteins. These data confirm the central conclusion of the colony screen, that BamA loop 6 sequence partitions Llp target range into short, long and broad classes, with the residual differences dominated by screen false negatives rather than mis-assigned specificities, and they also provide a panel of validated LlpA/B-BamA pairs across the three classes for future engineering efforts.

### *Pseudomonas* Llps engage BamA loop 6 via a distinct binding mode

We previously determined cryo-EM structures of the L-type pyocins L1 and L2 in complex with their cognate BAM complexes from *P. aeruginosa*. In these structures, the pyocin engages BamA loop 6 via a cleft formed between its two β-lectin domains^24^. Given the distant evolutionary relationship between the L-type pyocins and other *Pseudomonas* Llps, we wanted to determine if *Pseudomonas* LlpA/Bs bound the BAM complex in the same way (Fig. 1a). As archetypes of the two loop 6 classes, we selected the BAM complexes of *P. putida* DAR64825 (BAM_64825_; short BamA loop 6 group 4) and *P. syringae* DAR82441 (BAM_82441_; long BamA loop 6 group 5) (Fig. 1b, Fig. S4). Both were expressed and purified recombinantly from *E. coli* and screened for stable complex formation against several LlpA and LlpB proteins that target these strains. The only complex obtained after size-exclusion chromatography was between the short loop BAM_64825_ and LlpA-C16 (Fig. 3a). We determined its structure by cryo-EM to 2.66 Å (PDB ID = 26HD) and the structure of BAM_82441_ alone to 2.36 Å (PDB ID = 26HC) to provide an experimental structure of the long loop 6 BAM complex for downstream comparison and modelling (Fig. 3b,c; Figs. S8–S10; Table S2).

**Figure 3.**
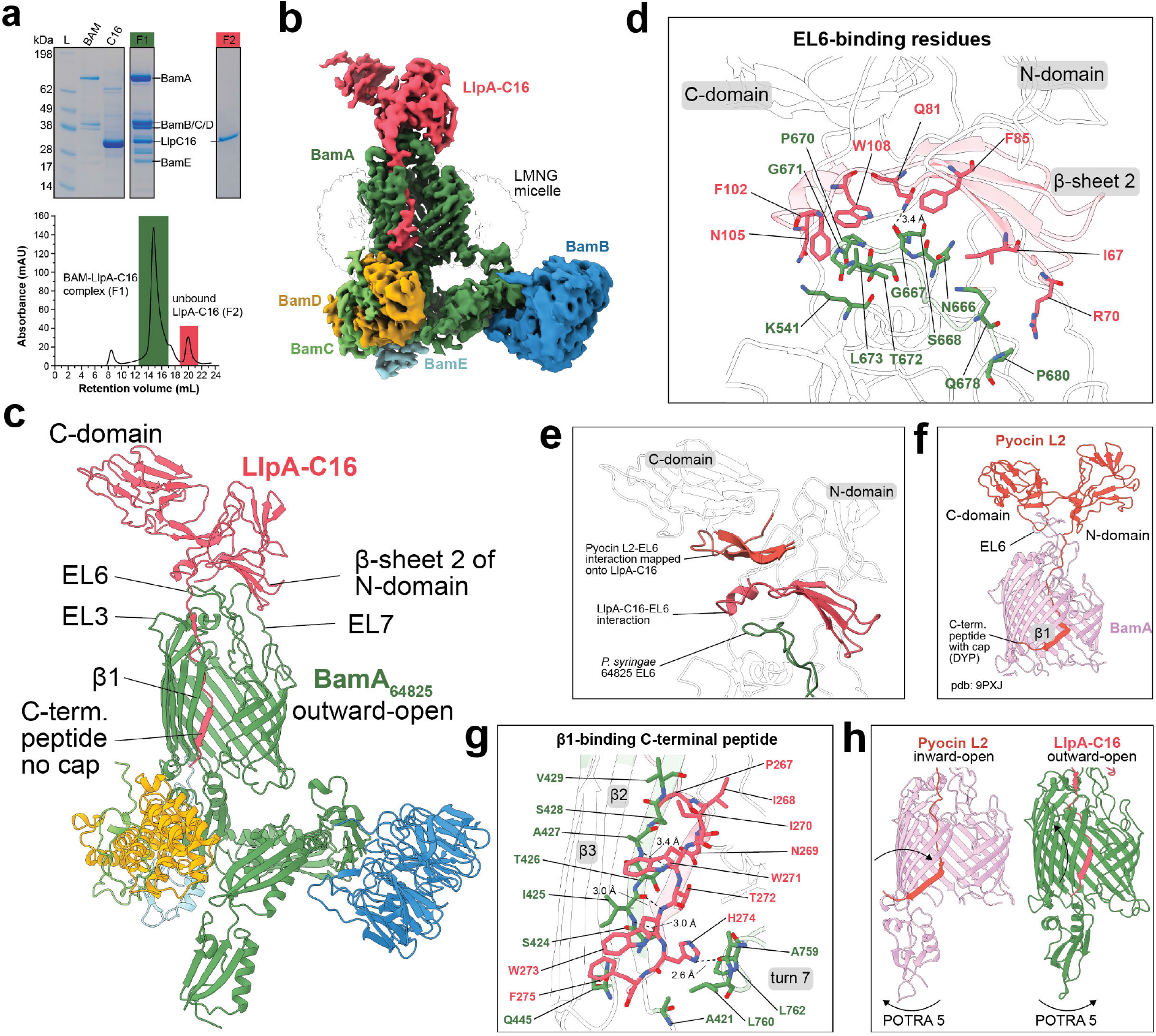
The structure of BAM_64825_-LlpA-C16 reveals LlpA/Bs bind BamA differently to L-type pyocins. (a) SDS-PAGE of purified BAM_64825_ and LlpA-C16 separately, and in complex (top) after isolation by size exclusion chromatography (SEC). SEC fractions and SDS-PAGE lanes corresponding to the BAM_64825_-LlpA-C16 complex and LlpA-C16 unbound are colour-coded for reference. (b) The cryo-EM density map of the BAM64825–LlpA-C16 complex, contoured at a threshold of 0.07, with LlpA-C16 and the BAM complex components colour-coded. (c) A cartoon representation of the BAM_64825_-LlpA-C16 complex, coloured as in panel b. (d) A zoomed cartoon view of the BAM_64825_-LlpA-C16 cryo-EM structure showing interaction between the LlpA-C16 N-terminal lectin domain and BamA loop 6, with key interacting residues shown as sticks. (e) A zoomed cartoon view of the BAM_64825_-LlpA-C16 cryo-EM structure showing the position of the LlpA-C16 BamA loop 6-binding interface, compared to the loop 6-binding interface from the cryo-EM structure of Pyocin L2. (f) A zoomed view of the pyocin L2-BAM_P28_ cryo-EM structure showing that Pyocin L2 interacts with BamA loop 6 via a cleft between its two β-lectin domains. (g) A zoomed cartoon view of the BAM_64825_-LlpA-C16 cryo-EM structure showing interaction between the LlpA-C16 C-terminal peptide and β-strand 1 at the lateral gate region of BamA, with key interacting residues shown as sticks. (h) Cartoon representation of the cryo-EM structures of BAM_64825_-LlpA-C16 and pyocin L2-BAM_P28_, showing that BamA in these complexes adopts distinct outward-open and inward-open conformations.

Consistent with our activity analysis, the BAM_64825_-LlpA-C16 cryo-EM structure shows that LlpA-C16 also engages BamA loop 6. However, it does so via an entirely different interface to the L-type pyocins. Rather than clasping loop 6 within an inter-domain cleft, LlpA-C16 binds via a face of the N-terminal β-lectin domain. The loop 6 residues of BamA (residues 666–680) are contacted exclusively by residues of the LlpA-C16 N-terminal β-prism (including I67, R70, Q81, F85, F102, N105 and W108), while the C-terminal, O-antigen-binding β-lectin domain makes no contact with BamA (Fig. 3c,d). This contrasts with the L-type pyocin-BAM complex, where loop 6 is gripped by residues drawn from both β-lectin domains of the pyocin (Fig. 3e,f). The N-terminal β-lectin domain binding reorients the Llp on BamA, positioning its C-terminal O-antigen-binding domain up and away from the membrane (Fig. 3c). Notably, all N-domain contacts with loop 6 are mediated by side-chain interactions (Tyr19, Ile67, Arg70, Gln81, Phe85, Leu96, Phe102, Asn105, and Trp108), so this interface can be readily tailored by mutation to recognise BamA loop 6 variants (Fig. 3d). BamA extracellular loop recognition is essentially confined to loop 6, with loop 7 not forming significant contacts with LlpA-C16, supporting our earlier analysis indicating loop 7 sequence does not determine Llp susceptibility. LlpA binding to loop 6 exclusively through its N-terminal β-lectin domain provides a structural rationale for how single-domain LlpBs achieve BamA loop 6 targeting: they retain the BamA-binding domain while lacking the C-terminal β-lectin domain required for the cleft-type binding mechanism of L-type pyocins. This also provides a rationale for the loss of the C-terminal β-lectin domain on two separate occasions, as it is dispensable for BamA binding in this branch of the Llp phylogeny (Fig. 1a).

Despite this divergent BamA extracellular loop 6-recognition between LlpA-C16 and Pyocin L2, the two families converge on the same inhibitory mechanism. The LlpA-C16 C-terminal peptide (residues 267–275) extends into the lumen of the BamA_64825_ β-barrel and binds β-strand 1, augmenting the barrel β-sheet. In contrast to the side-chain-specific EL6 interface, this pairing is made almost entirely through main-chain hydrogen bonds, consistent with the C-terminal peptide acting as a competitive inhibitor of the BAM β-barrel membrane protein substrates (Fig. 3c,g). Together with previous mutagenesis showing that deletion of the final two Llp residues abolishes antibacterial activity, this positioning strongly supports a conserved mechanism of BAM inhibition^30^. The LlpA-C16 inhibitory peptide shares significant similarity with pyocin L2, including an amphipathic structure, with hydrophobic side chains facing toward the membrane bilayer, and two centrally located bulky tryptophan residues. However, the extreme C-terminus of these peptides diverges, with LlpA-C16 terminating in a phenylalanine capping the C-terminal peptide, while the Pyocin L2 peptide has a histidine in this position followed by a DYP motif that extends towards the periplasm (Fig. 3g)^24^. Interestingly, while the C-terminal peptides of both LlpA-C16 and pyocin L2 bind β-strand 1 of BamA, the structure of BamA is resolved in different conformations in these complexes. LlpA-C16 binds in a β-strand 1 outward-open conformation, while pyocin L2 binds in an inward-open conformation (Fig. 3h).

### The LlpA/B-BamA loop 6 interface contributes to target specificity

To assess whether the BamA_64825_-LlpA-C16 binding mode is conserved, we modelled LlpA/Bs from across our library with BamA from susceptible strains using AlphaFold3^32^. Modelling the LlpA/B-BamA interaction is at the limit of AlphaFold3’s capability, and credible models were produced only in some cases. We therefore used the experimentally determined BAM_64825_-LlpA-C16 structure as a structure-guided filter, retaining the top-ranked prediction for each pair when it independently recapitulated the experimental loop 6-binding geometry. Model confidence scores (ipTM, pTM and PAE) were treated as descriptive indicators of model quality. The experimental structure was not part of the AlphaFold3 training set, so independent recovery of this geometry supports its recurrence across divergent family members but does not exclude additional binding modes for other Llp-BamA pairs. This yielded credible complexes for 17 diverse Llps, comprising seven short, three long and six broad loop 6 targeters together with LlpB-A26, which falls below the activity threshold of the colony screen, in complex with BamA from 10 *Pseudomonas* strains (Supplementary Data 11). In 12 of these models the C-terminal peptide was deployed into the BamA lumen and paired with β-strand 1, consistent with conservation of this mechanism across diverse Pseudomonas Llps. The accepted models spanned divergent LlpA and LlpB lineages and consistently recapitulated loop 6 engagement through the same N-terminal lectin-domain face observed experimentally, despite extensive sequence diversity in this region (Fig. 4a–d, Fig. S11a,b). As anticipated from sequence analysis, the single-β-lectin LlpBs bind loop 6 in a geometry analogous to the LlpAs (Fig. 4b, Fig. S11b).

**Figure 4.**
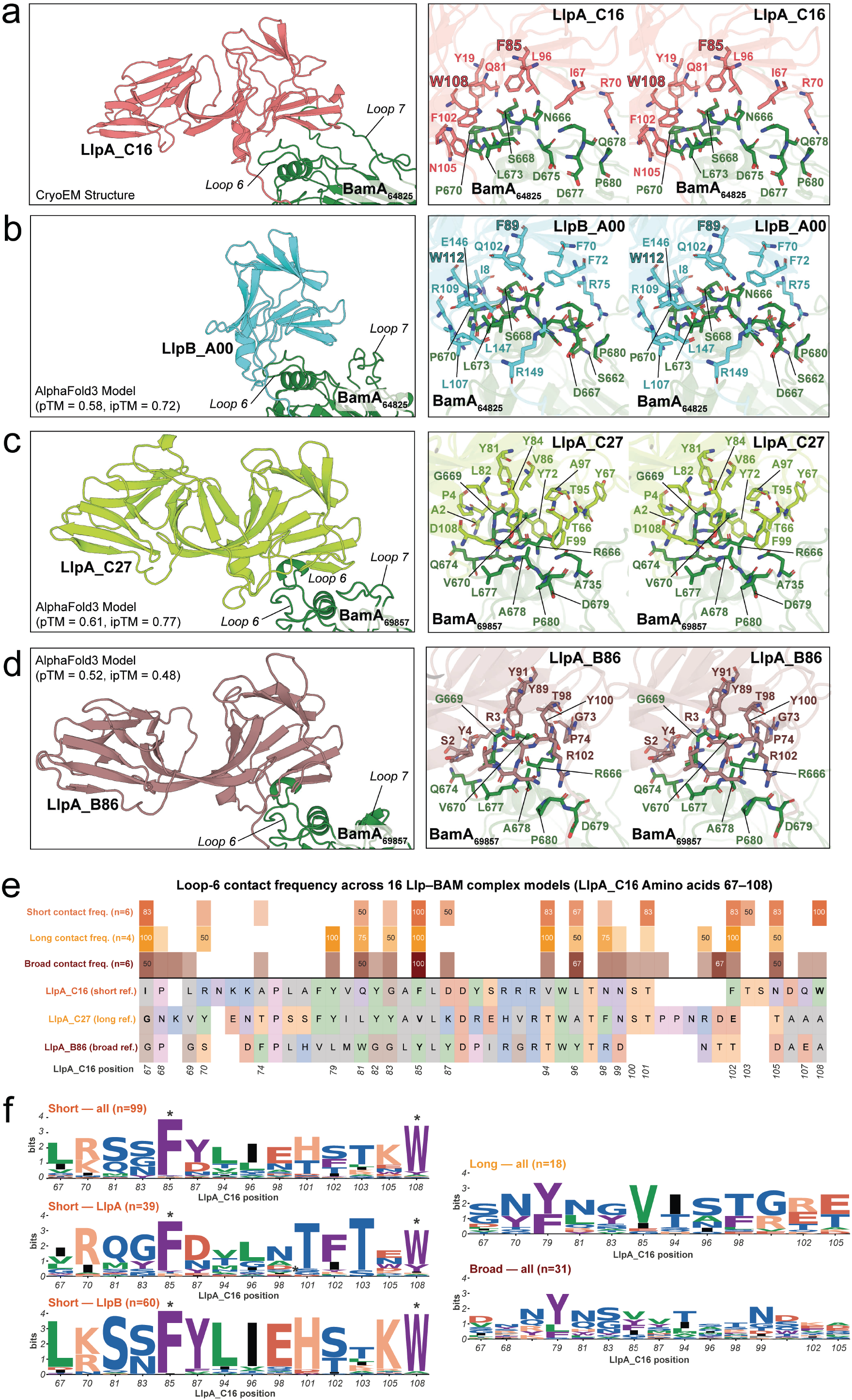
AlphaFold3 modelling indicates that diverse LlpA/Bs bind BamA loop 6 via the same N-terminal β-domain interface. (a) A cartoon view of the cryo-EM structure of BAM_64825_-LlpA-C16 showing LlpA-C16 positioning relative to BamA loop 6 (left), and right, a cross-eye stereo view of LlpA-C16 loop 6 interactions, with interacting residues shown as sticks. (b,c,d) AlphaFold3 models of LlpB_A00 (short loop 6 targeting), LlpA_C27 (long loop 6 targeting), and LlpA_B86 (broad loop 6 targeting, with long loop 6 BamA) in complex with BamA from susceptible *Pseudomonas* isolates, shown in the orientation and with the same details as BAM_64825_-LlpA-C16 in panel a. (e) A sequence-based contact map showing LlpA/B N-terminal β-lectin domain interactions with BamA loop 6 across the BAM_64825_-LlpA-C16 cryo-EM structure and 17 BamA-LlpA/B AlphaFold3 models. LlpA/Bs are divided into BamA loop 6 targeting type, with a heat map showing the frequency at which a model of that type uses a residue to form a contact with loop 6 (top), and an alignment of the sequence of the interacting region for a representative of each loop targeting type (bottom). Amino acid positions corresponding to LlpA_C16 positions are shown for reference. See Supplementary Data 11 for BamA-LlpA/B models and quality metrics. (f) Sequence logos of the conservation of the LlpA/B N-terminal β-lectin domain amino acids that make key contacts with BamA loop 6, for each of the three loop targeting types. Calculated from the sequence alignment of all LlpA/Bs in the screening panel. A key interacting residue is defined as one that contacts BamA loop 6 in >25% of the 18 structural models analysed.

For each modelled LlpA/B, we determined the amino acids from the N-terminal β-lectin domain that interact with BamA loop 6. We numbered these amino acids with the corresponding position in LlpA-C16 based on our Llp sequence alignment, and grouped the models, based on the LlpA/B BamA loop 6-targeting class (short = seven; long = three; broad = six) (Supplementary Data 1, Fig. 4e). To define the conserved interaction footprint for each loop 6-binding class, we retained contacting positions present in more than 25% of models. We then analysed conservation at these positions across these loop 6-targeting classes (short, 99 sequences; long, 19; broad, 30) in our LlpA/B library, to provide insight into the molecular basis for BamA loop-6 targeting (Fig. 4f, Supplementary Data 12).

The majority of short loop 6-targeting Llps share conserved anchoring residues at position F85 and W108 (Fig. 4a,b,e,f). These amino acids are nearly universally conserved across the short-targeting Llps and are by far the most conserved in the contact set. Separating short-targeting LlpA and LlpB sequences revealed additional class-specific conservation, with LlpAs containing a largely conserved arginine at position 70 and a TFT motif spanning positions 101 to 103. The loop 6 interface of the short-targeting LlpBs is more conserved overall than that of LlpA and diverges from it outside the F85 and W108 core. Short loop 6-targeting therefore rests on a binding core of two aromatics providing shape and non-polar complementarity, supported by peripheral contacts that vary in identity and number across the family and may contribute additional specificity or targeting breadth (Fig. 4a,b, Fig. S11b). Detailed sequence analysis indicates that F85/W108 anchoring is not the only short loop 6-binding solution. One divergent short targeting group retains the phenylalanine at position 85 but varies the second aromatic, replacing the tryptophan at position 108 and adding a tyrosine at position 105. These sequences are confined to a single clade of LlpA group 1 that is sister to a lineage of long and broad targeters, suggesting an independently derived solution to short loop recognition (Supplementary Data 12). In addition, a minority of short loop 6 targeters lack a recognisable aromatic anchor. While none of these LlpA/Bs was resolved in complex with BamA in our models, they likely represent a divergent molecular solution to short loop 6-binding (Supplementary Data 12).

The expanded loop 6 of the long loop 6 BamA variants introduces a helical turn that disrupts the surface profile recognised by the short targeters and explains why F85/W108-anchored Llps are poor targeters of long loop 6 types (Fig. 1b, Fig. S4, Fig. 4c). To accommodate this insertion, modelling indicates that long BamA loop 6-targeting LlpA/Bs have shaped their loop 6-binding face into a narrower, more concave surface that accommodates the extension, producing a deeper interface that grasps rather than caps the loop (Fig. 4c, Fig. S11a). Long loop 6 targeters engage loop 6 through a comparable number of residues to the short targeters (average of fifteen in both classes), but these positions are less conserved across the long-targeting family (Fig. 4e). The principal signatures are an aromatic residue at position 79 and a valine at position 85, both contacting the loop 6 extension directly (Fig. 4c,e). The greater binding interface divergence in this class is consistent with long loop 6-targeting LlpA/Bs having specialised towards specific loop 6 extension sequences. Separating the long targeters by loop 6 preference showed that most Llps preferring loop 6 group 3 belong to a single lineage within LlpA group 1 and share a highly conserved binding interface (Fig. S11c). One distantly related LlpB, LlpB-C14, shares this preference despite a highly divergent interface sequence, indicating that group 3 specificity has been selected for more than once (Supplementary Data 12).

In our structural models, broad-targeting Llps have the most diverse and least conserved loop 6-binding interfaces of the three classes, drawing on a larger and more variable set of contacting residues (24 positions, with relatively few contacts shared across models) (Fig. 4e,f). Although broad targeters are active against strains containing short loop BamA variants, they lack the conserved F85/W108 anchor, and their predicted interfaces differ substantially from those of short targeters in both sequence and the positions that contact loop 6, consistent with the evolution of a flexible interaction surface capable of recognising both long and short loop types (Fig. 4d,f, Supplementary Data 12). A hydrophobic residue at position 79 is the only consistently conserved position among the 30 broad targeters, with an aromatic residue at this position also a defining signature of long targeters. In broad loop 6 targeter models, position 79 contacts loop 6 in only a subset of complexes, suggesting that it shapes the binding surface rather than directly determining target recognition (Fig. 4e,f). These data suggest broad loop 6 targeters adopt a generalist strategy where multiple, relatively tolerant binding interfaces permit recognition of both loop types, contrasting with the more conserved and specialised solutions adopted by long-and short-targeting Llps.

### The *Pseudomonas* Llps inhibit the BAM complex via β-signal mimicry

In 12 of the credibly modelled AlphaFold3 Llp-BamA complexes, one for each Llp whose C-terminal peptide was modelled paired with β-strand 1, the C-terminal Llp peptide is deployed into the BamA lumen, where it binds β-strand 1 by β-augmentation (Supplementary Data 11). This mode of binding matches the C-terminal peptides of LlpA-C16 and Pyocin L2 in our cryo-EM structures, supporting the accuracy of this region of the models and providing a structurally diverse set of Llp C-terminal peptide-BamA interactions for analysis. Although the Llp C-terminal sequences of these models are highly divergent, they all bind BamA β-strand 1 using a conserved core architecture. Every C-terminal peptide β-pairs with BamA residues I425 and A427, extending to engage β-strand 1 across a region as broad as residues 422 to 433 depending on the peptide (Fig. 5a-d, Fig. S12a-d, Supplementary Data 11).

**Figure 5.**
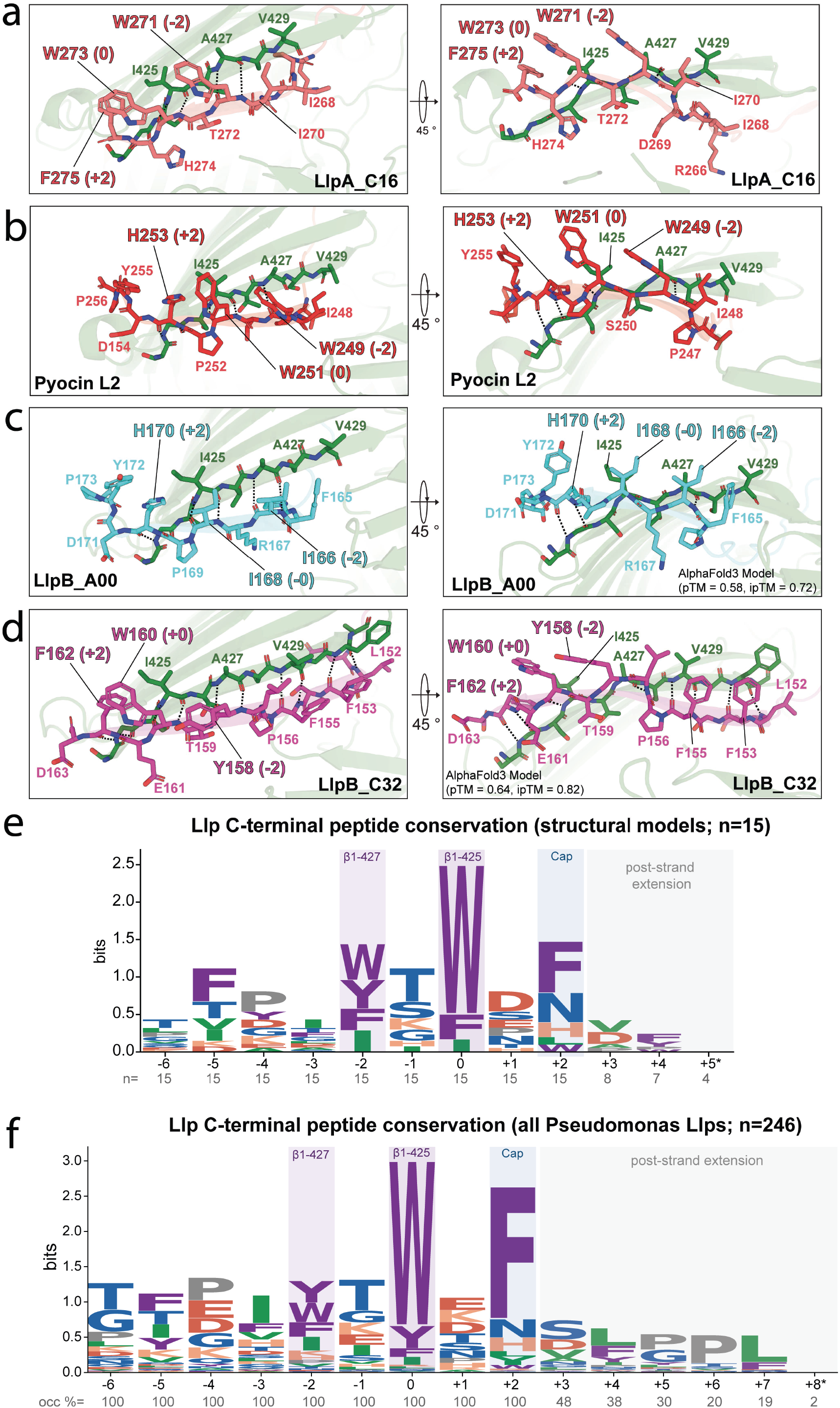
Diverse Llp C-terminal peptides inhibit BamA through conserved β-signal mimicry. (a,b) A zoomed view of the cryo-EM structures of BAM_64825_-LlpA-C16 and BAM_P28_-Pyocin L2 showing Llp C-terminal peptide interactions with BamA β-strand 1. (c,d) A zoomed view of AlphaFold3 models of LlpB_A00, and LlpB_C32 interactions with BamA β-strand 1, shown in the orientation and with the same details as BAM_64825_-LlpA-C16 in panel a. The core BamA β-strand 1 G423, I425 and A427 residues bound by Llp C-terminal peptides are labelled, as are interacting Llp C-terminal peptide residues. (e) A sequence logo based on a sequence alignment of Llp C-terminal peptides from BAM_64825_-LlpA-C16, Pyocin L2, and 12 AlphaFold3 predictions with modelled C-terminal peptide-BamA β-strand 1 interactions. (f) A sequence logo based on a sequence alignment of all C-terminal peptides from LlpA/B and L-type pyocins previously published and in our screening panel. Sequence logos from panels e and f were anchored on the amino acid aligned at position 0, with no gaps tolerated.

Consistent with the intra-membrane location of β-strand 1, the interacting regions of the Llp C-terminal peptides are amphipathic, presenting a core pair of aromatic or hydrophobic residues, designated positions 0 and -2, separated by a charged or polar residue that contacts BamA I425 and A427. Tryptophan is highly conserved in position 0 in the modelled sequences, while aromatic residues dominate at position -2 (Fig. 5e). Despite the high level of conservation, Llps like LlpB-A00 use isoleucine instead of aromatic residues at these positions, indicating that alternative bulky hydrophobic residues are viable substitutes (Fig. 5c). In addition to this conserved hydrophobic dyad, the Llp C-terminal peptide contains a cap residue, either aromatic or histidine, that reaches across BamA β-strand 1 at position G423 (Fig. 5a-e, Fig. S12a-d). This residue usually occupies the +2-position following the core aromatic/hydrophobic residue pair, but in some cases, it falls further along the strand (Fig. S12a,c).

Aligning Llp C-terminal peptides across our collection based on the conserved hydrophobic dyad indicates that the architecture of the C-terminal peptide is conserved across *Pseudomonas* Llps. The majority of Llp C-terminal peptides contain a core [Ω/Φ]-x-[Ω/Φ]-x-[Ω/H] motif that binds BamA β-strand 1 at A427, I425 and G423 (Fig. 5e,f, Supplementary Data 13). Given that BamA β-strand 1 is highly conserved across *Pseudomonas* spp., there is a surprising diversity in Llp C-terminal sequences, with 152 distinct C-terminal peptides among the 246 LlpA/B and L-type pyocin C-terminal sequences analysed. We divided the peptides into 17 sequence groups, plus a set of unassigned singletons and pairs, based on shared features (Fig. S12e, Supplementary Data 13). C-terminal peptide sequences appear to be largely inherited vertically, with similar sequence features also evolving convergently across distant lineages, showing that the family has repeatedly and independently explored the sequence space for BamA inhibition (Fig. S12e).

The architecture of the Llp C-terminal peptide mimics the β-signal, the consensus C-terminal motif (ζ-x-G-x-x-[Ω/Φ]-x-[Ω/Φ]) through which OMP substrates engage BamA β-strand 1 during assembly^34–37^. Within this motif, the [Ω/Φ]-x-[Ω/Φ] dyad forms a central element of substrate recognition. What the Llp peptides reproduce is this dyad and the augmentation register, not the consensus itself. The glycine of the β-signal is present at the equivalent register position in only 6 of the 246 Llp peptides, fewer than expected from their amino-acid composition alone, and unlike a substrate β-signal, the dyad aromatic is never the C-terminal residue, with a further two residues carrying the cap in the majority of peptides (Supplementary Data 13). The same dyad is present in an internal OMP assembly signal on the −5 strand (Φ-x-x-x-x-x-[Ω/Φ]-x-[Ω/Φ]), which is recognised by BamD, demonstrating the repeated use of this sequence feature during substrate recognition by the BAM complex^36^. The β-signal-mimetic antibiotic darobactin similarly exploits this dyad and β-strand geometry to bind BamA β-strand 1^7,8^. Llp C-terminal peptides thus represent a distinct, convergently evolved class of β-signal mimics. Rather than acting as canonical substrate signals, these hypervariable peptides deploy conserved substrate-like recognition chemistry to occupy BamA β-strand 1 and competitively block β-strand 1.

### A register-shifted state suggests stepwise deployment of the LlpA-C16 C-terminal peptide

During processing of the BAM_64825_-LlpA-C16 cryo-EM data, we identified a minor population of larger particles that reconstructed into a dimer of BAM complexes (Fig. 6a,b). The two complexes pack in a symmetrical, non-physiological head-to-tail arrangement, bridged by BamD, BamA POTRA1/2, and LlpA-C16 (Fig. 6b,c). As in the monomeric complex, the LlpA-C16 C-terminal peptide is deployed into the BamA lumen and β-augments strand 1 (Fig. 6d). Its register, however, is displaced by one step (two residues), toward the extracellular surface, with the amphipathic W271/W273 dyad and the F275 cap pairing with BamA V429, A427 and I425, rather than A427, I425 and G423 as in the fully augmented monomer, the Pyocin L2 structure, and the AlphaFold3-modelled complexes (Fig. 6e). This slippage is possible because the peptide grips β-strand 1 through side-chain-independent hydrogen bonds, a mode of binding that fixes the strand pairing but not the register, leaving the peptide free to ratchet along β-strand 1. In the dimer, turn 7 of BamA rests against the base of β-strand 1 and occludes the deeper register the peptide reaches in the monomer, apparently trapped there by the conformational constraint of dimerisation. This structure may represent an on-pathway intermediate, the inhibitory peptide caught one register step short of full deployment, with the higher register marking the earlier, less-inserted state. Strikingly, based on the length of the strand binding to BamA β-strand 1 and the position of turn 7, the Llp peptide in the dimer structure engages β-strand 1 in a way that is reminiscent of a BAM conformation during a late stage in the folding of its substrate (Fig. 6f)^6,38^. The Llp C-terminal peptide may therefore mimic not only the β-signal register of a BamA substrate, but also additional binding modes adopted by OMP substrates during insertion to achieve inhibition.

**Figure 6.**
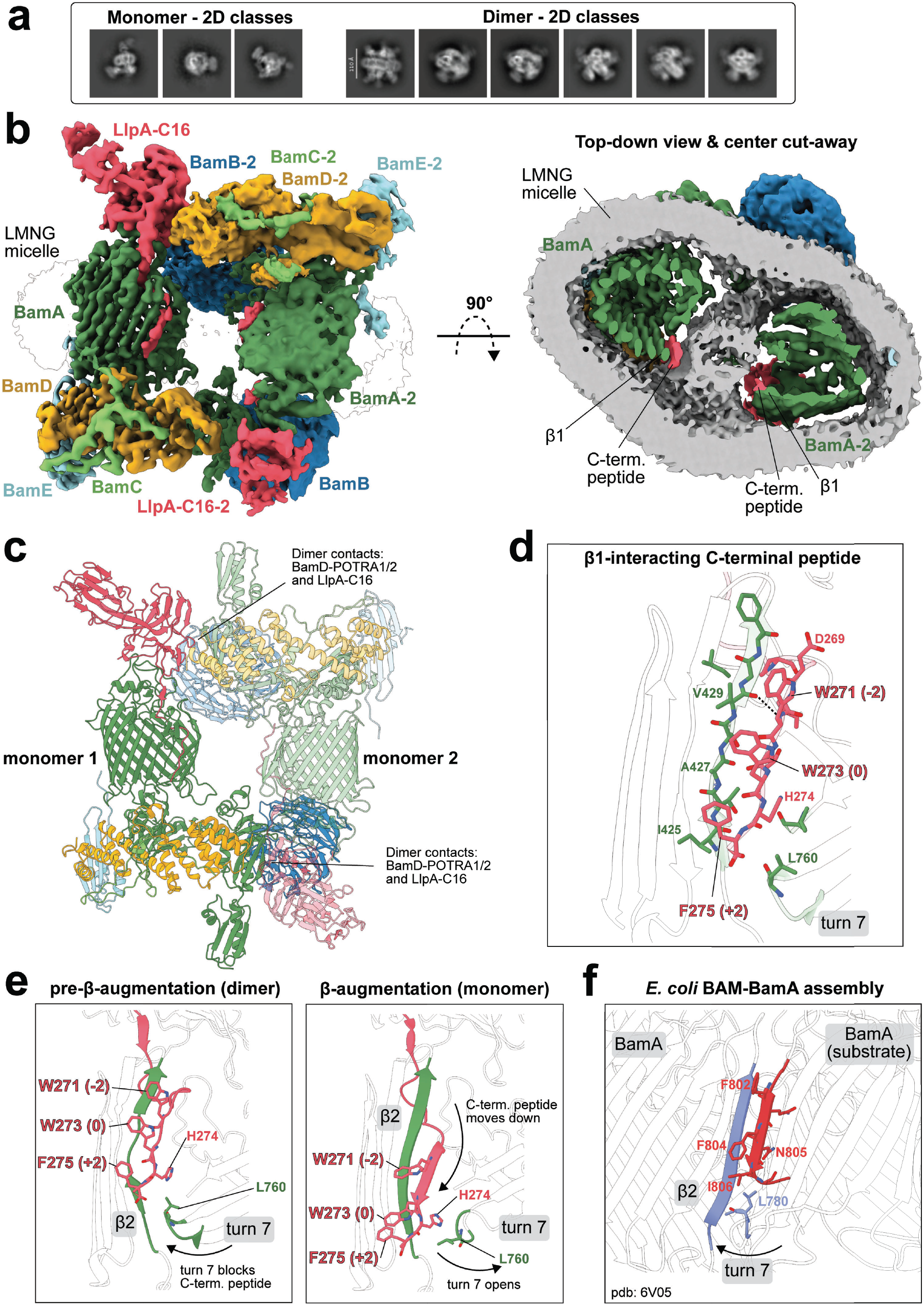
Cryo-EM of BAM-LlpA-C16 dimer subpopulation. (a) 2D classes of monomeric (left) and dimeric (right) BAM_64825_ *in complex with LlpA-C16. (b) Coulomb potential map of dimeric BAM*_64825_*-LlpA-C16 (side view) and the top-down view with centre cut-away to reveal LMNG micelle environment (grey) between BamA β-barrels. (c) Structure of dimeric BAM*_64825_*-LlpA-C16 (cartoons), with the second* protomer visualised at 50% transparency (PDB 26HE). (d) Close-up of the β-strand 1-interacting residues of the LlpA-C16 C-terminal peptide (sticks). (e) Comparison of the C-terminal peptide location in BamA from the monomeric and dimeric subpopulations. (f) A structure of the *E. coli* BamA (blue) folding a BamA substrate (red) with relevant β-strand 1-interacting residues shown as sticks, including the L780 of turn 7 (L760 in *Pseudomonas* BamA) (PDB ID = 6V05)^38^.

## Discussion

Our structural analysis of LlpA/Bs bound to BAM shows that target selection is distinct from inhibition in these protein antibiotics. A single face of the N-terminal β-lectin domain binds the variable extracellular loop 6 of BamA, while the C-terminal peptide engages the conserved β-strand 1 through main-chain β-augmentation. This division explains why loop 6 is a dominant determinant of susceptibility, why targeting preference is structured by Llp phylogeny, and how the single-domain LlpBs retain BamA-targeting capacity despite loss of the second β-lectin domain. It also distinguishes the LlpA/Bs from the L-type pyocins, which recognise the same loop using a distinct inter-domain cleft yet converge on the same inhibitory site at BamA β-strand 1^24^. *Pseudomonas* Llps have therefore diversified principally in how they recognise BamA rather than in the underlying mechanism by which they inhibit it.

Natural variation across the family shows that BamA loop 6 recognition has been solved in several distinct ways. Short-loop specialists are dominated by relatively conserved interaction signatures, including the F85/W108 aromatic anchor, whereas long-loop targeters accommodate the inserted, longer loop through deeper and more variable interfaces that themselves specialise towards different long-loop sequence families. Broad targeters represent a third solution. Their loop 6-binding surfaces are the least conserved and draw on a larger and more variable set of contacts, consistent with generalism arising from a comparatively tolerant interaction surface rather than from extension of the short loop binding mode. These targeting classes are substantially inherited within Llp lineages, but have also arisen repeatedly, indicating that BamA loop 6-recognition can be remodelled without disrupting the overall Llp architecture. The independent emergence of LlpBs provides an extreme example of this modularity. Because BamA recognition is carried entirely by the retained N-terminal β-lectin domain, loss of the C-terminal lectin domain can preserve target recognition while removing a module implicated in O-antigen binding. How LlpBs compensate for this loss during initial engagement with the bacterial surface, and how domain loss affects antibacterial potency, remain important unresolved questions.

The targeting and inhibitory modules are subject to strikingly different sequence constraints. BamA loop 6 is hypervariable, and the Llp surfaces that recognise it vary accordingly. By contrast, the β-strand 1 inhibitory site is effectively invariant across the Pseudomonas isolates examined here (mean per-position identity 99.8% across residues 419-433 of the 111 isolate BamA sequences, with G423, I425 and A427, the three residues β-paired by the Llp C-terminal peptides, invariant in every isolate), yet the peptides that engage it are the most sequence-diverse region of the Llps. Structural comparison resolves this apparent paradox. β-augmentation is dominated by main-chain hydrogen bonding, imposing relatively little side-chain specificity beyond an amphipathic register, a conserved hydrophobic or aromatic anchor, and a cap residue. The result is a family containing over 150 distinct C-terminal peptides that nevertheless reproduce the same interaction chemistry. Because BamA must accept the C-terminal strands of hundreds of different outer-membrane proteins at the same lateral gate, this permissiveness is likely intrinsic to the machine. The diversity of Llp C-terminal peptides is therefore likely an expected consequence of β-signal mimicry. This architecture parallels the independently evolved β-signal mimicry of darobactin^7^. BAM substrate recognition therefore represents a recurrent vulnerability, where unrelated antibacterials have converged on the chemistry BamA normally uses to recognise and fold its own substrates.

The register-shifted LlpA-C16 C-terminal peptide structure provides further insight into how this mimicry may operate. In the dimeric cryo-EM population, the inhibitory peptide remains β-paired to BamA β-strand 1 but is displaced towards the extracellular surface by one register position relative to the fully deployed state. Because the interaction is mediated principally through the peptide backbone, β-pairing can be maintained while the peptide changes register, providing a plausible mechanism by which the peptide could progressively advance along β-strand 1, towards its final inhibitory state. Although this state was captured in a non-physiological BAM dimer and should therefore be interpreted cautiously, its correspondence with the position occupied by an OMP β-signal during folding is notable^38^. Rather than simply mimicking the final bound state of a substrate, Llps may exploit the additional conformational states used by BamA to template substrate folding. In this model, substrate-like engagement becomes a kinetic route to inhibition where the Llp peptide enters the normal folding pathway but terminates in a stable, non-productive complex. This model raises the possibility that initial peptide engagement is favoured by an outward-open BamA conformation.

The Llp modular architecture is particularly attractive as a starting point for antibacterial engineering. The family provides diverse specialist and generalist targeting surfaces coupled to a conserved inhibitory mechanism. Altering the loop 6-binding face should in principle permit the spectrum to be redirected without redesigning the inhibitory peptide, while the sequence tolerance of the C-terminal peptide provides latitude to optimise deployment, stability or potency without changing its fundamental binding chemistry. More broadly, Llps illustrate a strategy that is relevant to advanced protein design. A variable protein-recognition module can be coupled to a conformationally responsive effector that acts through substrate mimicry at a conserved functional site. Current de novo binder design is particularly effective at constructing high-affinity interactions with defined protein surfaces^39–44^. The natural Llp family shows how such recognition can instead be integrated with sequential molecular interactions to produce a specific mechanistic outcome. Understanding this process not only provides principles for extracellular BAM inhibition but may also inform the development of protein-based molecular machines.

Several remaining questions will determine how general and useful this principle will be. Can the Llp spectrum be redirected predictably by exchanging or redesigning loop 6-recognition surfaces? Can the conserved inhibitory peptide be coupled to BamA binders targeting other Gram-negative genera, thereby extending β-signal mimicry beyond *Pseudomonas*? And how readily can pathogenic strains escape these molecules while retaining functional BamA and outer-membrane biogenesis? Targeting an invariant inhibitory site while encoding specificity in a separable surface-recognition module may facilitate the evolution of resistance, but the fitness costs and alternative mechanisms of escape remain to be established experimentally. Resolving these questions will determine whether Llps remain specialised products of competition between pseudomonads or provide a general framework for engineering precision antibacterials against Gram-negative bacteria.

## Methods

### Bacterial strains and culturing conditions

*Escherichia coli* DH5α was made chemically competent for cloning and grown in Lysogeny Broth (LB) (10 g/l Tryptone, 5 g/l Yeast extract, 5 g/l NaCl, 15 g/l agar) with appropriate antibiotics to maintain selection. *E. coli* cultures were grown at 37°C and 200 rpm or otherwise specified in the protein purification section. *Pseudomonas* strains were grown at 30°C on King’s B (KB) agar (20 g/L proteose peptone, 10 mL/L glycerol, 1.5 g/L K2HPO4, 1.5 g/L MgSO4 and 15 g/L agar) or in KB broth at 30°C and 180 rpm.

### Pseudomonas Llp identification

A representative set of putative Llp proteins from *Pseudomonas* was identified by searching the NCBI non-redundant (NR) protein database using BLASTP with the amino acid sequences of putidacin L1, LlpA_Pf-5_ pyocin L1 and pyocin L2, as the query, with no E-value cutoff applied. Identical or near-identical sequences were removed from the dataset using CD-HIT with a sequence identity cutoff of 98%. Remaining sequences were manually curated, retaining those with domain features consistent with either LlpA or LlpB proteins (one or two β-lectin (MMBL) domains, together with the C-terminal peptide and linker regions). This yielded a final, dereplicated set of 238 *Pseudomonas* Llp sequences (106 LlpA and 132 LlpB; Supplementary Data 1), which together with characterised L-type pyocin/putidacin sequences and *Burkholderia* and *Xanthomonas* Llp sequences comprise the 268-sequence set used for phylogenetic analysis. Codon-optimised genes encoding these 238 Llps (including 4 L-type pyocins, later identified as such) for expression in *E. coli* were synthesised by Twist Bioscience and cloned into pET29b with no affinity tag. These untagged Llp constructs were used for the colony-based Llp activity assays.

### Statistical Analysis

All statistical analyses were performed in Python 3 using NumPy, pandas, SciPy and scikit-learn, and unless stated otherwise multiple-testing correction used the Benjamini–Hochberg false discovery rate (FDR), with significance assessed at FDR < 0.05. To classify BamA loop 6 preference, the binary Llp × strain susceptibility matrix (238 Llps × 101 strains; 23,406 scored Llp–strain observations; Supplementary Data 4; 632 of the 24,038 possible Llp-strain combinations were not scored and were excluded pairwise) was first filtered to remove low-activity Llps, which are expected to contain a high proportion of false-positive calls: Llps that inhibited five or fewer strains across the panel were excluded, rejecting 90 Llps and retaining 148 confidently active Llps for categorisation. For each retained Llp, the odds ratio of long versus short loop 6 hits was computed with a Haldane-Anscombe 0.5 continuity correction to accommodate zero cells, using only the 98 strains with an assigned short (n = 52) or long (n = 46) loop 6 type; the three outgroup strains were excluded from this calculation. Llps with less than a two-fold skew (|log2 odds ratio| < 1) were classified as broad targeting, while those with a ≥2-fold skew were classified as short or long loop 6-targeting accordingly, unless the disfavoured loop 6 class was engaged at ≥3 hits and ≥10% of that class, in which case the Llp was reclassified as broad targeting. This engagement criterion prevented Llps with genuine dual loop activity from being misclassified as single loop specialists on the strength of a modest fold-skew alone (e.g. LlpA_C20, which killed 5/45 long loop and 14/53 short loop strains, a >2-fold skew towards short, but a clearly dual-active profile). Association between Llp activity and BamA loop 6 identity was tested in three complementary ways on the curated long-format matrix (Supplementary Data 5): per-Llp Fisher exact tests of each Llp against each loop 6 group (3,552 tests; 60 significant at FDR < 0.05), per-Llp chi-square tests of independence across all loop 6 groups (238 tests; 73 significant), and per-Llp long-versus-short 2 × 2 contrasts (238 tests; 84 significant), with P values corrected within each test family. To separate the contributions of the two adjacent hypervariable loops, pairwise Jaccard distances between strain susceptibility profiles (over the full 238-Llp panel, across all 101 strains) were correlated with pairwise sequence distances for loop 6 and for loop 7 using Mantel tests. Loop-sequence distance was defined as the fraction of differing positions across all alignment columns, and distance vectors were correlated using Spearman rank correlation, with significance assessed by random permutation of the distance matrices; because the loop 6 and loop 7 distance matrices are themselves strongly correlated, partial Mantel tests were used to estimate the correlation of each loop with the susceptibility profile while controlling for the other (Table S1).

Agreement between the three independent replicates of the purified-Llp overlay assay (21 Llps × 10 strains = 210 Llp–strain cells per replicate, scored 0–5) was quantified with the intraclass correlation coefficient for absolute agreement, ICC(2,1), treating replicates as raters and Llp–strain cells as subjects (ICC = 0.91), with Kendall’s coefficient of concordance across the three replicates (W = 0.61, computed without a tie correction) and, after binarising each cell as active/inactive, with Fleiss’ kappa (κ = 0.69); binary calls were unanimous across the three replicates in 166/210 (79%) cells, and a per-cell consensus score was taken as the median of the three replicates for all downstream comparisons. Concordance between the two assays was assessed over the Llp–strain pairs scored in both, comparing the binarised purified-protein consensus score (active if consensus ≥ 1) with the binary colony-screen call using percentage agreement and Cohen’s kappa, and comparing the graded consensus potency with the binary screen call using the area under the receiver operating characteristic curve (ROC-AUC) and Spearman’s rank correlation.

Loop 6 preference was compared between assays using PI = (long − short)/(long + short), where long and short denote the mean consensus inhibition scores for the corresponding strain classes in the purified-protein assay, and the corresponding fractions of strains inhibited in the colony screen. PI ranges from −1 (exclusive short-loop activity) through 0 (no preference) to +1 (exclusive long-loop activity). The three purified Llps below the colony-screen activity threshold (fewer than six strains inhibited) were flagged because their colony-screen PI estimates were less reliable. PI concordance was assessed across the seventeen Llps active in the colony screen; agreement in the direction of preference was also assessed for the twelve Llps with |PI| ≥ 0.3 in both assays (Fig. 2b; Supplementary Data 10).

### Solid media overlay growth inhibition assays

For inhibition assays with purified lectin-like bacteriocins, Pseudomonas strains (Table S5) were inoculated from glycerol stocks or fresh colonies and grown in KB medium at 30°C and 180 rpm. After reaching maximum optical density, 100 µL of culture was mixed with 6 mL of fresh 0.6% agar dissolved in milliQ water, cooled until it was warm but remained liquid, and overlaid on the KB agar plate. The concentration of purified lectin-like bacteriocins was adjusted to 0.5 mg/mL, after which 1 µL was spotted on top of the solidified agar plate. After the spots dried in sterile conditions (∼5 minutes), the plates were transferred to 30°C and grown for ∼16 hours for inhibition profile readout by eye.

### Lysis-based overlay growth inhibition assays

Each of the 238 untagged Llp expression constructs (pET29b) was transformed into *E. coli* BL21(DE3)C41 and arrayed as a discrete colony on nutrient agar with or without 0.1 mM IPTG in a 96-colony microplate format across three plates (Supplementary Data 3). Arrays were replicated with a pinning tool and grown at 30°C for 24 hours to establish colonies of even size. Colonies were lysed *in situ* by exposure to chloroform vapour for 60 seconds, followed by exposure to air for 600 seconds to allow residual solvent to evaporate. Lysed arrays were overlaid with 6 mL of 0.6% (w/v) soft agar seeded with 100 µL of a stationary-phase culture of a single *Pseudomonas* isolate from the panel, and incubated at 30°C for 16–24 h to establish confluent growth of the indicator strain. A clear zone of growth inhibition surrounding a colony was recorded as a positive activity call and the absence of a zone as negative, giving a binary Llp × strain susceptibility matrix (Fig. S2, Supplementary Data 4).

### *Pseudomonas* whole genome sequencing, genome assembly and phylogeny

Genomic DNA was extracted from overnight cultures of each *Pseudomonas* isolate using Monarch Spin gDNA Extraction Kit (NEB), and paired-end libraries were prepared and sequenced on an Illumina platform at GeneWiz, generating 2 × 150 bp reads (median 5.75 M read pairs per isolate; range 3.7–12.4 M). Read processing, assembly and annotation were performed with the ProkGenomics pipeline v1.0.0 (https://github.com/Grinter-Lab/ProkGenomics): read quality was assessed with FastQC before and after adapter and quality trimming with Trimmomatic; trimmed reads were assembled *de novo* with Unicycler; assembly completeness and contamination were assessed with CheckM; contigs were classified as chromosomal, plasmid or phage using PlasClass and CheckV; coding sequences were annotated with Prokka (bacterial contigs) and Pharokka (phage contigs); and taxonomy was assigned with GTDB-Tk using the bac120 marker set against the GTDB reference database. Assemblies passing quality control (CheckM completeness ≥ 98% and contamination ≤ 5%) were retained, giving 111 *Pseudomonas* isolate genomes (median completeness 100%, range 99.3–100%; median contamination 0.22%, range 0.05–2.11%) with a median size of 6.06 Mb (range 5.05–7.37 Mb), a median of 105 contigs (range 33–538) and a median contig N50 of 262 kb (range 49–1,489 kb). GTDB-Tk assigned all 111 isolates to the family Pseudomonadaceae: 108 to *Pseudomonas_E*, two to *Pseudomonas* (both *P. aeruginosa*) and one to *Pseudomonas_B*, spanning 35 named species (Supplementary Data 2); assemblies and raw reads are deposited under BioProject <u>PRJNA1530708</u>. To place the isolate panel in a genome-based phylogeny, GTDB-Tk (*align* step) was used to extract the 120 bacterial single-copy marker genes (bac120), align each to its reference HMM and concatenate the per-marker alignments into a single trimmed amino-acid supermatrix after GTDB-defined column masking (5,035 amino-acid sites; 71.4% constant sites, 1,120 parsimony-informative sites). A maximum-likelihood tree was inferred from this supermatrix with IQ-TREE v2.3.1, with the substitution model selected automatically by ModelFinder, which chose Q.insect+F+I+R3 by BIC (empirical amino-acid frequencies, a proportion of invariable sites and a three-category FreeRate model of among-site rate heterogeneity). Branch support was assessed with 1,000 ultrafast bootstrap (UFBoot) replicates together with the SH-aLRT test (random seed 24195), and nodes are annotated as SH-aLRT (%) / UFBoot (%). The final tree contains 110 taxa (one isolate, *P. amygdali* 65893, was not placed) and was visualised as a circular cladogram with tips labelled by GTDB species assignment and coloured by BamA loop 6 type (Fig. S5).

### Cloning of expression constructs

For BAM constructs, gene fragments were synthesised by Twist Bioscience and subsequently cloned into the L-rhamnose-inducible expression vector pScRhaB2 using seamless ligation cloning extract (SLiCE) and conventional restriction cloning^45^. Each construct encodes the *bamABCDE* operon, with a Twin-Strep-tag II fused to the C-terminus of BamA for affinity purification and the remaining subunits untagged. Lectin-like bacteriocin genes were synthesised by Twist Bioscience and cloned into pET29b, either untagged (for the colony-based screen) or with an N-terminal 6×His tag (for protein purification). Point mutants were generated with the Q5 Site-Directed Mutagenesis Kit (NEB) according to the manufacturer’s instructions, and all constructs were verified by whole-plasmid sequencing. Lists of all constructs and primers used in this study are available (Tables S3 and S4).

### BAM expression and purification

BAM expression and purification was performed as described in previous studies, with minor modifications^24^. For the overexpression of BAM_64825_ and BAM_82441_, chemically competent *E. coli* BL21(DE3)C41 cells (Table S5) were transformed with the appropriate plasmids (Table S3) and plated on LB agar with 100 µg/mL Trimethoprim to grow overnight at 37°C. The next day, multiple colonies were used to inoculate starter cultures in TB medium for growth overnight at 37°C and 200 rpm. Sterile TB medium was inoculated with starter cultures at an approximately 1:40 ratio and grown at 37°C and 200 rpm until an OD_600_ ∼0.9-1. The cultures were chilled at 4°C for 30 minutes before induction of BAM expression with 0.2% L-Rhamnose. Expression was performed for 20-22 hours at 25°C and 200 rpm. Usually, 8 L of cultures yielded ∼100-140 g cell pellets, which could be stored at -80°C for several weeks.

Cell pellets were partially thawed with lukewarm water for ∼5 minutes and resuspended in lysis buffer (20 mM Tris-HCl pH 8, 150 mM NaCl, 2 mM MgCl_2_, 0.1 mg/mL lysozyme, 0.05 mg/mL DNase, 1x EDTA-free protease inhibitor). Cell lysis was performed with a high-pressure cell disruptor, requiring 2-3 passes with cooling on ice in between each pass. The lysate was centrifuged at 29,000 *× g* for 20 minutes at 4°C. The supernatant was then centrifuged at 100,000 *× g* for 1 hour at 4°C, yielding brownish-red membrane pellets. The membranes were resuspended in solubilisation buffer (20 mM Tris-HCl pH 8, 150 mM NaCl, 1% N-dodecyl-β-D-maltoside (DDM), 300 µL biotin blocking buffer (BioLock, 2-0205-050)), homogenised with a glass homogeniser, and incubated at 20°C for 30 minutes with gentle orbital shaking. Then, solubilised membranes were centrifuged at 30,000 *× g* for 40 minutes at 20°C to pellet any cell debris. From here, the supernatant was kept on ice and loaded twice on a Strep-Tactin XT 1-mL column (Cytiva) using an ÄKTA Start purification system (Cytiva). Afterwards, DDM detergent was exchanged with lauryl maltose neopentyl glycol (LMNG) detergent by applying 40 column volumes (CV) of ice-cold detergent-exchange buffer (20 mM Tris-HCl pH 8, 150 mM NaCl, 0.025% lauryl maltose neopentyl glycol (LMNG)). Bound protein was eluted by hand with a syringe containing 10 mL elution buffer (20 mM Tris-HCl pH 8, 150 mM NaCl, 0.025% LMNG, 75 mM biotin) and collected in 1 mL fractions. After 3 mL elution, the column was incubated in elution buffer for 5 min on ice to improve protein elution. The fractions were analysed by SDS-PAGE with Coomassie staining. Clean fractions were pooled and concentrated in a 15-mL concentrator (Amicon Ultra Centrifugal Filter, 30 kDa MWCO) to a final volume of 500 µL for injection on a Superose 6 10/300 Increase column for size exclusion chromatography at room temperature (20 mM Tris-HCl pH 8, 150 mM NaCl, 0.01% LMNG). The BAM complex eluted according to its molecular weight at the expected retention volume, and appropriate fractions were concentrated to ∼500 µL volume in a 30 kDa cut-off concentrator (Amicon) to avoid protein loss by excessive concentration. Purified BAM could be stored in size exclusion buffer at -80°C without significant disassembly or degradation of the complex. Protein yields differed between BAM_64825_ and BAM_82441_ but ranged from approximately 20–35 µg per litre of expression culture.

### Removal of excess LMNG from purified BAM

To remove LMNG from the sample, purified BAM was thawed on ice and diluted once with 15 mL of ice-cold detergent-free buffer (20 mM Tris-HCl pH 8, 150 mM NaCl) and re-concentrated with new 15-mL concentrators and new 0.5-mL concentrators (Amicon Ultra Centrifugal Filters, 100 kDa MWCO). With this dilution technique, only minimal amounts of BAM-free LMNG micelles were observed during cryo-EM, which significantly improved sample quality.

### Lectin-like bacteriocin large scale expression and purification

For the overexpression of lectin-like bacteriocins from *Pseudomonas*, chemically competent *E. coli* BL21(DE3)C41 cells were transformed with the appropriate plasmids (Table S3) and plated on LB agar with 50 µg/mL kanamycin and incubated overnight at 37°C. The next day, multiple colonies were used to inoculate starter cultures in LB medium for growth overnight at 37°C and 200 rpm. LB medium was inoculated with starter cultures at an approximately 1:40 ratio and grown at 37°C and 200 rpm until an OD_600_ ∼1. The cultures were chilled at 4°C for 30 minutes before induction of Llp expression with 0.3 mM Isopropyl β-D-1-thiogalactopyranoside (IPTG). Expression was performed for ∼20 hours at 20°C and 200 rpm. Cell pellets were stored at -80°C for several months.

Cell pellets were partially thawed with lukewarm water for ∼5 minutes and resuspended in lysis buffer (20 mM Tris-HCl pH 8, 500 mM NaCl, 2 mM MgCl_2_, 5 mM Imidazole, 0.1 mg/mL lysozyme, 0.05 mg/mL DNase, 1x EDTA-free protease inhibitor). Cell lysis was performed with a high-pressure cell disruptor, and the lysate was centrifuged at 34,000 *× g* for 20 minutes at 4°C to remove cell debris and unbroken cells. The lysate was loaded twice on HisTrap 5-mL (Cytiva) immobilized metal affinity columns using an ÄKTA Start purification system (Cytiva). The column was transferred to an ÄKTA Pure purification system (Cytiva) and washed (20 mM Tris-HCl pH 8, 500 mM NaCl, 5 mM Imidazole) for several column volumes, the amount of which depended on the Llp sample and contamination. Elution was performed with a 5/10/25/50/100% step gradient (20 mM Tris-HCl pH 8, 500 mM NaCl, 1 M Imidazole) with Llps usually eluting at 100-500 mM Imidazole. The elution fractions were investigated via SDS-PAGE, and appropriate fractions pooled and subjected to size exclusion chromatography on a HighLoad^TM^ Superdex 75 pg column (Cytiva). The yield and solubility of N-terminally tagged Llps varied widely, ranging from 2–50 mg per litre of culture; solubility also differed between proteins. Llps could be stored at - 80°C for months, but some tended to aggregate after thawing on ice and temperature changes. For this reason, Llps were always centrifuged for 5 min at 15,000 *xg* at 4°C after thawing and before use.

### Lectin-like bacteriocin small scale expression and batch purification

First, a 10 mL 24-deep well block (Axygen, LOT: 19523070) was pre-chilled on ice, after which 20 µL of fresh chemically competent *E. coli* BL21(DE3)C41 were added to the bottom of each well. Then, 0.5 µL of Llp expression plasmids (∼200 ng/µL) were added into each well, mixed by careful shaking by hand, and incubated on ice for 3 minutes. For *E. coli* transformation, the bottom quarter of the 24-deep well block was submerged in a pre-heated 42°C water bath for 60 seconds and transferred on ice for another 5 minutes. 1 mL of LB broth was added to each well and the 24-deep well block was sealed with a pierceable sealing mat (Fisher Scientific, LOT: 08720029) for heat shock recovery for 1 hour at 37°C and 180 rpm. After recovery, the appropriate antibiotic was added to each well and the 24-deep well block was sealed again for growth overnight in the same conditions. The next day, overnight expression autoinduction TB broth (Millipore, LOT: 4309179) was prepared with the appropriate antibiotic following the manufacturer’s instructions. Into 250-mL conical flasks, 50 mL of TB broth was added, and each flask was inoculated with 500 µL of starter culture from the 24-deep well block. Cultures were grown for 16-24 hours at 30°C with shaking at 180 rpm, to allow for auto-induced Llp expression.

After expression, cells were harvested by centrifugation and the cell pellets were weighed (expected yield 0.5 - 0.6 g) and used immediately or stored at -20°C for weeks. For cell lysis, the B-PER™ Bacterial Protein Extraction reagent (Thermo Scientific, LOT: 3216730) was prepared as per the manufacturer’s instructions and half a tablet of proteinase inhibitor was added per 66 mL of B-PER solution. Then, 5 mL B-PER solution per 1 g of bacterial cell pellet was added to a 50-mL Falcon tube and the cells were thoroughly resuspended with a pipette. The cell suspension was incubated at room temperature for ∼15 minutes with gentle agitation on a rotary shaker, after which cell debris was pelleted by centrifuging for 15 minutes at 4°C and 3900 rpm. Then, the clear cell lysate (supernatant) was transferred to a fresh 24-well plate. For affinity purification, 100 µL of loose His60 Ni Superflow Resin (Takara Bio, LOT: 2407192A) in nickel binding buffer (50 mM Tris-HCl pH 7.9, 200 mM NaCl, 20 mM imidazole) was added, the plate sealed with a fresh pierceable 24-well plate mat and incubated at 4°C for 120 minutes at 180 rpm. The lysate-nickel resin mixture was transferred to a 2-mL 24-well filter plate (AcroPrep™, LOT: FL8318) and the lysate removed with a vacuum manifold (Pall Life Sciences). Then, 5 mL of nickel binding buffer was added to the wells, incubated for 3 minutes and removed via vacuum. This wash step was repeated 3 times. Afterwards, 500 µL of nickel gradient buffer (50 mM Tris-HCl pH 7.9, 200 mM NaCl, 500 mM imidazole) was added into each well and incubated at room temperature for 10 minutes. Then, the protein-buffer solution was moved from the 24-well filter plate into a clean 24-well plate by applying a vacuum. This elution step was then repeated with fresh 500 µL of nickel gradient buffer, and the second eluent was collected in a separate 24-well plate. The quality of the purified protein was assessed by SDS-PAGE.

### SDS-PAGE analysis

Protein samples were analysed using Bolt 4–12% SDS-PAGE according to the manufacturer’s instructions. Gels were stained with AcquaStain Protein Gel Stain (Bulldog) and subsequently washed in distilled H_2_O. Protein samples were generally loaded at a constant volume.

### Cryo-electron microscopy single particle imaging

All sample preparation and data collection were performed at the Ian Holmes Imaging Centre at the University of Melbourne. For grid preparation, UltrAuFoil gold grids (1.2/1.3, 300 mesh, Quantifoil GmbH) were glow discharged at 15 mA for 3 minutes in air with a PELCO easiGlow (Ted Pella Inc.). For apo BAM_82441_, 3 µL of 7 mg/mL sample was applied, and for BAM_64825_–LlpA-C16, 3 µL of a 6.5 mg/mL sample was used. The excess protein solution was blotted off using a blot force of -9 (apo BAM_82441_) and -12 (BAM_64825_-LlpA-C16), a blot time of 3 s, and a blot total of 1 with a Vitrobot Mark IV, FEI (Thermo Fisher Scientific) at 100% humidity and 4°C. Plunge-frozen grids were clipped and screened for sample quality and ice thickness using a Talos Arctica microscope with a Falcon 4 camera at 120,000× magnification (EFTEM mode) with a pixel size of 1.19 Å. Collecting between 100–400 exposures was sufficient to obtain good quality data that allowed initial 2D classification and ∼4 Å 3D reconstructions. Selected grids were stored for subsequent dataset collection on the Titan Krios G4 at 300 kV with a Falcon 4i camera at 96,000× magnification (EFTEM mode) with a pixel size of 0.808 Å in electron counting mode.

For apo BAM_82441_, 8,650 exposures were collected with three exposures per grid hole (730 nm illumination area, total dose 50.24 e^−/Å²^, and exposure time 5.62 seconds) fractionated into 192 frames (Fig. S8). All data processing was performed in cryoSPARC v4.6.2^46–49^, starting with patch motion correction and patch CTF estimation. Afterwards, the exposure groups were determined and the exposures curated, removing any exposures with an estimated CTF of >4 Å. Initially, 3,450,061 particles were extracted using the cryoSPARC blob picker with a box size of 320 pixels, binned to 80 pixels. After 2D classification and ab initio reconstruction with 5 classes (no similarity), 354,218 particles were determined to be high quality, re-extracted at 320 pixels without binning, and used to train the Topaz particle picker^50^. From here, 3,272,308 particles were extracted at 320 pixels, binned to 80 pixels, and used for 2D classification. Selected 2D classes yielded 1,167,789 particles, which were re-extracted at 320 pixels, unbinned, and used for ab initio reconstruction with 3 classes (no similarity). The best class (773,983 particles) was used for non-uniform refinement, yielding a Coulomb potential map with 2.66 Å global resolution. The same particles were used for Global CTF refinement and subjected to 2D classification, yielding 709,510 particles. Non-uniform refinement resulted in a cryo-EM map with 2.36 Å global resolution (B factor 66.7), which could not be further improved by adjusting aberration correction parameters or baited heterogeneous refinement. In ChimeraX 1.10, three masks were generated (1. BamA β-barrel-POTRA 5, 2. BamA-POTRA 2/3+BamB, 3. BamA-POTRA 2/3/4/5+BamCDE), for local refinement to account for independently flexible domains of apo BAM_82441_.

For BAM_64825_-LlpA-C16, 10,860 exposures were collected with three exposures per grid hole (750 nm illumination area, total dose 50 e^−/Å²,^ exposure time 3.95 seconds) fractionated into 135 frames (Fig. S9). All data processing was performed in cryoSPARC v4.6.2, starting with patch motion correction and patch CTF estimation. Afterwards, the exposure groups were determined and the exposures curated, removing any exposures with an estimated CTF of >4 Å. After particle picking with the cryoSPARC blob picker, 2,195,428 particles were extracted at 320 pixels, binned to 160 pixels. These were used for 2D classification, which yielded 2D classes accounting for 777,635 particles, which were used for ab initio reconstruction with 5 classes (no similarity). These particles from the best class (321,131 particles) were re-extracted at 320 pixels, unbinned, and used for ab initio reconstruction with 2 classes (high similarity). The best class with 221,280 particles was used for non-uniform refinement, which generated a Coulomb potential map with a global resolution of 2.88 Å. The same particles were used for Topaz training and particle picking, resulting in 3,079,353 particles which were extracted at 320 pixels, binned to 160 pixels. Two rounds of 2D classification and class selection yielded 873,675 particles, which were used for ab initio reconstruction with 6 classes (no similarity). 338,236 particles from the best class were re-extracted at 320 pixels, unbinned, subjected to a final 2D classification, resulting in 307,084 particles. These particles were used for a final ab initio reconstruction with 2 classes (no similarity) and used for Global CTF refinement. These 265,173 particles were used for non-uniform refinement (with parameters: EWS correction, Fit Anisotropic Magnification, and Optimize per-group CTF) and generated a cryo-EM map with 2.66 Å global resolution (B factor 65.7). To account for flexibility, two masks (1. top of BamA β-barrel+LlpA-C16, 2. BamA-POTRA 2/3+BamB) were made in ChimeraX 1.10 and used for local refinement, increasing local map quality.

For the same dataset, a significant subpopulation of BAM64825–LlpA-C16 dimers was identified during early 2D classifications (Fig. S13). The cryoSPARC blob picker was rerun, accounting for larger particle dimensions, resulting in 2,435,633 particles which were extracted at 320 pixels and binned to 80 pixels. These were used for two rounds of 2D classification and ab initio reconstruction with 3 classes (no similarity). The best class yielded 110,941 particles, which were used for ab initio reconstruction with 2 classes (no similarity), the best of which (80,696 particles) was consistent with two BAM_64825_–LlpA-C16 molecules that were rotated by 180° and enclosed by a single LMNG micelle. Multiple rounds of selecting different 2D classes, ab initio reconstruction, and non-uniform refinement resulted in a cryo-EM map clearly depicting the dimer at 6.89 Å (22,141 particles). These particles were re-extracted without binning and used for Topaz training and picking, where it was important to optimise the expected particles/micrograph. Here, 594,128 particles were extracted at 320 pixels, and binned to 160 pixels. After 2D classification, 304,957 particles were used for ab initio reconstruction with 5 classes (no similarity). Three of these classes were selected, which was confirmed by three 2D classifications with the particles of each class individually. The particles of the three ab initio classes (combined 134,979 particles) were used for non-uniform refinement, resulting in a 3.38 Å cryo-EM map. These particles were re-extracted at 320 pixels without binning and used for Global CTF Refinement, followed by non-uniform refinement. This generated a cryo-EM map with a global resolution of 2.95 Å (B factor 69), with one of the monomers having significantly higher resolution than the other. In ChimeraX 1.10, masks of each monomer and both monomers combined (without the LMNG micelle) were created for local refinement, further improving resolution for each monomer.

### Visualization of cryo-EM protein structures and predicted models

The structures of this study were solved by initially docking separate AlphaFold3 predictions of BAM_64825_, BAM_82441_, and LlpA-C16 into the cryoSPARC-generated Coulomb potential maps in ChimeraX 1.10^51^. Subsequently, the Phenix 1.20.1 software package, in combination with WinCoot 0.9.8.1, was used for building all structures and structure refinement. Models were rebuilt manually in Coot and refined against the sharpened maps by iterative rounds of real-space refinement in Phenix with secondary-structure and Ramachandran restraints, using the locally refined maps to build the flexible regions. Model quality was assessed and confirmed with the PDB validation web tool (https://validate-rcsb-1.wwpdb.org/), and refinement and validation statistics are reported in Table S2. Images were created using ChimeraX 1.10.

### Bioinformatics analysis of protein sequences

BamA sequences were extracted from the annotated whole-genome assemblies of the isolate panel using an in-house pipeline that searches the Prokka-annotated proteome of each genome for BamA homologues and retains the full-length hit. Sequences were manually validated as full-length, functional BamA, and where a *Pseudomonas* strain encoded more than one BamA homologue, secondary (non-orthologous) copies were removed, giving one BamA sequence per isolate (Supplementary Data 6). Reference *Xanthomonas* and *Burkholderia* BamA sequences were retrieved from the NCBI protein database by BLASTP using the *P. aeruginosa* BamA sequence as the query. All BamA sequences were aligned with Clustal Omega. Extracellular loop 6 sequences were clustered by pairwise identity using hierarchical agglomerative clustering, defining 15 loop 6 groups across the 101 screened strains (Supplementary Data 5), and each strain was assigned a loop 6 type from the presence or absence of the loop 6 insertion in the alignment (alignment columns 41–51). Strains with no insertion were classed as short loop 6 (n = 52) and those carrying a 4–7-residue insertion as long loop 6 (n = 46), with three outgroup strains not assigned to either class. Lectin-like bacteriocins were identified using published, active Llp sequences as BLASTP queries against the NCBI non-redundant protein database^52^. Hits were dereplicated with CD-HIT at a 98% sequence identity threshold and manually curated by domain architecture, retaining 238 *Pseudomonas* Llp sequences whose pairwise amino-acid identity spans 15–98%. Similarly, lectin-like bacteriocins from *Xanthomonas* and *Burkholderia* were identified with BLASTP searches using the sequences of published proteins confirmed to be active, after which newly discovered proteins were manually curated according to fold and sequence, yielding 7 *Xanthomonas* and 14 *Burkholderia* sequences. Multiple sequence alignments were generated using Clustal Omega (https://www.ebi.ac.uk/jdispatcher/msa/clustalo), and percentage identity matrices used for further in-depth analysis^53^. Multiple MSAs were performed to investigate specific clades or selected protein regions of interest, such as C-terminal peptides. To generate phylogenetic trees, IQ-TREE was used with the settings --seqtype AA -m -- msub nuclear --cmin 2 --cmax 10 --merit AIC --ninit 100 --ntop 20 --nbest 5 --nstop 100 --radius 6 --perturb 0.5 --sup-min 0.0 --ufboot 1000 --nmax 1000 --nstep 100 –bcor 0.99 --beps 0.5^54^. The Llp phylogeny in Fig. 1a was inferred from an alignment of 268 sequences comprising the 238 *Pseudomonas* Llps (106 LlpA, 132 LlpB), characterised L-type pyocins and putidacin, and *Burkholderia* (n = 14) and *Xanthomonas* (n = 7) Llp sequences, with the sequence of a plant dual β-lectin protein (ABQ32294) used to root the tree, and branch support was assessed with 1,000 ultrafast bootstrap replicates. Phylogenetic trees were visualised with iTOL^55^. While the genus-level clades and tip-level groups were strongly supported (bootstrap 99–100), several of the deepest backbone nodes were more weakly resolved (∼84–86); the precise branching order among these basal splits should therefore be interpreted with caution.

Relationships within the Llp family were visualised by principal coordinates analysis of the Clustal Omega multiple sequence alignment of 238 Llp sequences. Pairwise amino-acid identity was computed for each pair of sequences over the alignment columns at which neither sequence carried a gap, and converted to a distance as D = 1 − identity. The distance matrix was double-centred and eigendecomposed (classical metric multidimensional scaling), and each Llp was plotted by its position on the first two principal coordinates, coloured by sequence group or by BamA loop 6 targeting preference. No column weighting, sequence-redundancy correction or other preprocessing was applied, and the ordination is a deterministic function of the alignment with no stochastic component. The first two axes account for 28.7 % and 7.6 % of the total variance respectively, expressed relative to the sum of the positive eigenvalues, since the identity distance is not guaranteed to be Euclidean. Ordinations were computed in Python 3.11.15 with NumPy 2.4.4 and scikit-learn 1.8.0.

### Protein structure model prediction and structural analysis of Llp–BamA interfaces

AlphaFold3 was used for the prediction of protein structures using the AlphaFold Server (https://alphafoldserver.com, state 2025 and 2026)^32^. Llp–BamA complexes were predicted as two-chain systems comprising the full-length mature Llp sequence and the BamA β-barrel domain of the target strain. Five models were generated per complex, and the top-ranked model by the server ranking score was retained for analysis. Models were then assessed by interface confidence (ipTM, PAE) and by agreement with the experimental BAM_64825_–LlpA-C16 structure. Interface contacts were computed as heavy-atom pairs within 4.5 Å, separately for the Llp N-terminal β-lectin domain against BamA loop 6 and for the Llp C-terminal peptide against the BamA barrel, and deployment of the C-terminal peptide into the barrel lumen was scored from the minimum distance between the peptide and the barrel centroid. Models were retained when the lectin domain engaged BamA loop 6 in the same orientation as in the cryo-EM structure. Of the 64 LlpA/B–BamA pairs predicted *de novo* in this study, 17 met the ipTM > 0.6 criterion (ipTM 0.66–0.85), while 9 were borderline (ipTM 0.4– 0.6) and 38 showed no confident interface (ipTM < 0.4); models in the lower-ipTM groups also generally failed the experimental-interface criterion, with the C-terminal peptide left undeployed outside the barrel: the median distance from the peptide to the barrel centroid was 35.5 Å in the ipTM < 0.4 group and only 7 of 38 pairs placed the peptide within the lumen, compared with 4.9 Å and 15 of 17 pairs among the ipTM > 0.6 models. Together with previously generated models, this yielded a final structure-guided set spanning short-, long-and broad-loop targeters (Supplementary Data 11). Twelve models were taken forward for analysis of the C-terminal peptide, comprising the Llps whose peptide was modelled deployed into the BamA lumen and β-paired with strand 1. Where the same Llp had been modelled against BamA from more than one strain, a single model was retained, since BamA β-strand 1 is invariant across the panel and the peptide interaction is therefore the same in each. In these models the peptide β-pairs with strand 1, reproducing the arrangement seen in the BAM_64825_– LlpA-C16 and Pyocin L2–BAM cryo-EM structures. Interface residues were defined from the cryo-EM structure and from the retained AlphaFold3 models as heavy-atom contacts within 4.5 Å, computed with a k-d tree search on structures parsed with Biopython (Bio.PDB MMCIFParser/PDBParser).

For the loop 6 interface, contacting Llp positions were mapped onto LlpA-C16 numbering via the Llp multiple-sequence alignment, models were grouped by the loop 6-targeting class of the Llp, and positions present in more than 25% of models within a class were retained as that class’s conserved contact footprint; conservation at these positions was then computed across the corresponding class of the full Llp library and displayed as sequence logos. For the C-terminal peptide interface, the β-augmentation register was defined structurally rather than by sequence: the backbone hydrogen-bond ladder between each Llp C-terminus and BamA β-strand 1 (motif P-S-G-S-I-T-A-S-V) was mapped in the cryo-EM complexes and in the AlphaFold3 models, scoring a hydrogen bond when the minimum donor-N···acceptor-O distance was ≤3.5 Å in either donor/acceptor direction, such that the residue paired to BamA β1 residue 425 defines register position 0 and the residue paired to β1 residue 427 falls at position −2. To propagate this register to all 246 C-terminal peptides, a position-specific scoring matrix was built from the structurally observed registers over offsets −8 to +4 using a log2 likelihood with 0.5 pseudocounts and a background estimated from the C-terminal 25-residue windows of all peptides; every candidate frame within the last 16 residues of each peptide was scored and the highest-scoring frame taken as the register, with the score margin between the best and second-best frame recorded as a confidence tier (high, margin ≥ 3; medium, ≥ 1.2; low, below 1.2). C-terminal similarity groups were defined from the terminal 12 residues of each peptide, the segment that carries the β1-augmenting strand: pairwise alignment scores were computed with the Biopython PairwiseAligner in local mode using BLOSUM62 (gap open −10, gap extend −0.5) and normalised as S(i,j) = s(i,j)/√(s(i,i)·s(j,j)), and the distance matrix D = 1 − S was clustered by hierarchical agglomerative clustering (average linkage) with a flat-cluster cutoff of 0.6, retaining clusters with at least three members and designating them G01– G17 in order of size, with smaller clusters pooled as unassigned. Sequence logos were generated with logomaker, with per-column information content computed as Schneider information, R = log2(20) − (H + eₙ), using the small-sample correction eₙ = (20−1)/(2·ln2·N_occ) and scaling column heights by occupancy so that positions reached by only the longer peptides are not inflated; residues are coloured by chemistry (aromatic FWY, aliphatic AILMV, polar STNQCG, acidic DE, basic KRH, proline).

## Data Availability

Cryo-EM maps and atomic coordinates have been deposited in the Electron Microscopy Data Bank and the Protein Data Bank under accession codes [EMD-80655 / PDB 26HC] (apo BAM_82441_), [EMD-80656 / PDB 26HD] (BAM_64825_–LlpA-C16 monomer) and [EMD-80657 / PDB 26HE] (BAM_64825_–LlpA-C16 dimer). Whole-genome sequencing reads and assemblies for the 111 *Pseudomonas* isolates are available under BioProject <u>PRJNA1530708</u>. Llp and BamA sequences, the Llp × strain susceptibility matrix, loop 6 group assignments, C-terminal peptide alignments and the AlphaFold3 model coordinates are provided as Supplementary Data 1–13. The previously determined Pyocin L2–BAM_P28_ structure used for comparison is available under PDB 9PXJ. All other data supporting the findings of this study are available from the corresponding authors on reasonable request.

## Code Availability

All custom code used for the analyses in this paper is available as llp-bama-analysis_code v1.1 *to be added on final publication*. The package contains the BamA orthologue extraction, the BamA loop 6 grouping and short/long typing, the loop 6 preference classification (long versus short odds ratios with Haldane-Anscombe continuity correction and the dual-engagement reclassification rule), the per-Llp Fisher exact and chi-square test families with Benjamini-Hochberg correction across all 238 Llps, the preference-index comparison between the colony and purified-protein screens, the Mantel and partial Mantel analyses of BamA loop sequence against susceptibility, the AlphaFold3 model triage, the loop 6 interface contact analysis, the C-terminal register assignment and similarity grouping, and the figure panels. It ships with a run_all.sh driver, the derived input tables, and a VERIFICATION.md recording which published values each script reproduces and which remain open. Analyses were run with Python 3.11.16, NumPy 2.4.6, pandas 3.0.5, SciPy 1.17.1, scikit-learn 1.9.1, Biopython 1.88, Matplotlib 3.11.2, openpyxl 3.1.5, logomaker 0.8.7 and BLAST+ 2.17.0. Cryo-EM data processing used the software and versions listed in the Methods and Table S2, and structural predictions used AlphaFold3 with the job configurations provided in Supplementary Data 7 and 11.

## Supporting information

All supplementary data files

## Acknowledgements

The authors acknowledge the use of electron microscopy and cryo-sample preparation facilities at the Ian Holmes Imaging Centre of the Bio21 Molecular Science & Biotechnology Institute, the University of Melbourne. We thank the NSW DPI Plant Pathology & Mycology Herbarium (DAR) for providing *Pseudomonas* isolates used in this study. This research was supported by the ARC discovery project grant awarded to R.G., G.K. and J.P.R.C. (DP230102150). F.M. and R.G. are members of the Australian Research Council Industrial Transformation Training Centre for Cryo-Electron Microscopy of Membrane Proteins for Drug Discovery (IC200100052). R.G. was funded by an NHMRC EL1 investigator grant (APP1197376). G.K. was supported by the Snow Medical Research Foundation (SMRF2021-276). J.P.R.C. is supported by the UKRI Medical Research Council (MR/X007197/1), and Biotechnology and Biological Sciences Research Council (UKRI797).

## Author Contributions

Conceptualization: R.G., F.M.

Data collection: F.M., C.W., L.J.

Data curation: F.M., R.G., C.W., L.J.

Formal analysis: F.M., R.G.

Funding acquisition: R.G., G.J.K., J.P.R.C.

Investigation: F.M., R.G., C.W., L.J.

Methodology: F.M., R.G., G.J.K., J.P.R.C.

Project administration: R.G., F.M.

Resources: R.G.

Supervision: R.G. Validation: F.M., R.G.

Visualization: F.M., R.G.

Writing – original draft: F.M., R.G.

Writing – review and editing: all authors

## Supplementary Tables

**Table S1: Statistical data for correlation between BamA variable extracellular loop sequence and Llp activity.**

**Table S2: Cryo-EM data-processing and model statistics**

**Table S3: Plasmids used in this study for molecular biology**

**Table S4: Primers used in this study**

**Table S5: Strains used in this study**

## Supplementary Data

**Supplementary Data 1:** Llp sequences and tree

**Supplementary Data 2:** Pseudomonas strains

**Supplementary Data 3:** Llp screening inventory

**Supplementary Data 4:** Llp susceptibility matrix

**Supplementary Data 5:** Llp susceptibility analysis

**Supplementary Data 6:** Pseudomonas BamA sequences and tree

**Supplementary Data 7:** BamA AF3 models

**Supplementary Data 8:** Llp loop preference summary

**Supplementary Data 9:** Purified Llp activity

**Supplementary Data 10:** Environmental Llp screening rescored

**Supplementary Data 11:** Llp-BAM AF3 models

**Supplementary Data 12:** Loop 6-binding residues

**Supplementary Data 13:** Llp C-terminal peptide analysis

## Supplementary Figures

**Figure S1:**
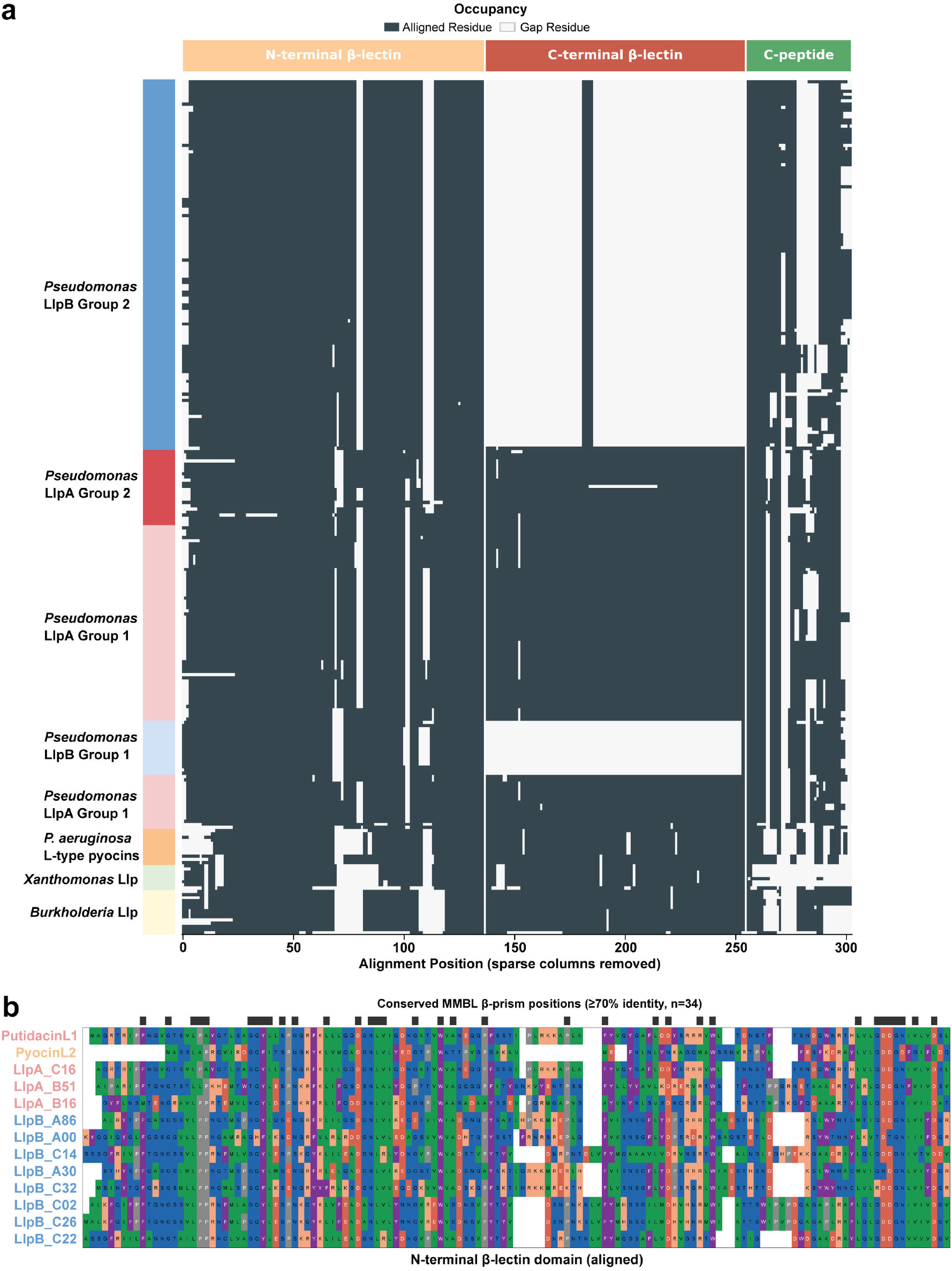
Llp multiple-sequence alignment shows that LlpBs retain the N-terminal β-lectin domain. (a) A shaded representation of the occupancy of the alignment used to construct the Llp phylogeny, showing that the LlpB β-lectin domain aligns with the N-terminal domain from LlpAs, L-type pyocins and non-Pseudomonas Llps. (b) A multiple-sequence alignment of a limited set of LlpAs, Pyocin L2 and LlpBs, showing the alignment and conservation of this domain.

**Figure S2:**
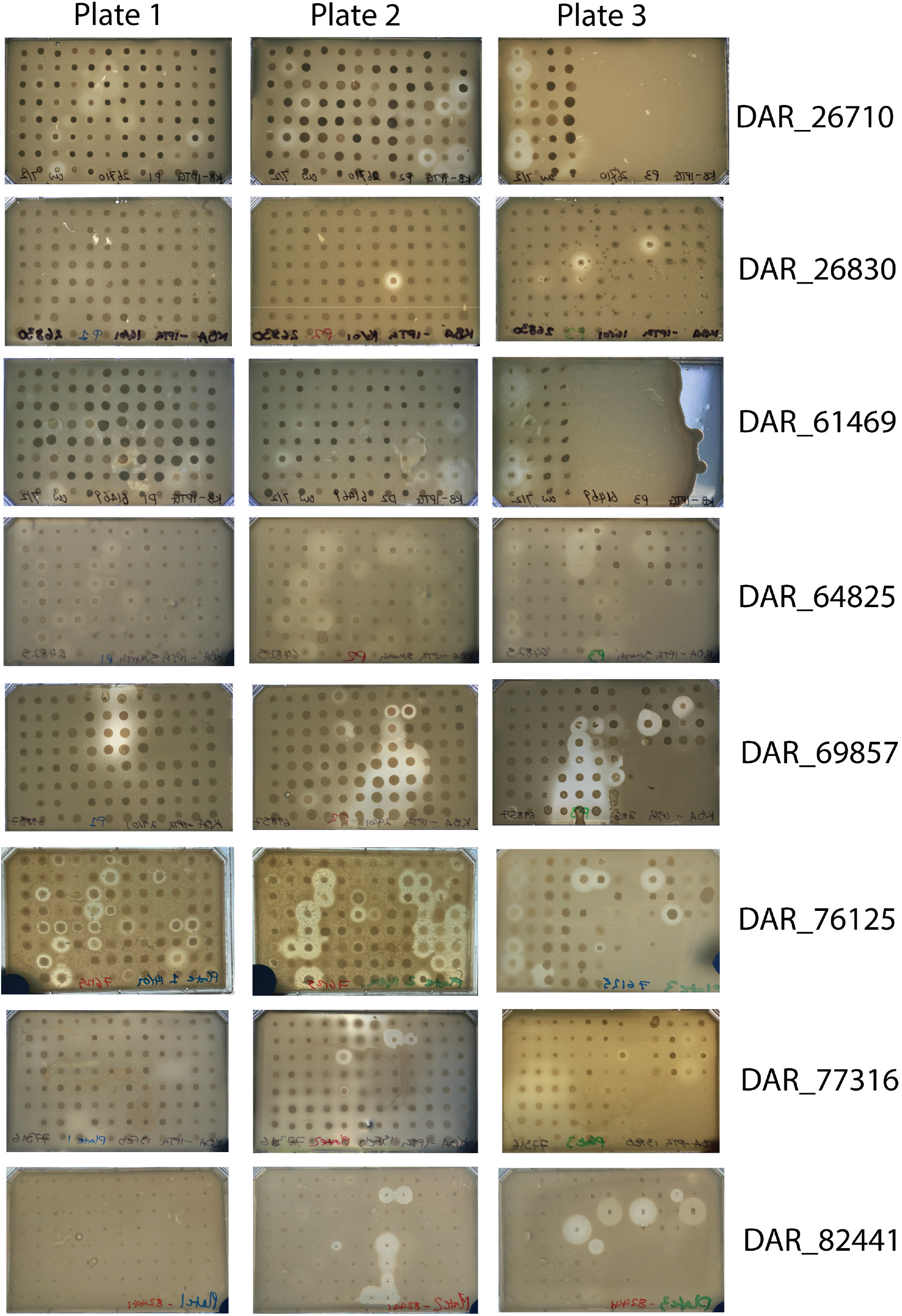
**Examples of agar-overlay inhibition plates from the broad colony-based LlpA/B screen.**

**Figure S3:**
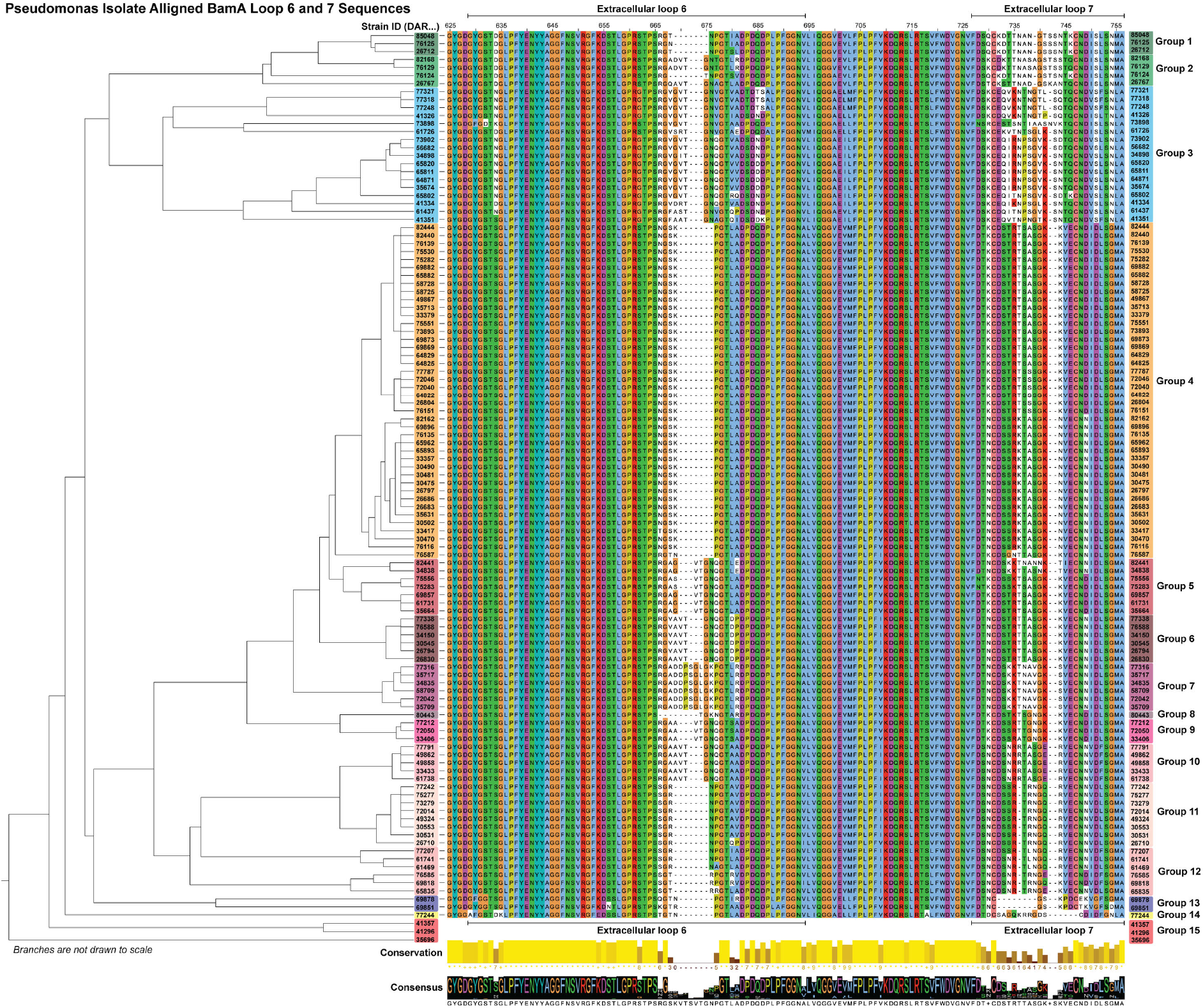
**A phylogeny and multiple sequence alignment of extracellular loops 6 and 7 of BamA from the Pseudomonas isolates used in our LlpA/B activity screen.**

**Figure S4:**
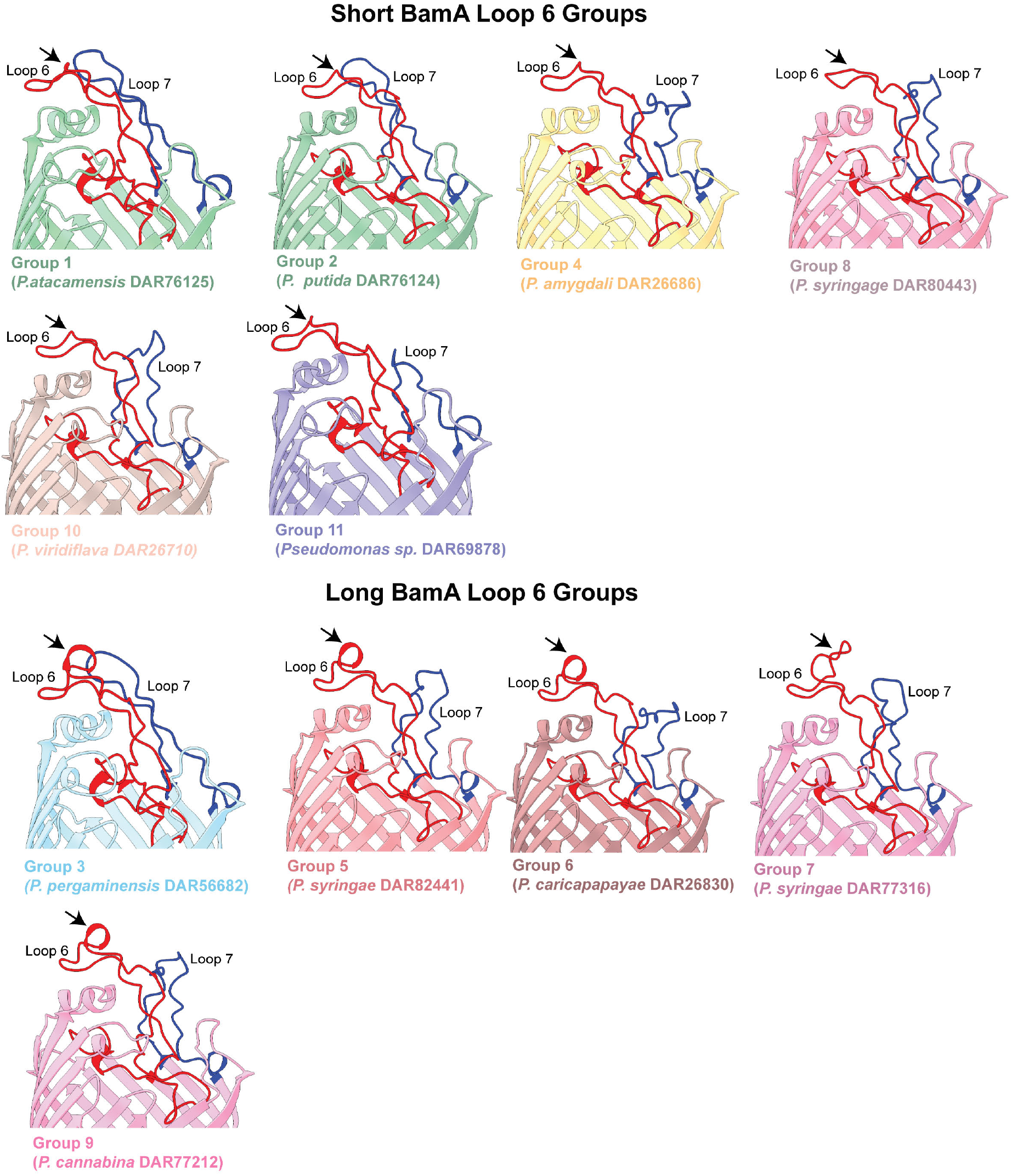
**Example AlphaFold3 models of BamA from different loop 6 sequence groups, showing variation in the structure of loop 6 induced by the long loop expansion.**

**Figure S5.**
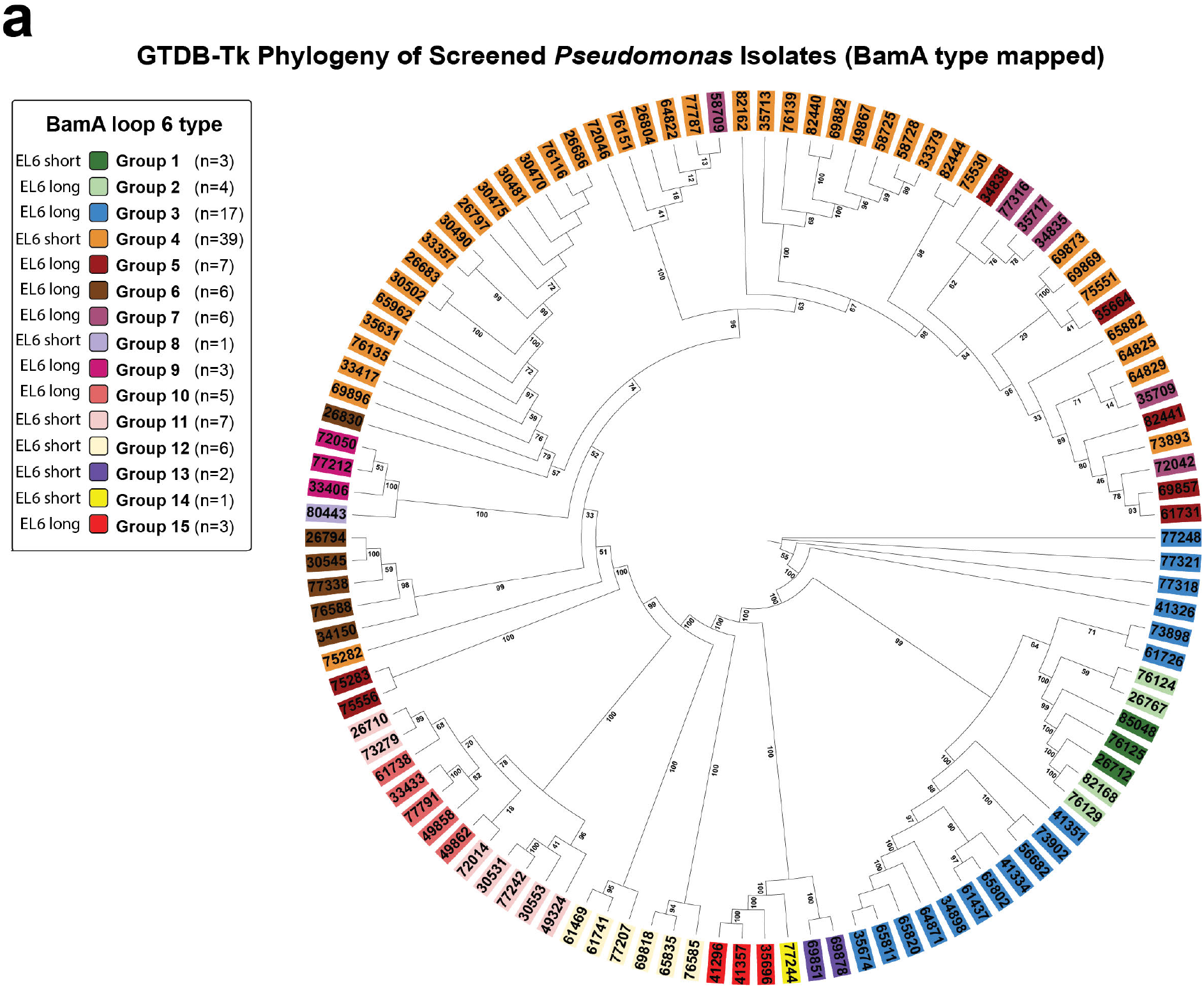
**Genome phylogeny of the sequenced Pseudomonas isolates with BamA loop 6 type mapped**. Maximum-likelihood tree of 110 sequenced isolates inferred with IQ-TREE v2.3.1 from a concatenated alignment of the 120 GTDB bacterial single-copy marker genes (bac120; 5,035 amino-acid positions) extracted and aligned with GTDB-Tk, under the ModelFinder-selected Q.insect+F+I+R3 model; node values are SH-aLRT (%) / ultrafast-bootstrap (%) support from 1,000 replicates. Tips are coloured by BamA loop 6 group (1–15; short and long loop 6 types as keyed), showing that loop 6 type is broadly conserved within clades with several independent long/short transitions across the genus.

**Figure S6.**
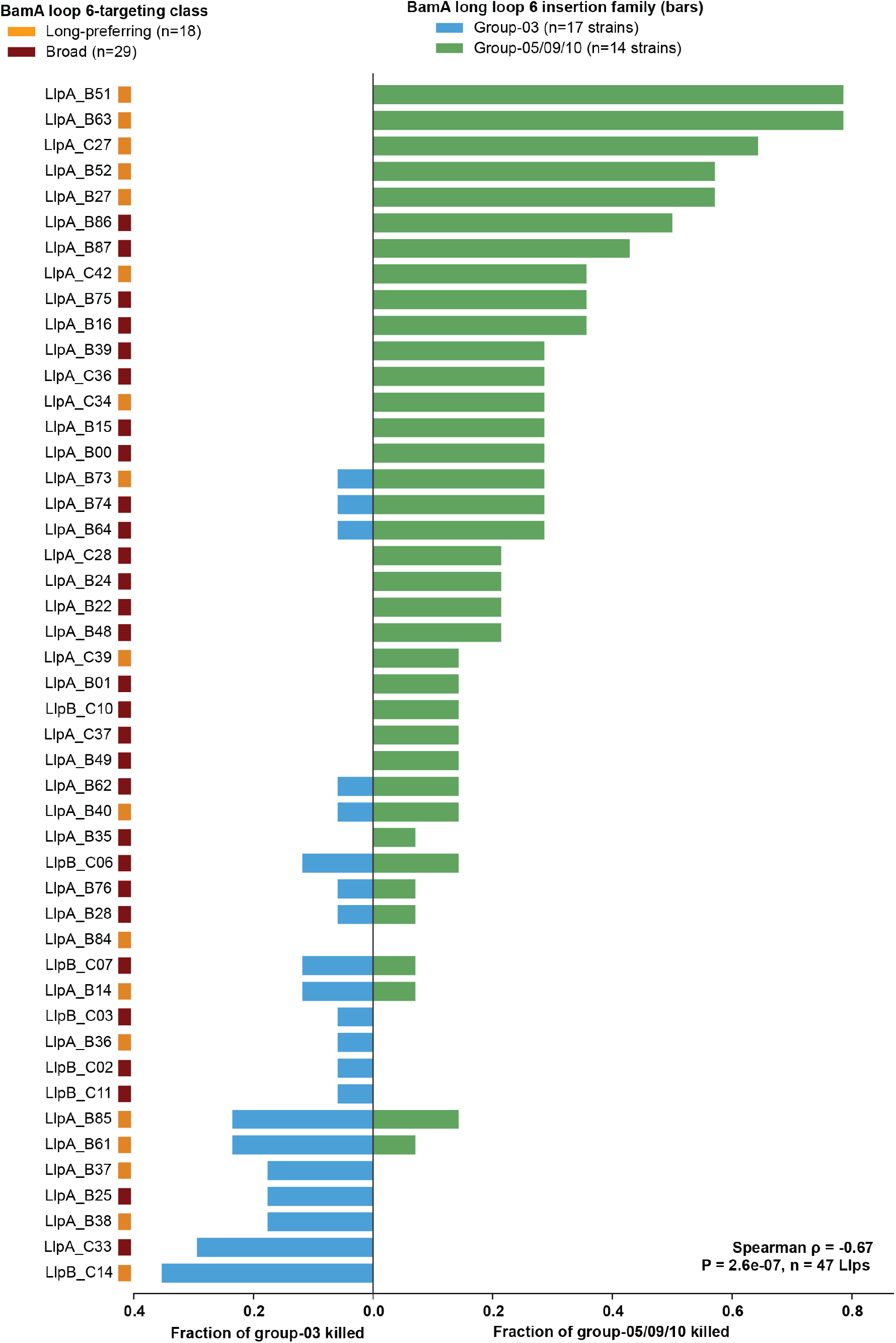
Comparison of LlpA/B activity against strains with BamA long loop 6 type groups 3 and 5, 9 and 10. A strong negative correlation is observed between targeting these long loop types, indicating molecular recognition of both loop groups by LlpA/Bs is difficult.

**Figure S7:**
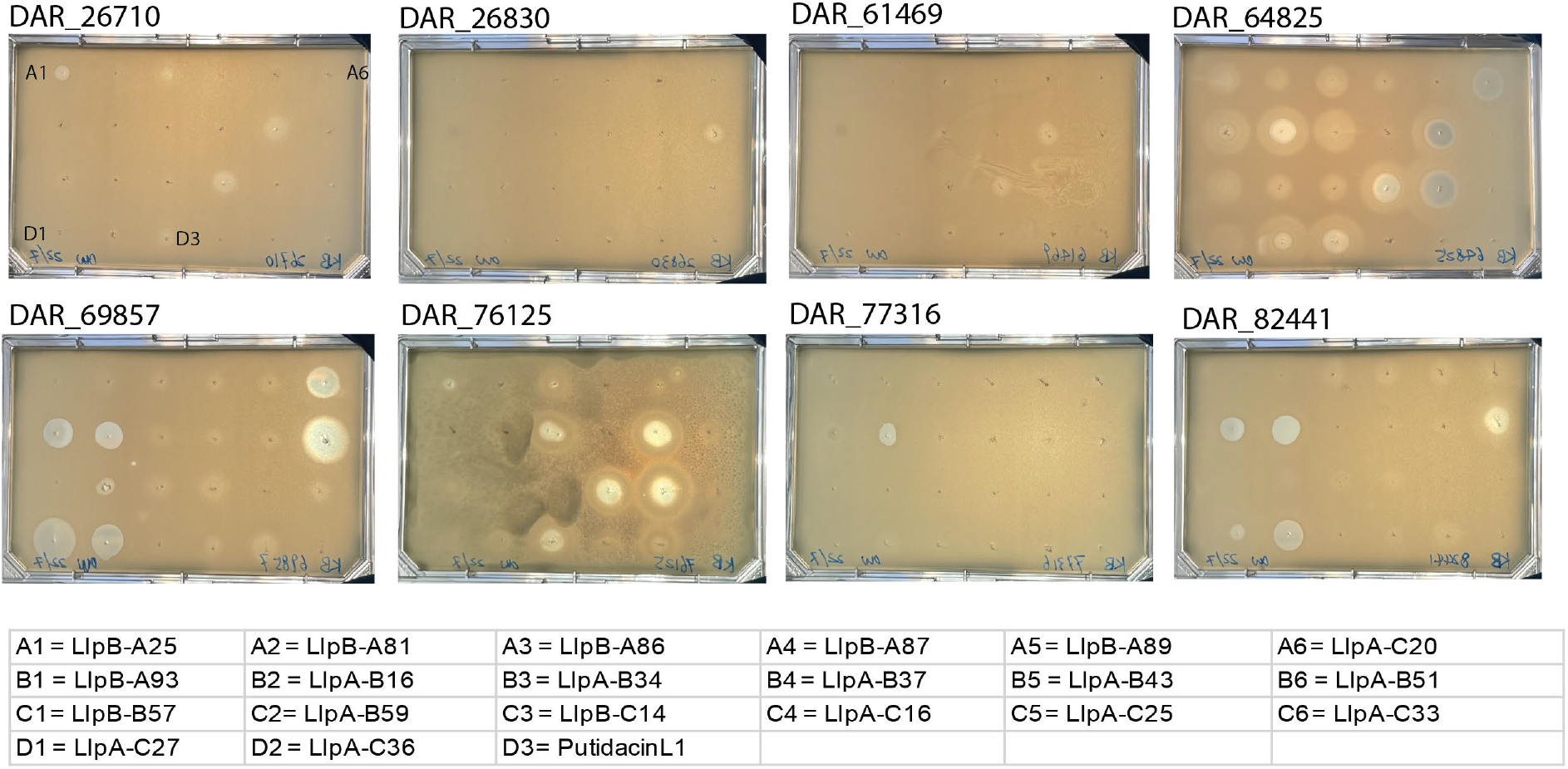
A soft agar overlay inhibition assay with purified LlpA/Bs. Purified N-terminally 6×His-tagged LlpA/B were spotted in a grid pattern labelled in the top left image, with LlpA/B positions shown in the lower table. Pseudomonas strains are indicated by ID number. These are representative images of an experiment conducted three times (n = 3, biological replicates).

**Figure S8:**
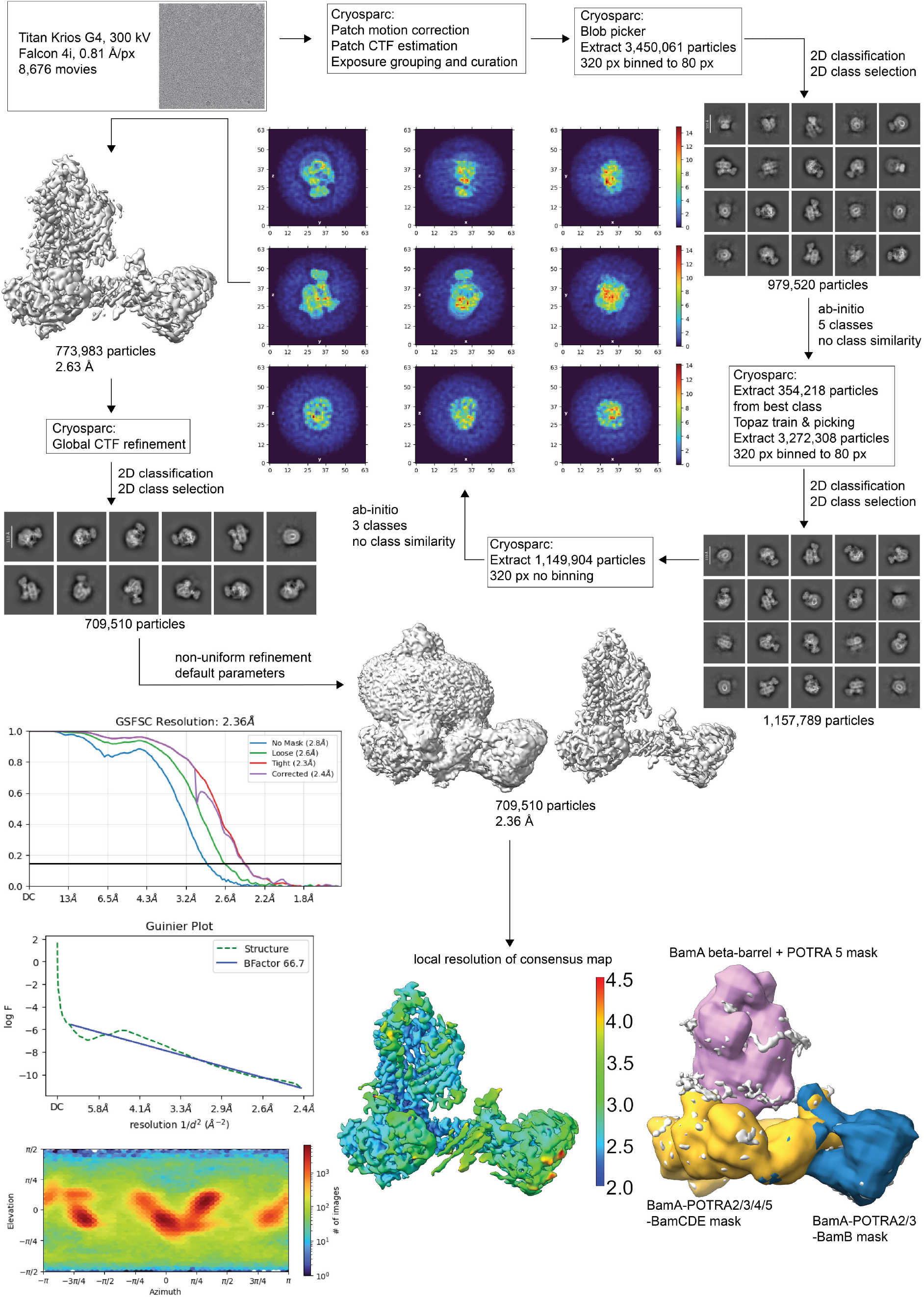
**Cryo-EM processing workflow of apo BAM_82441_.**

**Figure S9:**
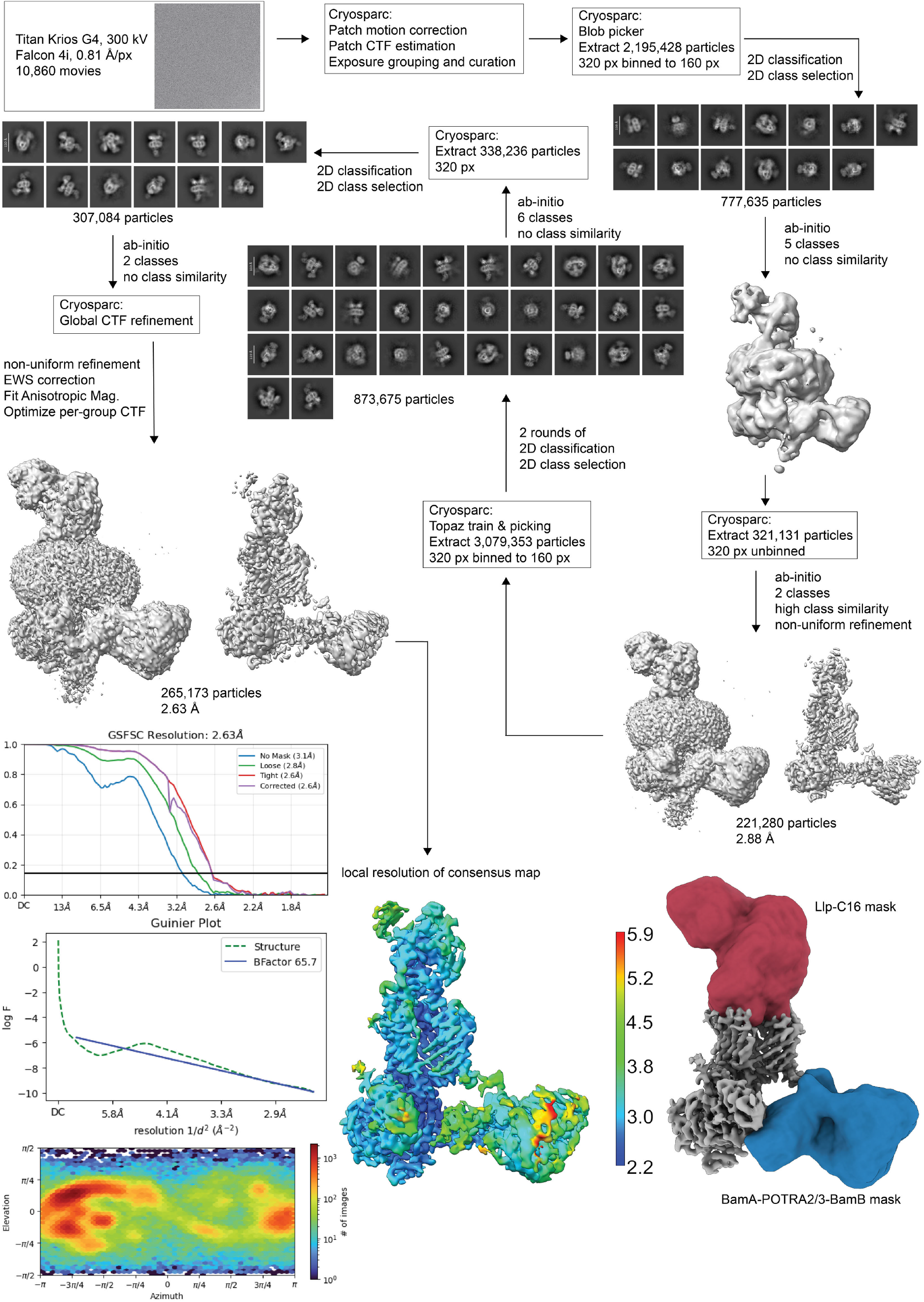
**Cryo-EM processing workflow of BAM_64825_ in complex with LlpA-C16.**

**Figure S10:**
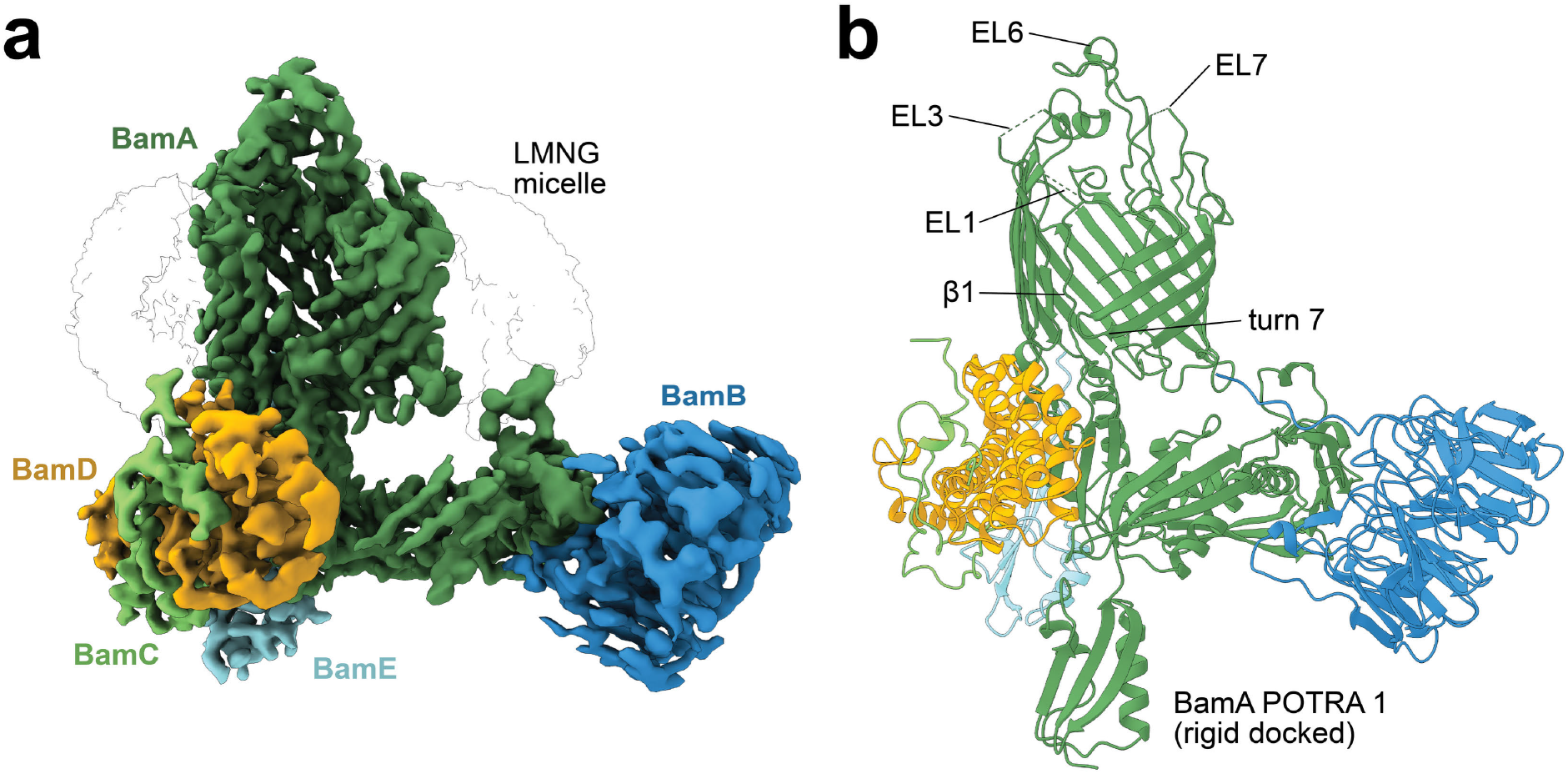
Cryo-EM structure of apo BAM from *Pseudomonas syringae* strain DAR82441. (a) Coulomb potential map of BAM from *Pseudomonas syringae* DAR82441, BamA (dark green), BamB (blue), BamC (green), BamD (orange), and BamE (cyan). (b) The structure of apo BAM_82441_ at 2.36 Å (PDB 26HC). Labelled are the surface-exposed extracellular loop 6 (EL6) of BamA, as well as EL1, EL3, and EL7, which are highly flexible and not completely resolved. The first β-strand (β1) of the BamA β-barrel is flexible in this outward-open BAM conformation and turn 7 as well as POTRA 5 close the β-barrel from the periplasmic side. BamA POTRA 1 was rigidly docked into low-resolution density.

**Figure S11:**
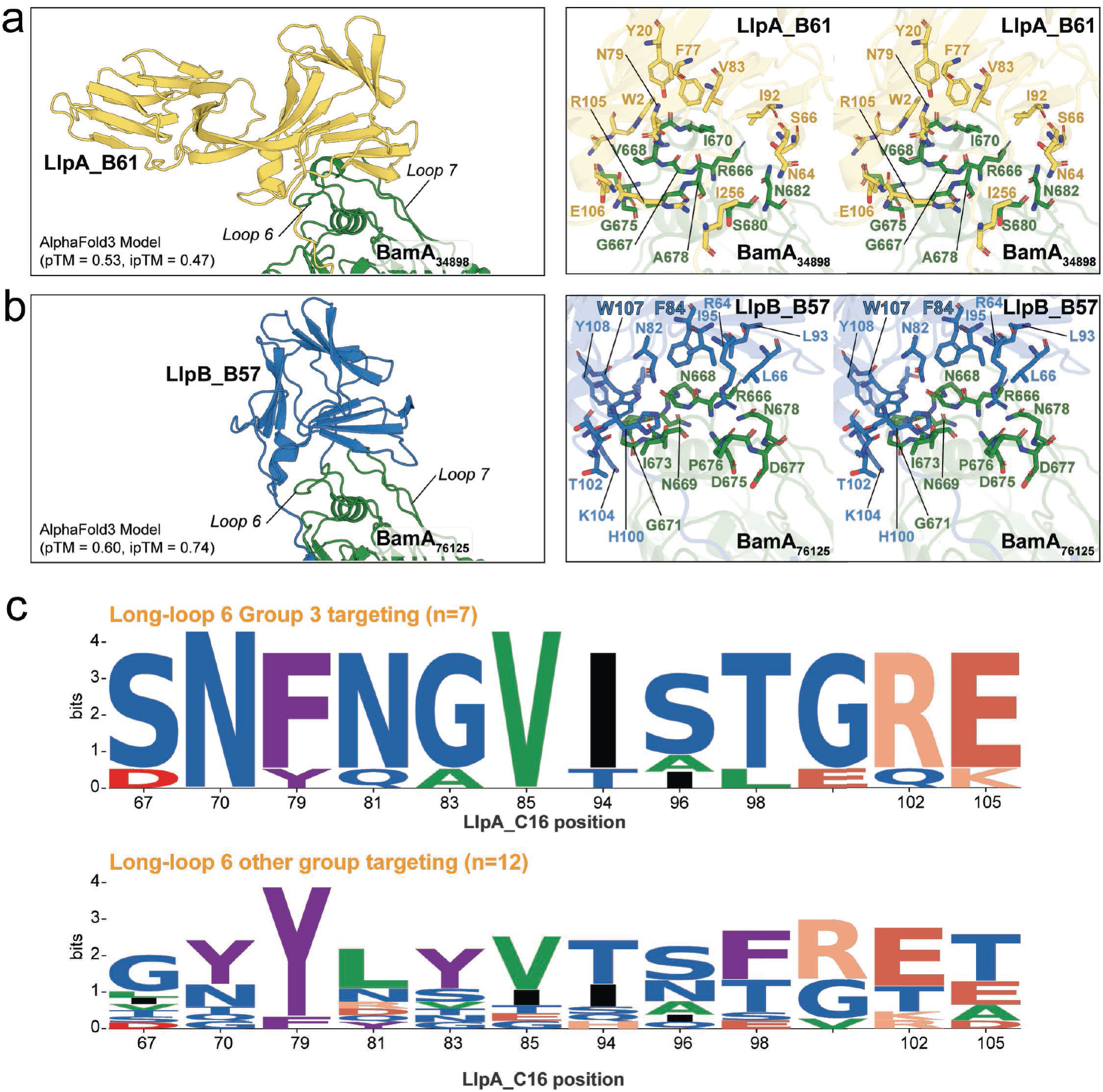
Additional data from AlphaFold3 modelling of BamA loop 6-binding by the LlpA/B N-terminal β-domain interface. (a,b) AlphaFold3 models of LlpA_B61 (long loop 6-targeting), and LlpB_B57 (short loop 6-targeting) in complex with BamA from susceptible Pseudomonas isolates, shown in the orientation and with the same details as BAM_64825_*–LlpA-C16 in* Fig. 4a*. (c) Sequence logos of the conservation of the LlpAIB N-terminal β-lectin domain amino acids from BamA long loop 6-targeting LlpAIBs, separated into LlpAIBs with a targeting preference for BamA loop 6 group 3, vs other long loop groups. Calculated from the sequence alignment of all LlpAIBs in the screening panel*.

**Figure S12:**
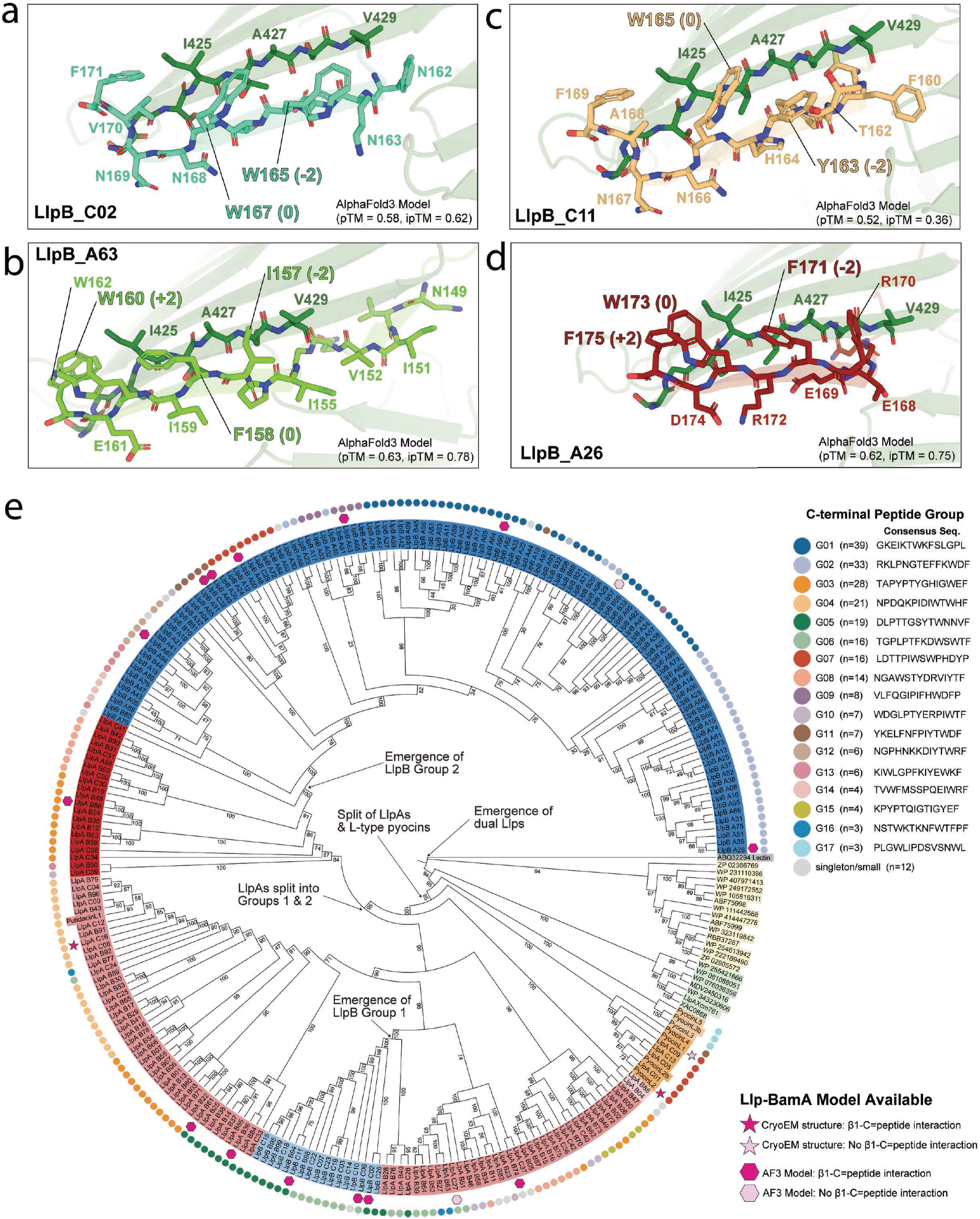
Che C-terminal peptide of Llps is highly di+erse. (a-d) Additional AlphaFold3 models of Llp C-terminal peptides in complex with BamA β-strand 1. (e) The maximum-likelihood (ML) phylogeny of Llp proteins shown in Fig. 1a. Annotated with C-terminal peptide group (see Supplementary Data 13, for full analysis of C-terminal peptide sequences and grouping), and whether experimental structures or plausible AlphaFold3 models are available for the Llp in complex with BamA.

**Figure S13:**
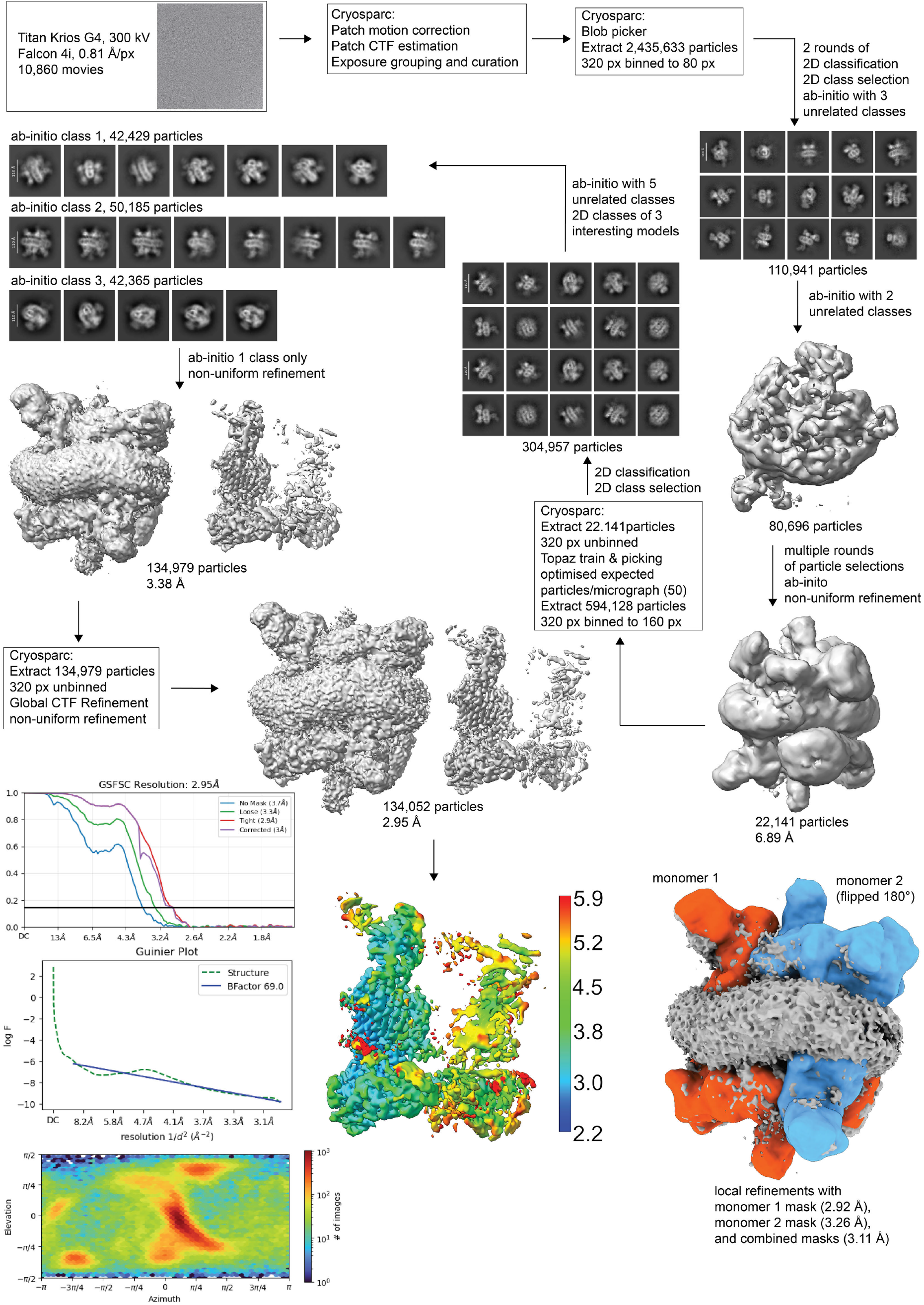
**Cryo-EM processing workflow of dimeric BAM_64825_ in complex with LlpA-C16.**

## References

1 Theuretzbacher, U., Blasco, B., Duffey, M. & Piddock, L. J. Unrealized targets in the discovery of antibiotics for Gram-negative bacterial infections. Nature Reviews Drug Discovery 22, 957–975 (2023).

2 Theuretzbacher, U., Jumde, R. P., Hennessy, A., Cohn, J. & Piddock, L. J. V. Global health perspectives on antibacterial drug discovery and the preclinical pipeline. Nature Reviews Microbiology 23, 474–490 (2025). 10.1038/s41579-025-01167-w

3 Ntallis, C., Martin, N. I., Edwards, A. M. & Weingarth, M. Bacterial cell envelope-targeting antibiotics. Nature Reviews Microbiology 24, 183–196 (2026).

4 Maher, C. & Hassan, K. A. The Gram-negative permeability barrier: tipping the balance of the in and the out. MBio 14, e01205–01223 (2023). 10.1128/mbio.01205-23

5 Draveny, M. & Masi, M. Targeting of the Gram-negative outer membrane for antibiotic discovery and potentiation. ACS Infectious Diseases 12, 1010 (2026).

6 Doyle, M. T., Jimah, J. R., Dowdy, T., Ohlemacher, S. I., Larion, M., Hinshaw, J. E. & Bernstein, H. D. Cryo-EM structures reveal multiple stages of bacterial outer membrane protein folding. Cell 185, 1143–1156.e1113 (2022). 10.1016/j.cell.2022.02.016

7 Kaur, H., Jakob, R. P., Marzinek, J. K., Green, R., Imai, Y., Bolla, J. R., Agustoni, E., Robinson, C. V., Bond, P. J., Lewis, K., Maier, T. & Hiller, S. The antibiotic darobactin mimics a β-strand to inhibit outer membrane insertase. Nature 593, 125-129 (2021). 10.1038/s41586-021-03455-w

8 Imai, Y., Meyer, K. J., Iinishi, A., Favre-Godal, Q., Green, R., Manuse, S., Caboni, M., Mori, M., Niles, S., Ghiglieri, M., Honrao, C., Ma, X., Guo, J. J., Makriyannis, A., Linares-Otoya, L., Böhringer, N., Wuisan, Z. G., Kaur, H., Wu, R., Mateus, A., Typas, A., Savitski, M. M., Espinoza, J. L., O’Rourke, A., Nelson, K. E., Hiller, S., Noinaj, N., Schäberle, T. F., D’Onofrio, A. & Lewis, K. A new antibiotic selectively kills Gram-negative pathogens. Nature 576, 459–464 (2019). 10.1038/s41586-019-1791-1

9 Doyle, M. T. & Bernstein, H. D. Bacterial outer membrane proteins assemble via asymmetric interactions with the BamA β-barrel. Nature Communications 10, 3358 (2019). 10.1038/s41467-019-11230-9

10 Miller, R. D., Iinishi, A., Modaresi, S. M., Yoo, B.-K., Curtis, T. D., Lariviere, P. J., Liang, L., Son, S., Nicolau, S., Bargabos, R., Morrissette, M., Gates, M. F., Pitt, N., Jakob, R. P., Rath, P., Maier, T., Malyutin, A. G., Kaiser, J. T., Niles, S., Karavas, B., Ghiglieri, M., Bowman, S. E. J., Rees, D. C., Hiller, S. & Lewis, K. Computational identification of a systemic antibiotic for Gram-negative bacteria. Nature Microbiology 7, 1661–1672 (2022). 10.1038/s41564-022-01227-4

11 Storek, K. M., Sun, D. & Rutherford, S. T. Inhibitors targeting BamA in gram-negative bacteria. Biochimica et Biophysica Acta (BBA) - Molecular Cell Research 1871, 119609 (2024). 10.1016/j.bbamcr.2023.119609

12 Sun, D., Storek, K. M., Tegunov, D., Yang, Y., Arthur, C. P., Johnson, M., Quinn, J. G., Liu, W., Han, G., Girgis, H. S., Alexander, M. K., Murchison, A. K., Shriver, S., Tam, C., Ijiri, H., Inaba, H., Sano, T., Yanagida, H., Nishikawa, J., Heise, C. E., Fairbrother, W. J., Tan, M.-W., Skelton, N., Sandoval, W., Sellers, B. D., Ciferri, C., Smith, P. A., Reid, P. C., Cunningham, C. N., Rutherford, S. T. & Payandeh, J. The discovery and structural basis of two distinct state-dependent inhibitors of BamA. Nature Communications 15, 8718 (2024). 10.1038/s41467-024-52512-1

13 Storek, K. M., Auerbach, M. R., Shi, H., Garcia, N. K., Sun, D., Nickerson, N. N., Vij, R., Lin, Z., Chiang, N., Schneider, K., Wecksler, A. T., Skippington, E., Nakamura, G., Seshasayee, D., Koerber, J. T., Payandeh, J., Smith, P. A. & Rutherford, S. T. Monoclonal antibody targeting the b-barrel assembly machine of Escherichia coli is bactericidal. Proceedings of the National Academy of Sciences 115, 3692–3697 (2018). 10.1073/pnas.1800043115

14 Cascales, E., Buchanan, S. K., Duché, D., Kleanthous, C., Lloubes, R., Postle, K., Riley, M., Slatin, S. & Cavard, D. Colicin biology. *Microbiology and molecular biology reviews* **71**, 158–229 (2007).

15 Suleman, M., Yaseen, A. R., Ahmed, S., Khan, Z., Irshad, A., Pervaiz, A., Rahman, H. H. & Azhar, M. Pyocins and beyond: exploring the world of bacteriocins in Pseudomonas aeruginosa. Probiotics and Antimicrobial Proteins 17, 240–252 (2025).

16 Parret, A. H. & De Mot, R. Bacteria killing their own kind: novel bacteriocins of Pseudomonas and other γ-proteobacteria. Trends in microbiology 10, 107–112 (2002).

17 Parret, A. H., Wyns, L., De Mot, R. & Loris, R. Overexpression, purification and crystallization of bacteriocin LlpA from Pseudomonas sp. BW11M1. Biological Crystallography 60, 1922–1924 (2004).

18 Ghequire, M. G., Dingemans, J., Pirnay, J. P., De Vos, D., Cornelis, P. & De Mot, R. O serotype-independent susceptibility of Pseudomonas aeruginosa to lectin-like pyocins. Microbiologyopen 3, 875–884 (2014). 10.1002/mbo3.210

19 Ghequire, M. G., Li, W., Proost, P., Loris, R. & De Mot, R. Plant lectin-like antibacterial proteins from phytopathogens Pseudomonas syringae and Xanthomonas citri. Environmental microbiology reports 4, 373–380 (2012). 10.1111/j.1758-2229.2012.00331.x

20 Ghequire, M. G., De Canck, E., Wattiau, P., Van Winge, I., Loris, R., Coenye, T. & De Mot, R. Antibacterial activity of a lectin-like Burkholderia cenocepacia protein. Microbiologyopen 2, 566–575 (2013). 10.1002/mbo3.95

21 Ghequire, M. G., Öztürk, B. & De Mot, R. Lectin-like bacteriocins. Frontiers in Microbiology 9, 2706 (2018).

22 Ghequire, M. G. & De Mot, R. LlpB represents a second subclass of lectin-like bacteriocins. Microbial Biotechnology 12, 567–573 (2019).

23 McCaughey, L. C., Grinter, R., Josts, I., Roszak, A. W., Waløen, K. I., Cogdell, R. J., Milner, J., Evans, T., Kelly, S. & Tucker, N. P. Lectin-like bacteriocins from Pseudomonas spp. utilise D-rhamnose containing lipopolysaccharide as a cellular receptor. PLoS pathogens 10, e1003898 (2014).

24 Munder, F., Johnson, M. D., Samuels, I., McCaughey, L., Zdorevskyi, O., Wang, C., Kropp, A., Zavan, L., Price, E. P., Sarovich, D. S., Varshney, S., McDevitt, C. A., Venugopal, H., Sharma, V., Doyle, M. T., Short, F., Ghosal, D., Connolly, J. P. R., Knott, G. J. & Grinter, R. L-type pyocins inhibit the BAM complex to kill without cell entry. Nature Communications (2026). 10.1038/s41467-026-74995-w

25 Ghequire, M. G., Swings, T., Michiels, J., Buchanan, S. K. & De Mot, R. Hitting with a BAM: selective killing by lectin-like bacteriocins. MBio **9**, 10.1128/mbio.02138-02117 (2018).

26 McCaughey, L. C., Ritchie, N. D., Douce, G. R., Evans, T. J. & Walker, D. Efficacy of species-specific protein antibiotics in a murine model of acute Pseudomonas aeruginosa lung infection. Scientific reports 6, 30201 (2016).

27 Rooney, W. M., Grinter, R. W., Correia, A., Parkhill, J., Walker, D. C. & Milner, J. J. Engineering bacteriocin-mediated resistance against the plant pathogen Pseudomonas syringae. Plant Biotechnology Journal 18, 1296–1306 (2020).

28 Melander, R. J., Zurawski, D. V. & Melander, C. Narrow-spectrum antibacterial agents. *MedChemComm* **9**, 12-21 (2018). 10.1039/c7md00528h

29 Lamichhane, J. R., Osdaghi, E., Behlau, F., Köhl, J., Jones, J. B. & Aubertot, J.-N. Thirteen decades of antimicrobial copper compounds applied in agriculture. A review. Agronomy for Sustainable Development 38, 28 (2018). 10.1007/s13593-018-0503-9

30 Ghequire, M. G. K., Garcia-Pino, A., Lebbe, E. K. M., Spaepen, S., Loris, R. & De Mot, R. Structural Determinants for Activity and Specificity of the Bacterial Toxin LlpA. PLOS Pathogens **9**, e1003199 (2013). 10.1371/journal.ppat.1003199

31 Ghequire, M. G., De Canck, E., Wattiau, P., Van Winge, I., Loris, R., Coenye, T. & De Mot, R. Antibacterial activity of a lectin-like B urkholderia cenocepacia protein. Microbiologyopen 2, 566–575 (2013).

32 Abramson, J., Adler, J., Dunger, J., Evans, R., Green, T., Pritzel, A., Ronneberger, O., Willmore, L., Ballard, A. J. & Bambrick, J. Accurate structure prediction of biomolecular interactions with AlphaFold 3. Nature 630, 493–500 (2024). 10.1038/s41586-024-07487-w

33 Chaumeil, P.-A., Mussig, A. J., Hugenholtz, P. & Parks, D. H. GTDB-Tk: a toolkit to classify genomes with the Genome Taxonomy Database. Bioinformatics 36, 1925–1927 (2020). 10.1093/bioinformatics/btz848

34 Kutik, S., Stojanovski, D., Becker, L., Becker, T., Meinecke, M., Krüger, V., Prinz, C., Meisinger, C., Guiard, B., Wagner, R., Pfanner, N. & Wiedemann, N. Dissecting Membrane Insertion ofC#xa0;Mitochondrial C-Barrel Proteins. Cell 132, 1011–1024 (2008). 10.1016/j.cell.2008.01.028

35 Hagan, C. L., Wzorek, J. S. & Kahne, D. Inhibition of the β-barrel assembly machine by a peptide that binds BamD. Proceedings of the National Academy of Sciences 112, 2011–2016 (2015). doi:10.1073/pnas.1415955112

36 Germany, E. M., Thewasano, N., Imai, K., Maruno, Y., Bamert, R. S., Stubenrauch, C. J., Dunstan, R. A., Ding, Y., Nakajima, Y., Lai, X., Webb, C. T., Hidaka, K., Tan, K. S., Shen, H., Lithgow, T. & Shiota, T. Dual recognition of multiple signals in bacterial outer membrane proteins enhances assembly and maintains membrane integrity. eLife 12, RP90274 (2024). 10.7554/eLife.90274

37 Struyvé, M., Moons, M. & Tommassen, J. Carboxy-terminal phenylalanine is essential for the correct assembly of a bacterial outer membrane protein. Journal of Molecular Biology 218, 141–148 (1991). 10.1016/0022-2836(91)90880-F

38 Tomasek, D., Rawson, S., Lee, J., Wzorek, J. S., Harrison, S. C., Li, Z. & Kahne, D. Structure of a nascent membrane protein as it folds on the BAM complex. Nature 583, 473–478 (2020). 10.1038/s41586-020-2370-1

39 Fox, D. R., Taveneau, C., Clement, J., Grinter, R. & Knott, G. J. Code to complex: AI-driven de novo binder design. Structure 33, 1631–1642 (2025). 10.1016/j.str.2025.08.007

40 Fox, D. R., Asadollahi, K., Samuels, I., Spicer, B. A., Kropp, A., Lupton, C. J., Lim, K., Wang, C., Venugopal, H., Dramicanin, M., Knott, G. J. & Grinter, R. Inhibiting heme piracy by pathogenic Escherichia coli using de novo-designed proteins. Nature Communications 16, 6066 (2025). 10.1038/s41467-025-60612-9

41 Clement, J., Lkhagvajargal, T., Hoare, B. L., Myint, T., Fox, D. R., Wang, C., Knott, G. J., Bathgate, R. A. & Grinter, R. A complete RXFP1-relaxin interaction model unlocks the design of potent mini-protein modulators. bioRxiv, 2026.2006. 2019.733483 (2026).

42 Taveneau, C., Chai, H. X., D’Silva, J., Bamert, R. S., Chen, H., Hayes, B. K., Calvert, R. W., Purcell, J., Curwen, D. J., Munder, F., Martin, L. L., Barr, J. J., Rosenbluh, J., Fareh, M., Grinter, R. & De Knott, G. J. novo design of potent CRISPR–Cas13 inhibitors. Nature Chemical Biology (2026). 10.1038/s41589-025-02136-3

43 Muratspahić, E., Feldman, D., Kim, D. E., Qu, X., Bratovianu, A.-M., Rivera-Sánchez, P., Voss, J. H., Hertz, E. P. T., Jeppesen, M., Dimitri, F., Sakamoto, K., Nallathambi, A., Peceli, P., Cao, J., Cary, B. P., Belousoff, M. J., Keov, P., Trinh, P. N. H., Chen, Q., Ren, Y., Fine, J., Mishra, S., Dalal, A., Sinha, S., Banerjee, R., Ganguly, M., Karuppusamy, K. V., Sappington, I., Schlichthaerle, T., Zhang, J. Z., Pillai, A., Coventry, B., Mihaljević, L., Bauer, M. S., Torres, S. V., Motmaen, A., Lee, G. R., Tran, L., Wang, X., Goreshnik, I., Vafeados, D. K., Svendsen, J. E., Hosseinzadeh, P., Lindegaard, N., Brandt, M., Waltenspühl, Y., Deibler, K., Deweid, L., Bennett, A., Schöppe, J., Dong, T., Yan, X., Oostdyk, L., Cao, W., Anantharaman, L., Weisser, J. J., Bastlund, J. F., Bundgaard, C., Asuni, A. A., English, J. G., Stewart, L. J., Halloran, L., Spangler, J. B., Lieber, A., Shukla, A. K., Sexton, P. M., Roth, B. L., Krumm, B. E., Wootten, D., Tate, C. G., Norn, C. & Baker, D. De novo design of miniproteins targeting GPCRs. Nature (2026). 10.1038/s41586-026-10656-8

44 Hartojo, A., Luu, L. D. W., Adamson, L., Majors, K., Paparella, A. S., Cotter, P. A., Johnson, R. M. & Doyle, M. T. First inhibitor of a bacterial two-partner secretion system. bioRxiv (2026). 10.64898/2026.01.11.698920

45 Zhang, Y., Werling, U. & Edelmann, W. SLiCE: a novel bacterial cell extract-based DNA cloning method. Nucleic acids research 40, e55–e55 (2012).

46 Punjani, A., Rubinstein, J. L., Fleet, D. J. & Brubaker, M. A. cryoSPARC: algorithms for rapid unsupervised cryo-EM structure determination. Nature Methods 14, 290–296 (2017). 10.1038/nmeth.4169

47 Punjani, A., Zhang, H. & Fleet, D. J. Non-uniform refinement: adaptive regularization improves single-particle cryo-EM reconstruction. Nature Methods 17, 1214–1221 (2020). 10.1038/s41592-020-00990-8

48 Zheng, S. Q., Palovcak, E., Armache, J.-P., Verba, K. A., Cheng, Y. & Agard, D. A. MotionCor2: anisotropic correction of beam-induced motion for improved cryo-electron microscopy. Nature Methods 14, 331–332 (2017). 10.1038/nmeth.4193

49. Zivanov, J., Nakane, T. & Scheres, S. H. W. Estimation of high-order aberrations and anisotropic magnification from cryo-EM data sets in RELION-3.1. IUCrJ 7, 253–267 (2020). doi:10.1107/S2052252520000081

50 Bepler, T., Morin, A., Rapp, M., Brasch, J., Shapiro, L., Noble, A. J. & Berger, B. Positive-unlabeled convolutional neural networks for particle picking in cryo-electron micrographs. Nature Methods 16, 1153–1160 (2019). 10.1038/s41592-019-0575-8

51 Meng, E. C., Goddard, T. D., Pettersen, E. F., Couch, G. S., Pearson, Z. J., Morris, J. H. & Ferrin, T. E. UCSF ChimeraX: Tools for structure building and analysis. Protein Science 32, e4792 (2023). 10.1002/pro.4792

52 Altschul, S. F., Gish, W., Miller, W., Myers, E. W. & Lipman, D. J. Basic local alignment search tool. Journal of molecular biology 215, 403–410 (1990).

53 Sievers, F., Wilm, A., Dineen, D., Gibson, T. J., Karplus, K., Li, W., Lopez, R., McWilliam, H., Remmert, M., Söding, J., Thompson, J. D. & Higgins, D. G. Fast, scalable generation of high-quality protein multiple sequence alignments using Clustal Omega. Molecular Systems Biology 7, 539 (2011). 10.1038/msb.2011.75

54 Wong, T. K. F., Ly-Trong, N., Ren, H., Demotte, P., Baños, H., Roger, A. J., Susko, E., Bielow, C., De Maio, N., Goldman, N., Hahn, M. W., dos Reis, M., Vinh, L. S., Huttley, G., Lanfear, R. & Minh, B. Q. IQ-TREE 3: phylogenomic inference software using complex evolutionary models. Molecular Biology and Evolution 43, msag117 (2026). 10.1093/molbev/msag117

55 Letunic, I. & Bork, P. Interactive Tree of Life (iTOL) v6: recent updates to the phylogenetic tree display and annotation tool. Nucleic Acids Research 52, W78–W82 (2024). 10.1093/nar/gkae268

