## Supplementary material for "A diverse family of protein antibiotics inhibits the BAM complex by β-signal mimicry": Table S2

**Table S2: Cryo-EM data-processing and model statistics**

|  | apo BAM_82441_ 26HC | BAM_64825_LlpA-C16 26HD | BAM_64825_LlpA-C16 dimer 26HE |
| --- | --- | --- | --- |

| **Data collection and processing** |  |  |  |
| --- | --- | --- | --- |
| Magnification | 96,000X | 96,000X | 96,000X |
| Voltage (kV) | 300 | 300 | 300 |
| Electron exposure (e–/Å^2^) | 50.24 | 50 | 50 |
| Defocus range (µm) | 0.6-1.2 | 0.6-1.2 | 0.6-1.2 |
| Pixel size (Å) | 0.808 | 0.808 | 0.808 |
| Symmetry imposed | C1 | C1 | C1 |
| Initial particle images (no.) | 3,450,061 | 2,195,426 | 2,435,633 |
| Final particle images (no.) | 709,510 | 221,280 | 134,052 |
| Map resolution (Å) | 2.36 | 2.63 | 2.95 |
| FSC threshold | 0.143 | 0.143 | 0.143 |
| Map resolution range (Å) | 1.99 – 3.72 | 2.24 – 5.06 | 2.69 – 31.51 |
| **Refinement** |  |  |  |
| Initial model used | AlphaFold3 of BAM_82441_ | AlphaFold3 of BAM_64825_ & LlpA-C16 | BAM_64825_-LlpA-C16 (26HD) |
| Model resolution (Å)  FSC threshold | 2.36  0.143 | 2.63  0.143 | 2.95  0.143 |
| Model resolution range (Å) | 1.99 – 3.72 | 2.24 – 5.06 | 2.69 – 31.51 |
| Map sharpening *B* factor (Å^2^) | 66.7 | 64.7 | 69 |
| Model composition  Non-hydrogen atoms  Protein residues  Ligands | 11,443  1,473  0 | 13,670  1,762  0 | 27,294  3,520  0 |
| *B* factors (Å^2^)  Protein  Ligand | 25.99 – 423.73  / | 28.38 – 383.33  / | 11.24 – 1013.96  / |
| R.m.s. deviations  Bond lengths (Å)  Bond angles (°) | 0.002  0.509 | 0.002  0.512 | 0.006  0.648 |
| Validation  MolProbity score  Clashscore  Poor rotamers (%) | 1.69  8.18  0 | 1.64  7.91  0 | 1.77  11.32  0 |
| Ramachandran plot  Favored (%)  Allowed (%)  Disallowed (%) | 96.36  3.64  0 | 96.69  3.31  0 | 96.8  3.2  0 |
