## Supplementary material for "A diverse family of protein antibiotics inhibits the BAM complex by β-signal mimicry": Table S3

**Table S3: Plasmids used in this study.**

| **Plasmid** | **Details** | **Purpose** | **Origin** |
| --- | --- | --- | --- |
| pScRhaB2-BAM-64825 | N-terminal twin-strep tag at BamA;  Genes: BamA, BamD, BamE, BamB, BamC | For overexpression in *E. coli* BL21(DE3)C41 and subsequent purification | 2 gene fragments ordered from Twist Bioscience. *P. putida* DAR64825 |
| pScRhaB2-BAM-82441 | N-terminal twin-strep tag at BamA;  Genes: BamA, BamD, BamE, BamB, BamC | For overexpression in *E. coli* BL21(DE3)C41 and subsequent purification | 2 gene fragments ordered from Twist Bioscience. *P. syringae* DAR82441 |
| pET29b-LlpB-A25 | N-terminal His6-tag | For overexpression in *E. coli* BL21(DE3)C41 | Q5 Site-Directed Mutagenesis |
| pET29b-LlpB-A93 | N-terminal His6-tag | For overexpression in *E. coli* BL21(DE3)C41 | Q5 Site-Directed Mutagenesis |
| pET29b-LlpB-B57 | N-terminal His6-tag | For overexpression in *E. coli* BL21(DE3)C41 | Q5 Site-Directed Mutagenesis |
| pET29b-LlpB-C27 | N-terminal His6-tag | For overexpression in *E. coli* BL21(DE3)C41 | Synthesized by Twist Bioscience |
| pET29b-LlpB-A81 | N-terminal His6-tag | For overexpression in *E. coli* BL21(DE3)C41 | Q5 Site-Directed Mutagenesis |
| pET29b-LlpB-B16 | N-terminal His6-tag | For overexpression in *E. coli* BL21(DE3)C41 | Q5 Site-Directed Mutagenesis |
| pET29b-LlpA-B59 | N-terminal His6-tag | For overexpression in *E. coli* BL21(DE3)C41 | Q5 Site-Directed Mutagenesis |
| pET29b-LlpA-C36 | N-terminal His6-tag | For overexpression in *E. coli* BL21(DE3)C41 | Synthesized by Twist Bioscience |
| pET29b-LlpB-A86 | N-terminal His6-tag | For overexpression in *E. coli* BL21(DE3)C41 | Q5 Site-Directed Mutagenesis |
| pET29b-LlpA-B34 | N-terminal His6-tag | For overexpression in *E. coli* BL21(DE3)C41 | Q5 Site-Directed Mutagenesis |
| pET29b-LlpB-C14 | N-terminal His6-tag | For overexpression in *E. coli* BL21(DE3)C41 | Q5 Site-Directed Mutagenesis |
| pET29b-LlpB-A87 | N-terminal His6-tag | For overexpression in *E. coli* BL21(DE3)C41 | Q5 Site-Directed Mutagenesis |
| pET29b-LlpA-B37 | N-terminal His6-tag | For overexpression in *E. coli* BL21(DE3)C41 | Q5 Site-Directed Mutagenesis |
| pET29b-LlpA-C16 | N-terminal His6-tag | For overexpression in *E. coli* BL21(DE3)C41 | Q5 Site-Directed Mutagenesis |
| pET29b-LlpB-A89 | N-terminal His6-tag | For overexpression in *E. coli* BL21(DE3)C41 | Q5 Site-Directed Mutagenesis |
| pET29b-LlpA-B43 | N-terminal His6-tag | For overexpression in *E. coli* BL21(DE3)C41 | Q5 Site-Directed Mutagenesis |
| pET29b-LlpA-C25 | N-terminal His6-tag | For overexpression in *E. coli* BL21(DE3)C41 | Synthesized by Twist Bioscience |
| pET29b-LlpA-C20 | N-terminal His6-tag | For overexpression in *E. coli* BL21(DE3)C41 | Q5 Site-Directed Mutagenesis |
| pET29b-LlpA-B51 | N-terminal His6-tag | For overexpression in *E. coli* BL21(DE3)C41 | Q5 Site-Directed Mutagenesis |
| pET29b-LlpA-C33 | N-terminal His6-tag | For overexpression in *E. coli* BL21(DE3)C41 | Synthesized by Twist Bioscience |
| pET29b-Llp | No His6-tag; All Llps used in this study | For expression in *E. coli* BL21(DE3)C41 in colony-based growth inhibition screens | Synthesized by Twist Bioscience |
| pET22b-PutidacinL1 | N-terminal His6-tag | For overexpression in *E. coli* BL21(DE3)C41 | Previous work |
