## Supplementary material for "A diverse family of protein antibiotics inhibits the BAM complex by β-signal mimicry": Table S4

**Table S4: Primers used in this study.**

| **Primer** | **Sequence** | **Purpose** |
| --- | --- | --- |
| Q5 SDM forward | CTCGAGCACCACCAC | Forward primer for Q5 Site-Directed Mutagenesis of pET29b-Llp plasmids to remove stop codon |
| WP_110604901.1 reverse | GAAGTGCCACGTCCAAATG | Reverse primer for Q5 Site-Directed Mutagenesis of pET29b-LlpB-B43 |
| WP_123443777.1  reverse | AAATTCCCACTTGAAAAATTCAG | Reverse primer for Q5 Site-Directed Mutagenesis of pET29b-LlpB-A25 |
| WP_095964177.1  reverse | CCATTCCCAATGAAAAATCG | Reverse primer for Q5 Site-Directed Mutagenesis of pET29b-LlpB-A81 |
| WP_201231604.1  reverse | GAAGTCCCATTTGTAAACCTG | Reverse primer for Q5 Site-Directed Mutagenesis of pET29b-LlpB-A86 |
| WP_015097182.1  reverse | CAAAGGGTCTAAACTGAACTTC | Reverse primer for Q5 Site-Directed Mutagenesis of pET29b-LlpB-A87 |
| WP_083454196.1  reverse | CAGAGGTGGCAGTGAG | Reverse primer for Q5 Site-Directed Mutagenesis of pET29b-LlpB-A89 |
| WP_242485727.1  reverse | CAGGGGACCCAGTGAG | Reverse primer for Q5 Site-Directed Mutagenesis of pET29b-LlpB-A93 |
| WP_060708345.1  reverse | AAATTTGTAAATTGGTTGATCATAGG | Reverse primer for Q5 Site-Directed Mutagenesis of pET29b-LlpB-B16 |
| WP_042557906.1  reverse | AAAGGTCCACACCGG | Reverse primer for Q5 Site-Directed Mutagenesis of pET29b-LlpA-B34 |
| WP_161777916.1  reverse | AAACACGTTGTTCCAGG | Reverse primer for Q5 Site-Directed Mutagenesis of pET29b-LlpA-B37 |
| WP_109522175.1  reverse | AAAAGTCCAAATAAAATCTTTAAAGGTC | Reverse primer for Q5 Site-Directed Mutagenesis of pET29b-LlpA-B51 |
| WP_207267578.1  reverse | CAGAGGTCCGAGGCTG | Reverse primer for Q5 Site-Directed Mutagenesis of pET29b-LlpB-B57 |
| WP_122293702.1  reverse | AAAATCCCAGCCGATC | Reverse primer for Q5 Site-Directed Mutagenesis of pET29b-LlpA-B59 |
| WP_236265098.1  reverse | AAACGTCCAACTCCATAC | Reverse primer for Q5 Site-Directed Mutagenesis of pET29b-LlpB-C14 |
| WP_050704433.1  reverse | AAAATGCCAGGTCCAAATG | Reverse primer for Q5 Site-Directed Mutagenesis of pET29b-LlpA-C16 |
| WP_074893631.1  reverse | AAATGTGTAAATCACACGATCG | Reverse primer for Q5 Site-Directed Mutagenesis of pET29b-LlpA-C20 |
