## Supplementary material for "A diverse family of protein antibiotics inhibits the BAM complex by β-signal mimicry": Table S5

**Table S5: Strains used in this study.**

| **Organism** | **Strain** | **Purpose** |
| --- | --- | --- |
| *Escherichia coli* | DH5α | Cloning |
| *Escherichia coli* | BL21(DE3)C41 | Protein production and purification |
| *Pseudomonas fluorescens* | DAR26712 | Growth inhibition assays |
| *Pseudomonas chlororaphis* | DAR76125 | Growth inhibition assays |
| *Pseudomonas granadensis* | DAR85048 | Growth inhibition assays |
| *Pseudomonas fluorescens* | DAR26767 | Growth inhibition assays |
| *Pseudomonas chlororaphis* | DAR76124 | Growth inhibition assays |
| *Pseudomonas fuscovaginae* | DAR82168 | Growth inhibition assays |
| *Pseudomonas putida* | DAR76129 | Growth inhibition assays |
| *Pseudomonas fluorescens* | DAR34898 | Growth inhibition assays |
| *Pseudomonas tolaasii* | DAR35674 | Growth inhibition assays |
| *Pseudomonas tolaasii* | DAR41326 | Growth inhibition assays |
| *Pseudomonas tolaasii* | DAR41351 | Growth inhibition assays |
| *Pseudomonas fluorescens* | DAR56682 | Growth inhibition assays |
| *Pseudomonas sp.* | DAR61437 | Growth inhibition assays |
| *Pseudomonas sp.* | DAR64871 | Growth inhibition assays |
| *Pseudomonas aureofaciens* | DAR65802 | Growth inhibition assays |
| *Pseudomonas putida* | DAR65811 | Growth inhibition assays |
| *Pseudomonas chlororaphis* | DAR73898 | Growth inhibition assays |
| *Pseudomonas putida* | DAR73902 | Growth inhibition assays |
| *Pseudomonas putida* | DAR77318 | Growth inhibition assays |
| *Pseudomonas aureofaciens* | DAR65820 | Growth inhibition assays |
| *Pseudomonas fuscovaginae* | DAR77321 | Growth inhibition assays |
| *Pseudomonas marginalis* | DAR41334 | Growth inhibition assays |
| *Pseudomonas syringae* | DAR77248 | Growth inhibition assays |
| *Pseudomonas syringae* | DAR61726 | Growth inhibition assays |
| *Pseudomonas savastanoi* | DAR26683 | Growth inhibition assays |
| *Pseudomonas savastanoi* | DAR26686 | Growth inhibition assays |
| *Pseudomonas savastanoi* | DAR26797 | Growth inhibition assays |
| *Pseudomonas syringae* | DAR26804 | Growth inhibition assays |
| *Pseudomonas savastanoi* | DAR30475 | Growth inhibition assays |
| *Pseudomonas savastanoi* | DAR30481 | Growth inhibition assays |
| *Pseudomonas savastanoi* | DAR30490 | Growth inhibition assays |
| *Pseudomonas syringae* | DAR30502 | Growth inhibition assays |
| *Pseudomonas savastanoi* | DAR33357 | Growth inhibition assays |
| *Pseudomonas syringae pv. pisi* | DAR33379 | Growth inhibition assays |
| *Pseudomonas syringae pv. morsprunorum* | DAR33417 | Growth inhibition assays |
| *Pseudomonas sp.* | DAR35631 | Growth inhibition assays |
| *Pseudomonas viridiflava* | DAR35713 | Growth inhibition assays |
| *Pseudomonas syringae pv. syringae* | DAR49867 | Growth inhibition assays |
| *Pseudomonas syringae pv. syringae* | DAR58725 | Growth inhibition assays |
| *Pseudomonas syringae pv. coronafaciens* | DAR58728 | Growth inhibition assays |
| *Pseudomonas syringae* | DAR64822 | Growth inhibition assays |
| *Pseudomonas putida* | DAR64825 | Growth inhibition assays |
| *Pseudomonas syringae* | DAR64829 | Growth inhibition assays |
| *Pseudomonas syringae pv. pisi* | DAR65882 | Growth inhibition assays |
| *Pseudomonas syringae pv. mori* | DAR65962 | Growth inhibition assays |
| *Pseudomonas syringae pv. pisi* | DAR69869 | Growth inhibition assays |
| *Pseudomonas syringae pv. pisi* | DAR69873 | Growth inhibition assays |
| *Pseudomonas syringae pv. syringae* | DAR69882 | Growth inhibition assays |
| *Pseudomonas syringae* | DAR69896 | Growth inhibition assays |
| *Pseudomonas syringae* | DAR73893 | Growth inhibition assays |
| *Pseudomonas syringae* | DAR75282 | Growth inhibition assays |
| *Pseudomonas syringae pv. syringae* | DAR75530 | Growth inhibition assays |
| *Pseudomonas syringae* | DAR75551 | Growth inhibition assays |
| *Pseudomonas savastanoi pv. phaseolicola* | DAR76116 | Growth inhibition assays |
| *Pseudomonas savastanoi pv. savastanoi* | DAR76135 | Growth inhibition assays |
| *Pseudomonas syringae pv. syringae* | DAR76139 | Growth inhibition assays |
| *Pseudomonas syringae pv. lachrymans* | DAR76151 | Growth inhibition assays |
| *Pseudomonas syringae pv. syringae* | DAR77787 | Growth inhibition assays |
| *Pseudomonas syringae pv. syringae* | DAR82440 | Growth inhibition assays |
| *Pseudomonas syringae pv. syringae* | DAR82444 | Growth inhibition assays |
| *Pseudomonas syringae* | DAR72046 | Growth inhibition assays |
| *Pseudomonas syringae pv. syringae* | DAR82162 | Growth inhibition assays |
| *Pseudomonas savastanoi pv. phaseolicola* | DAR30470 | Growth inhibition assays |
| *Pseudomonas syringae pv. tabaci* | DAR65893 | Growth inhibition assays |
| *Pseudomonas sp.* | DAR34838 | Growth inhibition assays |
| *Pseudomonas syringae pv. tomato* | DAR35664 | Growth inhibition assays |
| *Pseudomonas syringae pv. lachrymans* | DAR61731 | Growth inhibition assays |
| *Pseudomonas syringae* | DAR69857 | Growth inhibition assays |
| *Pseudomonas syringae pv. porri* | DAR75283 | Growth inhibition assays |
| *Pseudomonas syringae pv. porri* | DAR75556 | Growth inhibition assays |
| *Pseudomonas syringae pv. syringae* | DAR82441 | Growth inhibition assays |
| *Pseudomonas syringae pv. tomato* | DAR26794 | Growth inhibition assays |
| *Pseudomonas syringae pv. syringae* | DAR26830 | Growth inhibition assays |
| *Pseudomonas syringae pv. tomato* | DAR30545 | Growth inhibition assays |
| *Pseudomonas syringae* | DAR34150 | Growth inhibition assays |
| *Pseudomonas syringae pv. maculicola* | DAR76588 | Growth inhibition assays |
| *Pseudomonas syringae pv. maculicola* | DAR77338 | Growth inhibition assays |
| *Pseudomonas sp.* | DAR34835 | Growth inhibition assays |
| *Pseudomonas syringae pv. syringae* | DAR35709 | Growth inhibition assays |
| *Pseudomonas sp.* | DAR35717 | Growth inhibition assays |
| *Pseudomonas syringae* | DAR58709 | Growth inhibition assays |
| *Pseudomonas syringae pv. syringae* | DAR72042 | Growth inhibition assays |
| *Pseudomonas syringae pv. aptata* | DAR77316 | Growth inhibition assays |
| *Pseudomonas syringae pv. lachrymans* | DAR80443 | Growth inhibition assays |
| *Pseudomonas syringae pv. maculicola* | DAR33406 | Growth inhibition assays |
| *Pseudomonas syringae pv. coriandricola* | DAR72050 | Growth inhibition assays |
| *Pseudomonas syringae* | DAR77212 | Growth inhibition assays |
| *Pseudomonas sp.* | DAR33433 | Growth inhibition assays |
| *Pseudomonas viridiflava* | DAR49858 | Growth inhibition assays |
| *Pseudomonas viridiflava* | DAR49862 | Growth inhibition assays |
| *Pseudomonas viridiflava* | DAR61738 | Growth inhibition assays |
| *Pseudomonas viridiflava* | DAR77791 | Growth inhibition assays |
| *Pseudomonas cichorii* | DAR26710 | Growth inhibition assays |
| *Pseudomonas viridiflava* | DAR30531 | Growth inhibition assays |
| *Pseudomonas syringae pv. maculicola* | DAR30553 | Growth inhibition assays |
| *Pseudomonas viridiflava* | DAR49324 | Growth inhibition assays |
| *Pseudomonas cichorii* | DAR72014 | Growth inhibition assays |
| *Pseudomonas viridiflava* | DAR77242 | Growth inhibition assays |
| *Pseudomonas syringae pv. maculicola* | DAR73279 | Growth inhibition assays |
| *Pseudomonas cichorii* | DAR61469 | Growth inhibition assays |
| *Pseudomonas cichorii* | DAR61741 | Growth inhibition assays |
| *Pseudomonas syringae pv. actinidiae* | DAR65835 | Growth inhibition assays |
| *Pseudomonas syringae pv. syringae* | DAR76585 | Growth inhibition assays |
| *Pseudomonas cichorii* | DAR77207 | Growth inhibition assays |
| *Pseudomonas cichorii* | DAR69818 | Growth inhibition assays |
| *Pseudomonas fluorescens* | DAR69851 | Growth inhibition assays |
| *Pseudomonas sp.* | DAR69878 | Growth inhibition assays |
| *Pseudomonas mendocina* | DAR77244 | Growth inhibition assays |
| *Pseudomonas flectens* | DAR35696 | Growth inhibition assays |
| *Pseudomonas aeruginosa* | DAR41296 | Growth inhibition assays |
| *Pseudomonas aeruginosa* | DAR41357 | Growth inhibition assays |
