## Supplementary figures and images for "A diverse family of protein antibiotics inhibits the BAM complex by β-signal mimicry"

### fold_bama30545_llpa_b35_full_data_0.png

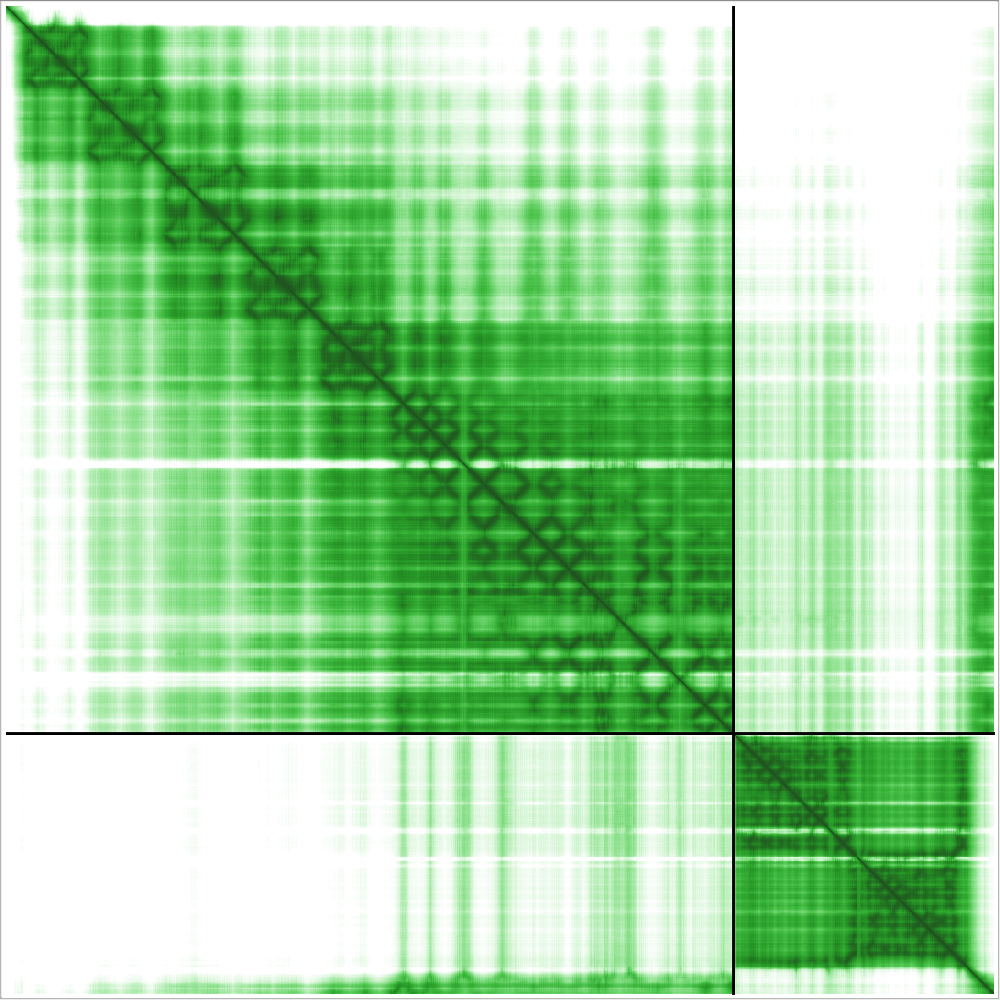

### fold_bama34898_llpa_b36_full_data_0.png

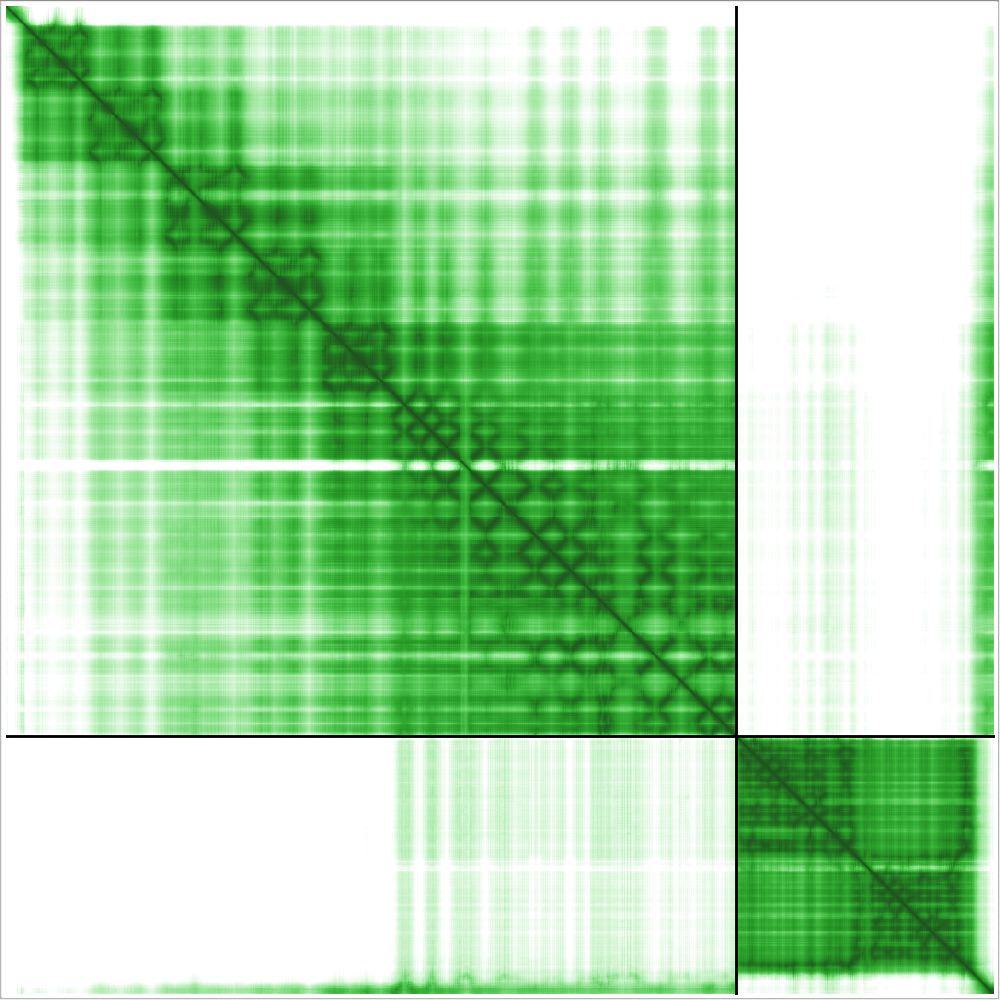

### fold_bama34898_llpa_b61_full_data_0.png

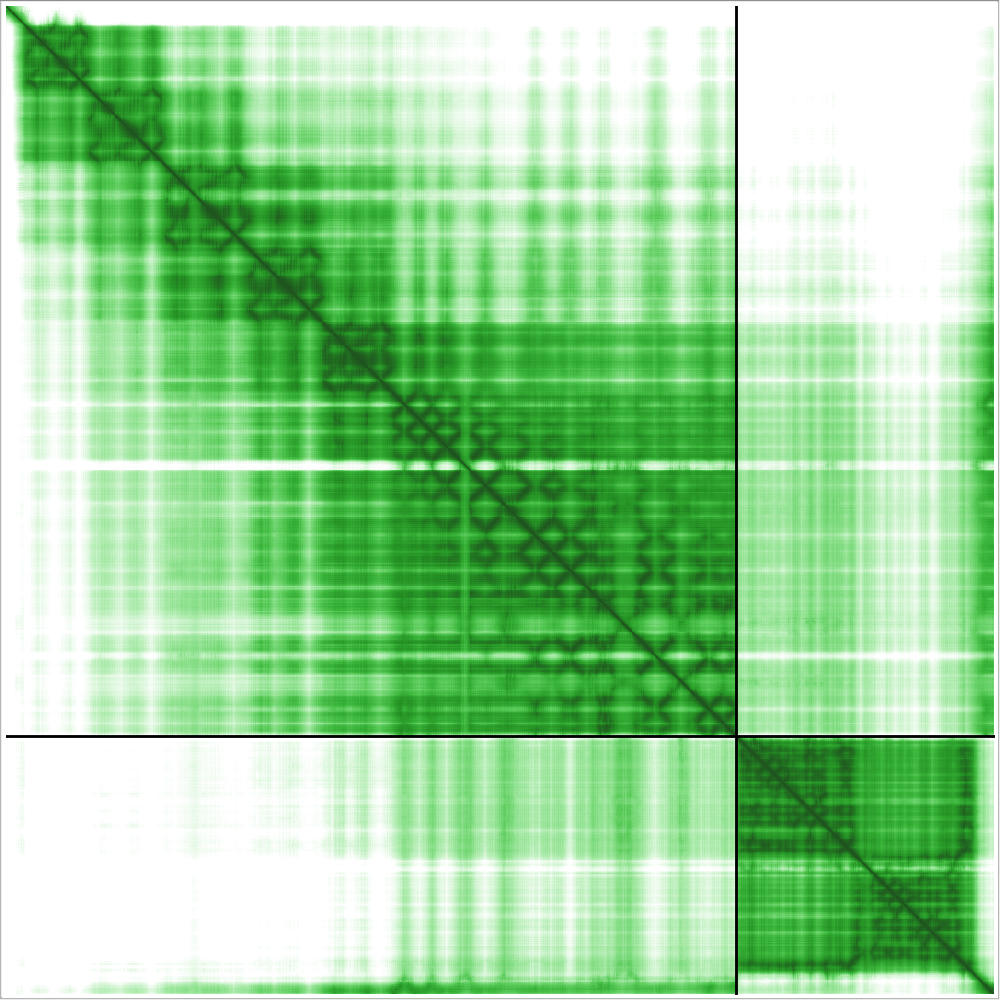

### fold_bama34898_llpa_b74_full_data_0.png

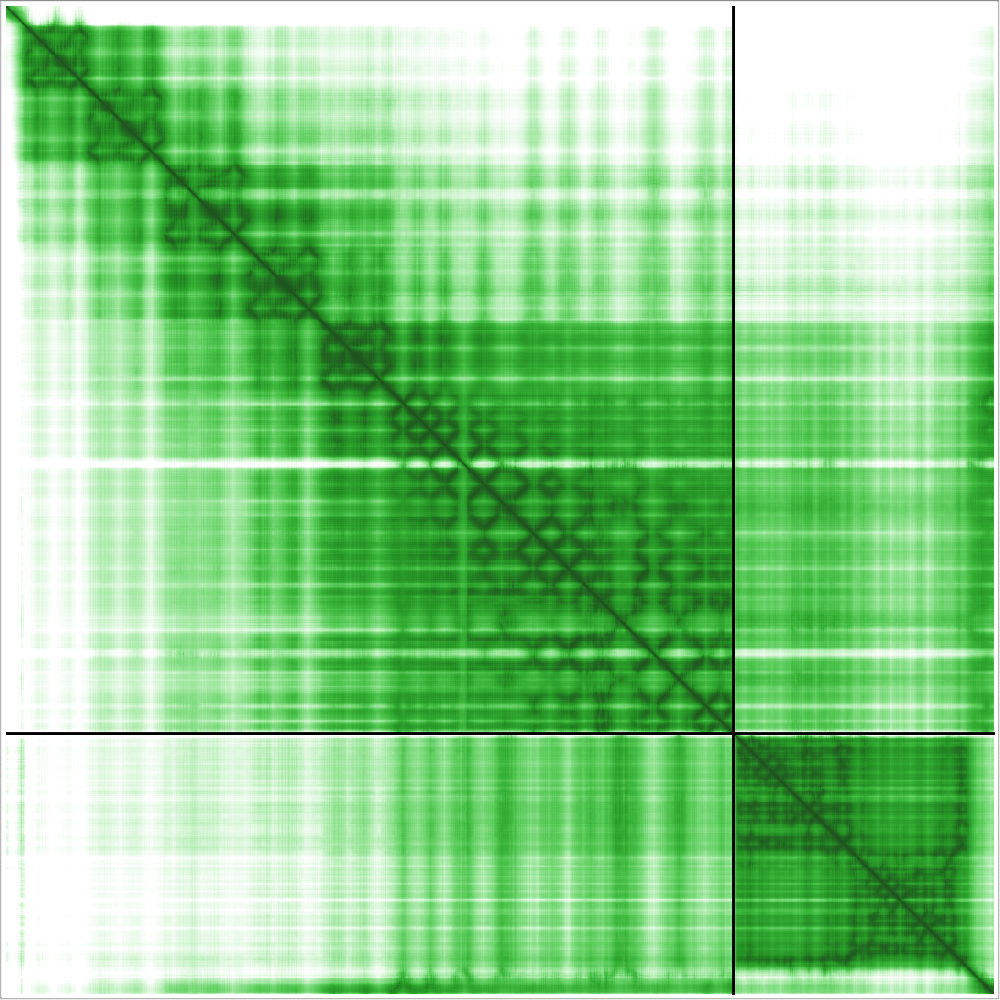

### fold_bama69818_llpb_a26_full_data_0.png

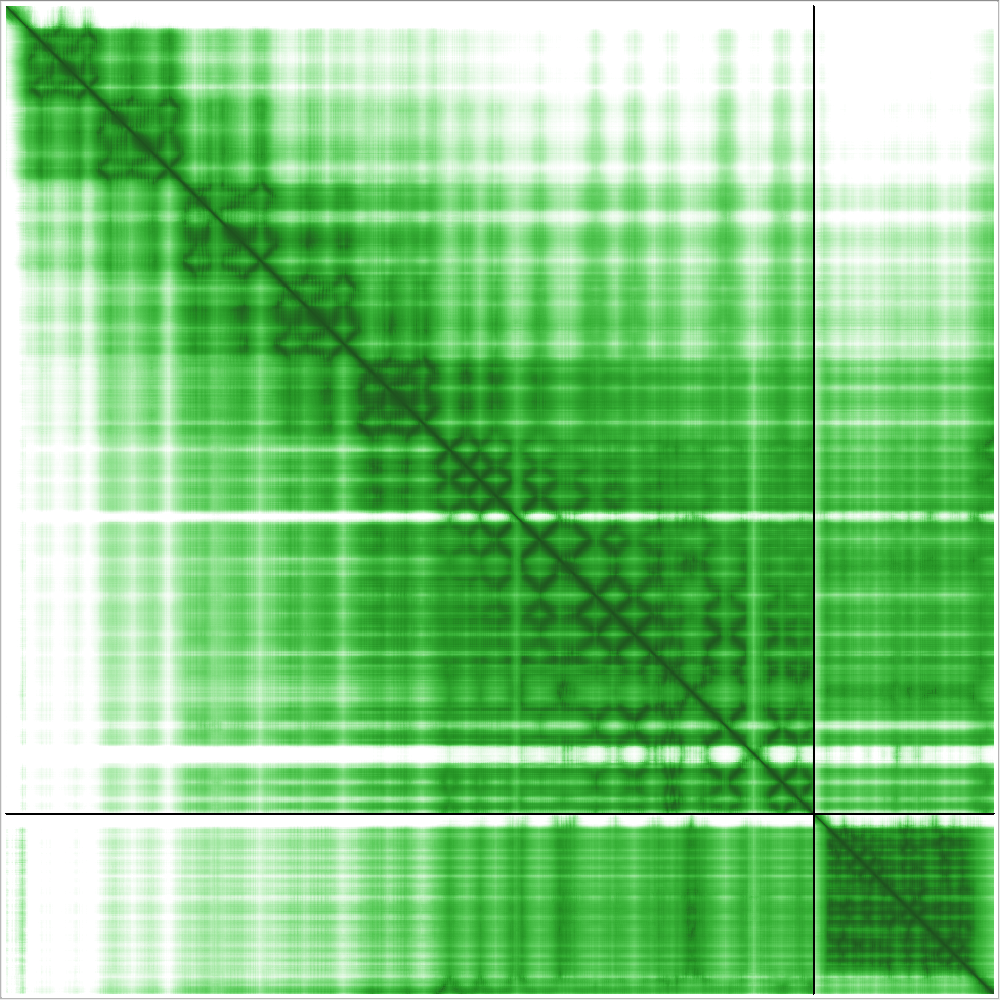

### fold_bama76125_llpb_a49_full_data_0.png

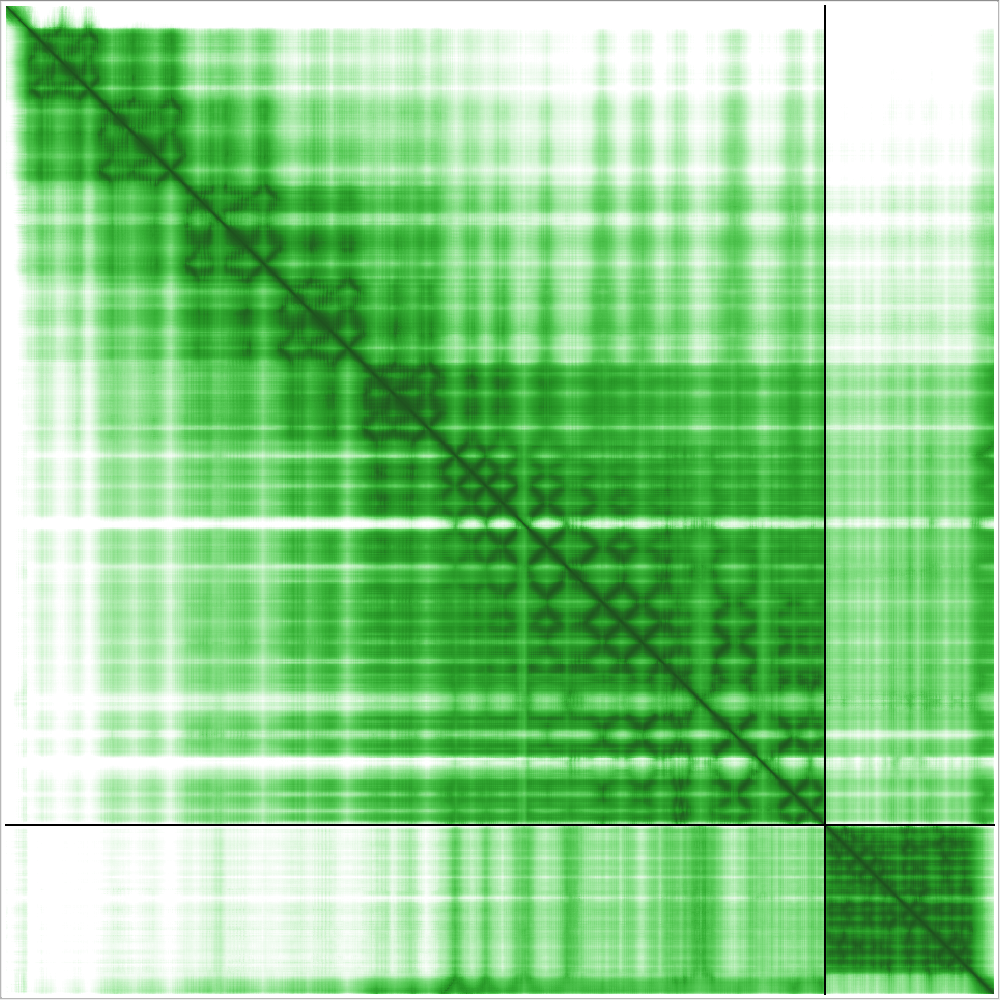

### fold_bama76125_llpb_a90_full_data_0.png

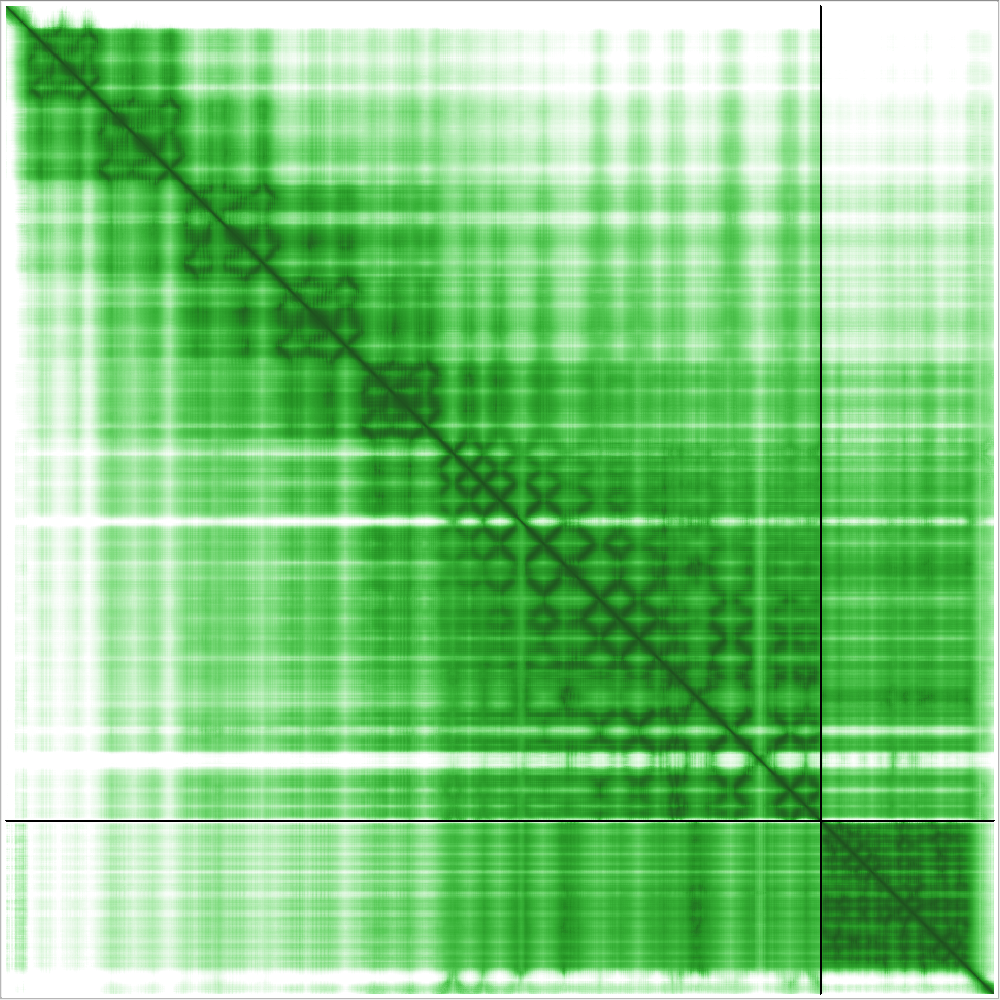

### fold_bama76125_llpb_b57_full_data_0.png

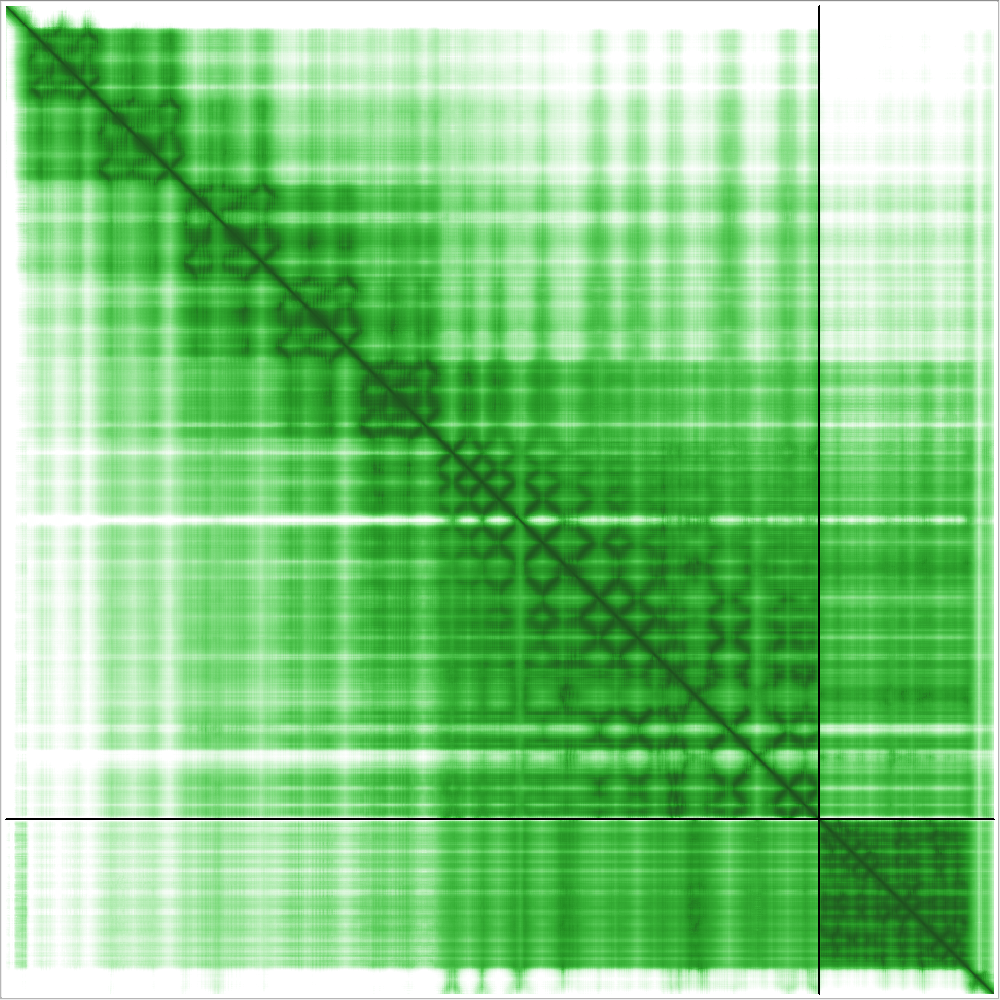

### fold_bama76125_llpb_c32_full_data_0.png

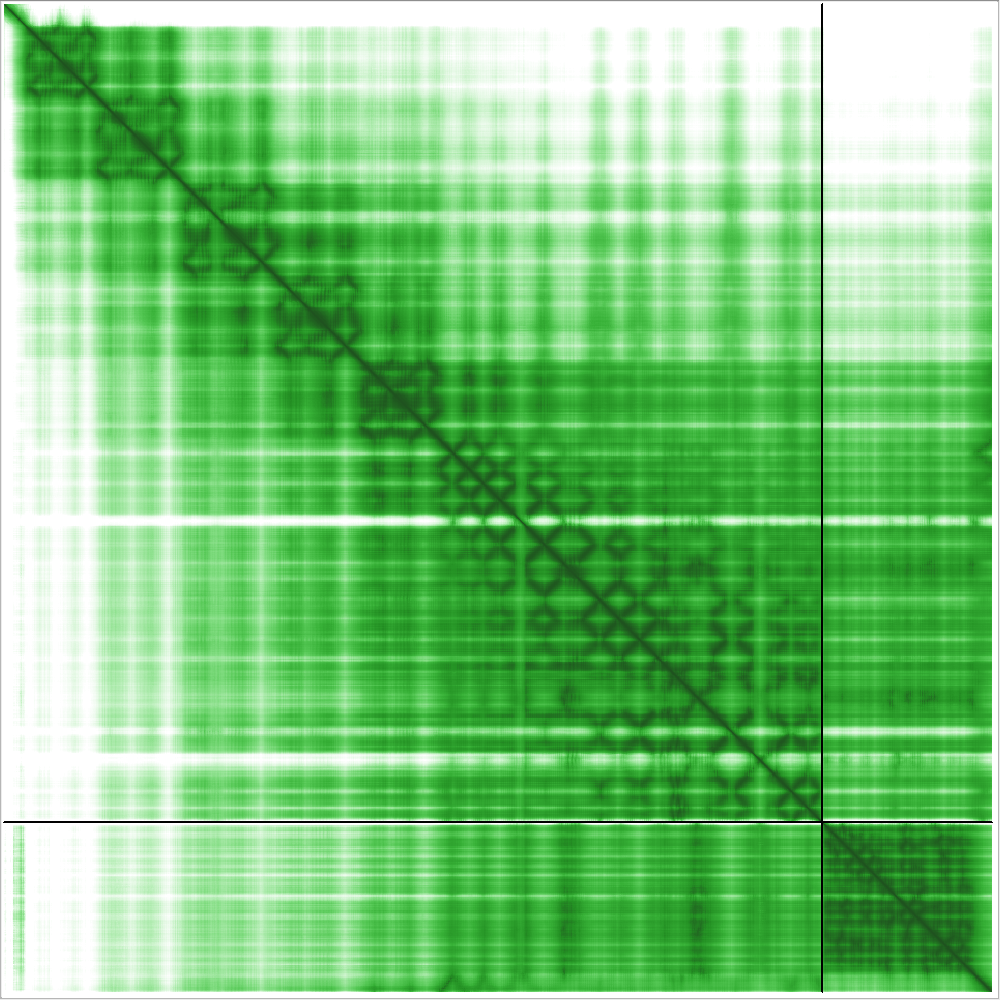

### fold_bama76129_llpb_c02_full_data_0.png

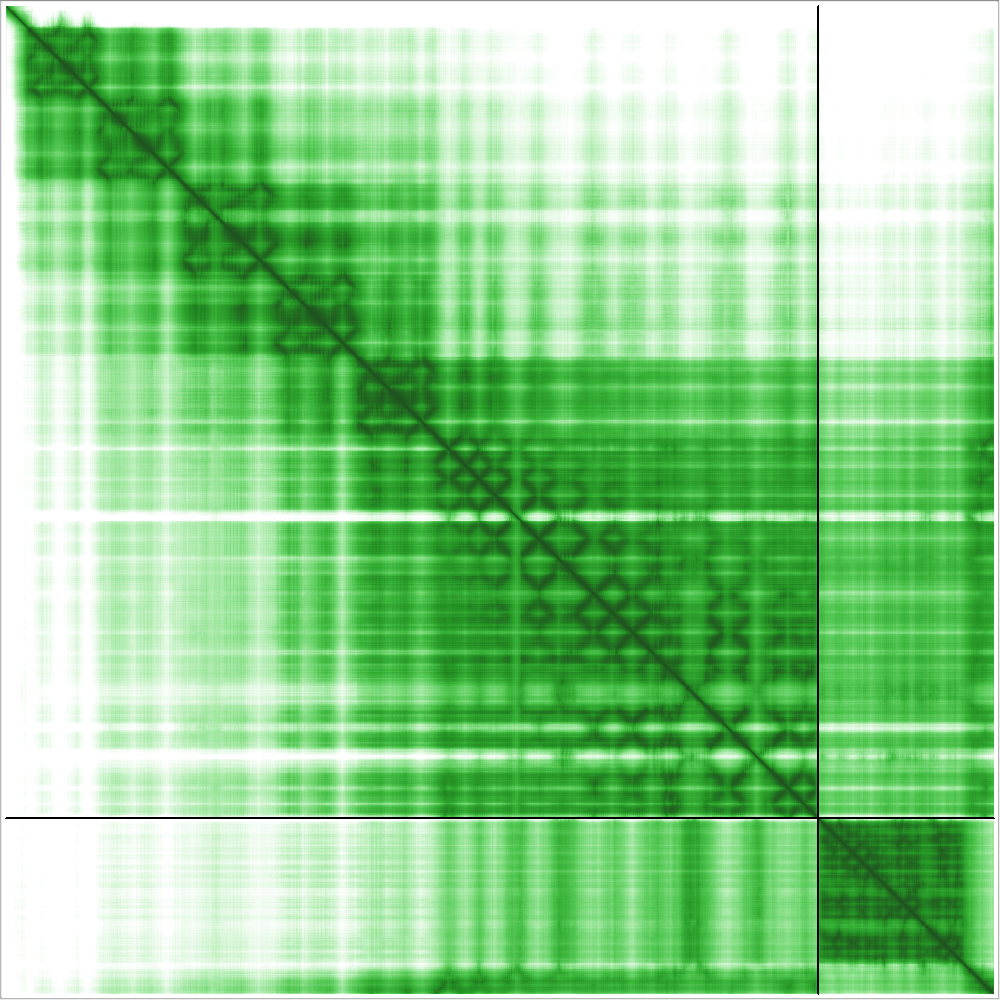

### fold_bama76129_llpb_c06_full_data_0.png

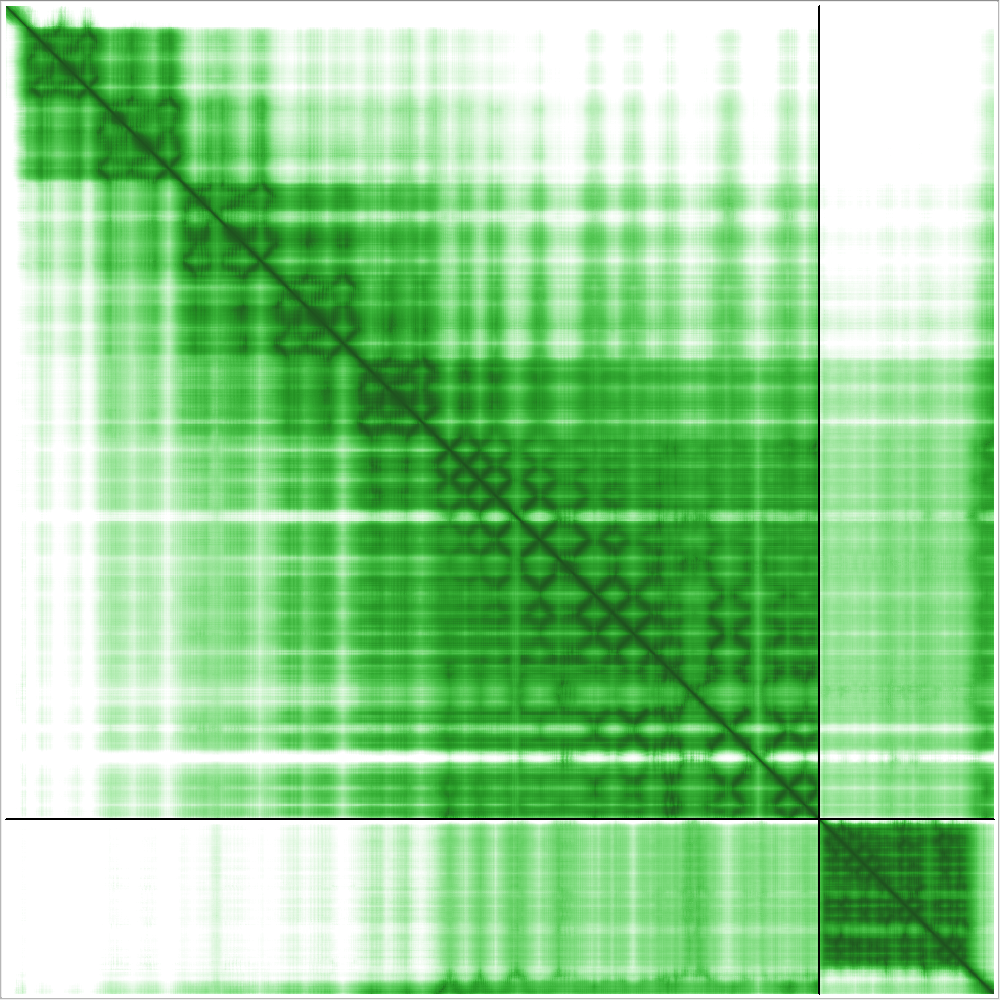

### fold_bama76129_llpb_c11_full_data_0.png

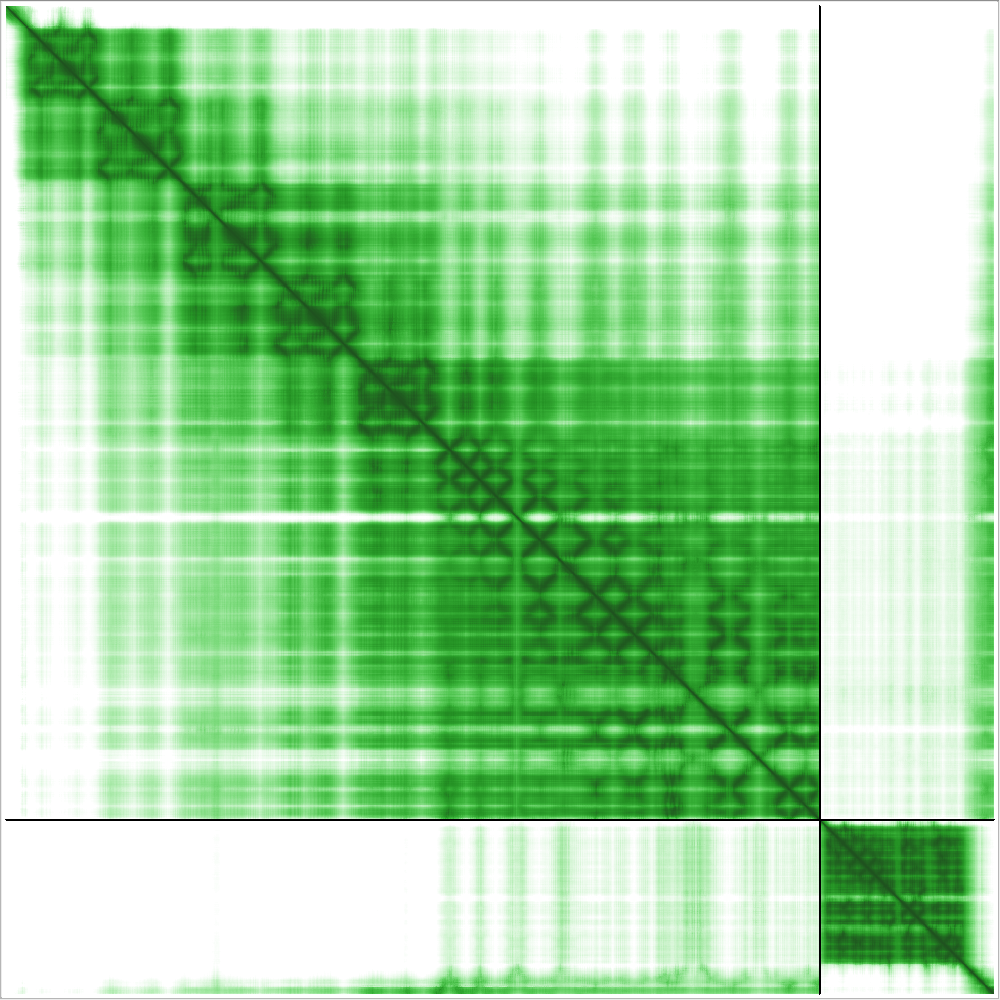

### fold_bama76585_llpb_a67_full_data_0.png

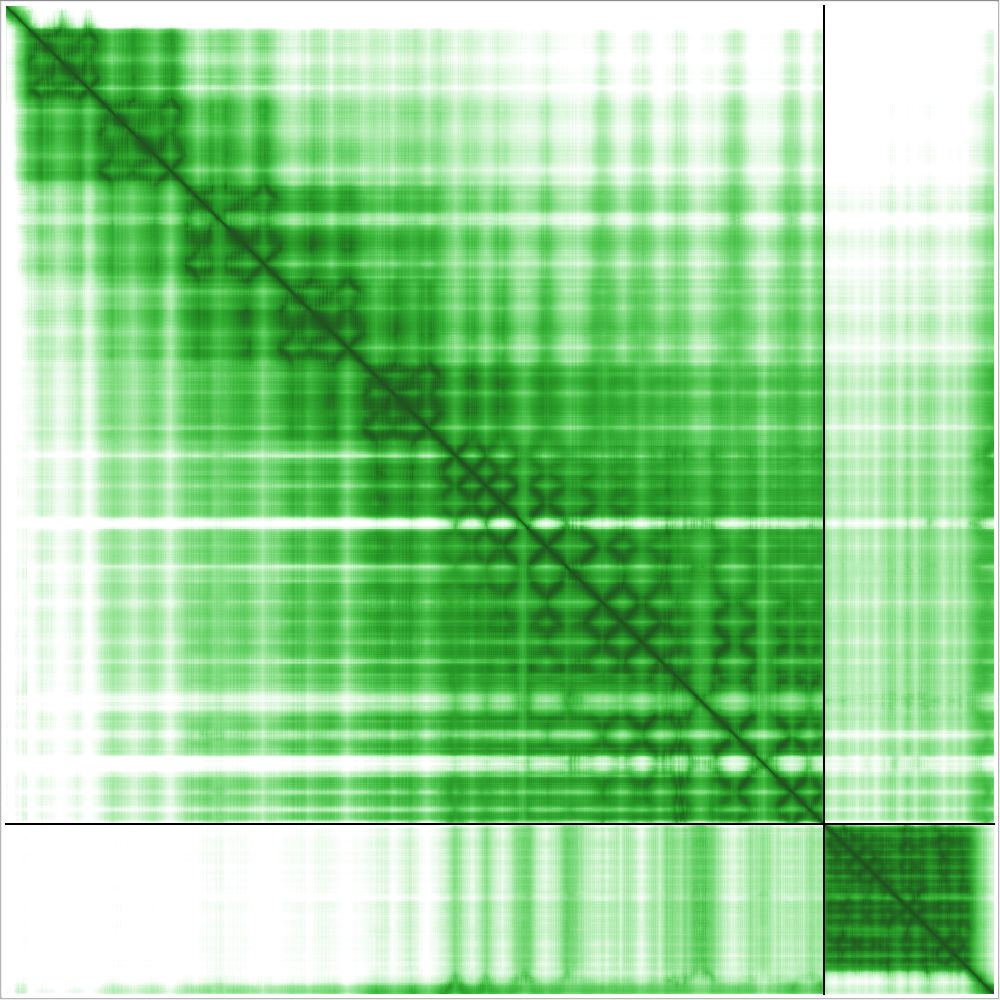

### fold_bama85048_llpb_a63_full_data_0.png

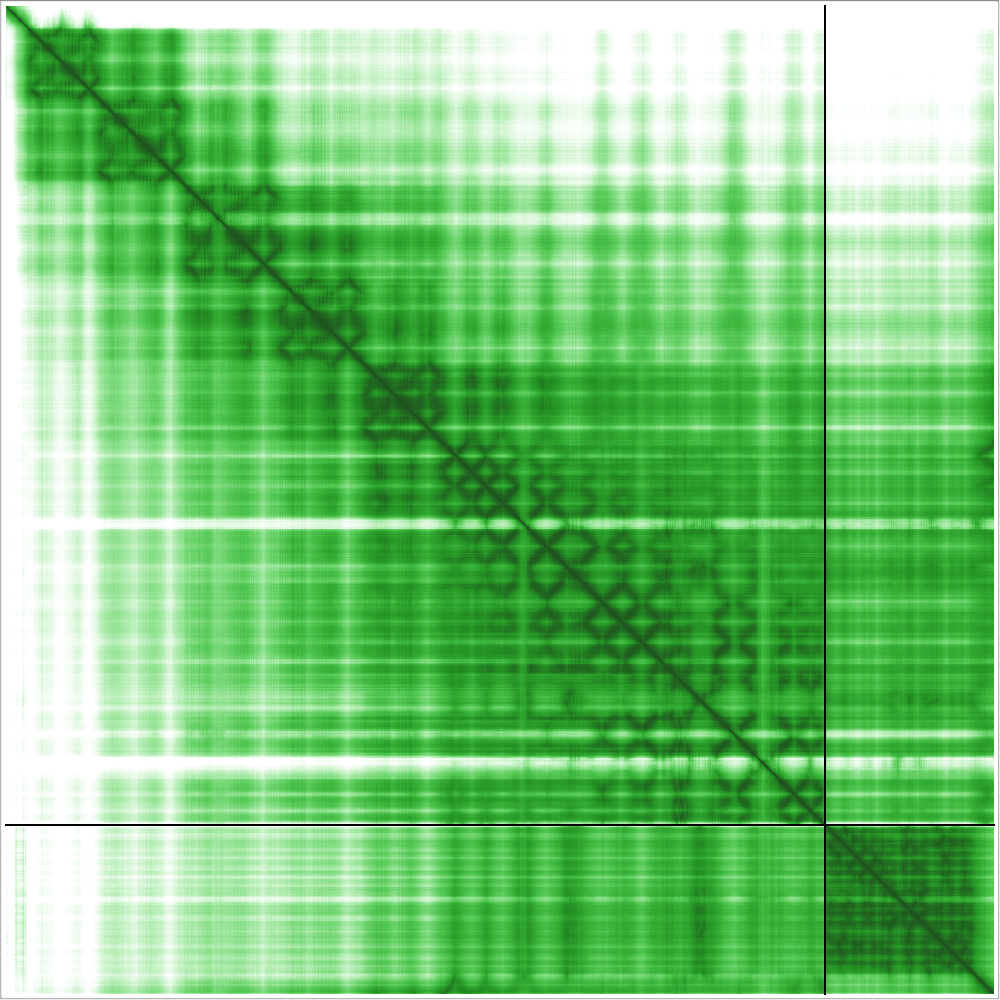

### fold_bama_69857_llpb_b86_full_data_0.png

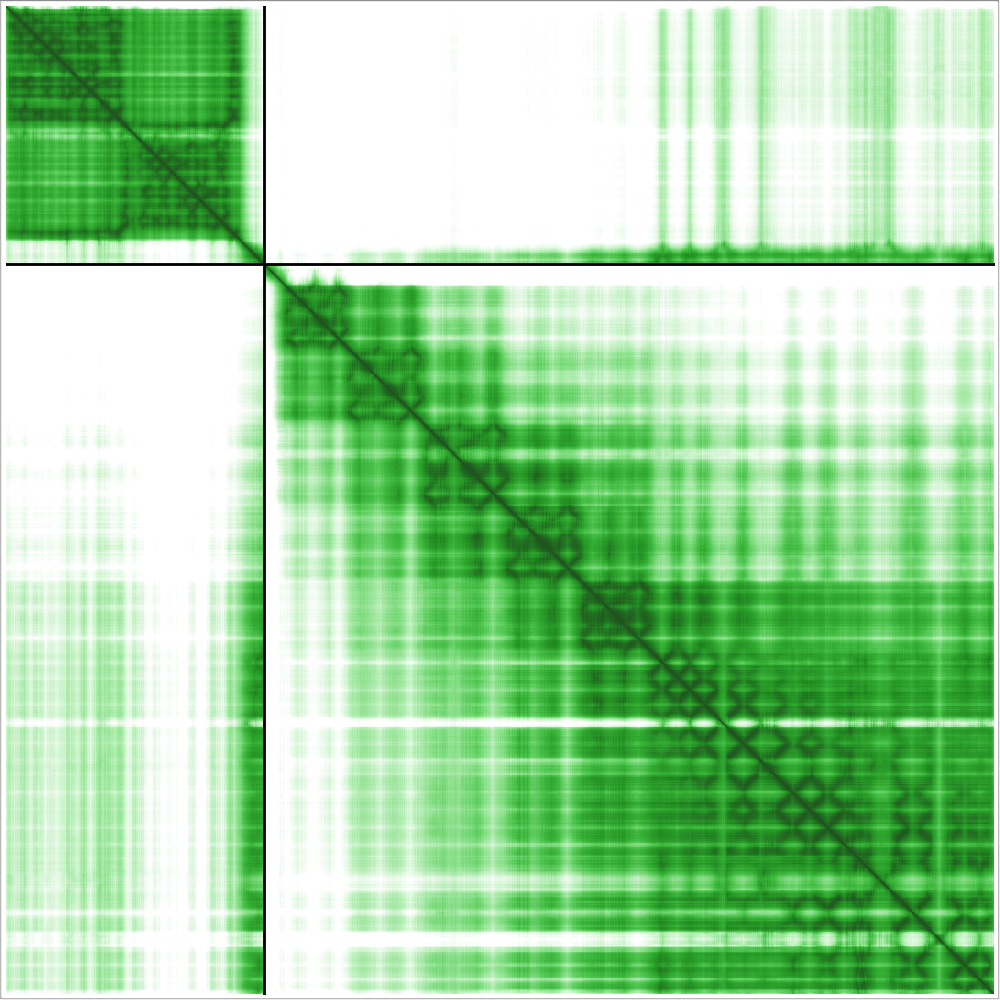

### fold_bama_69857_llpb_c27_full_data_0.png

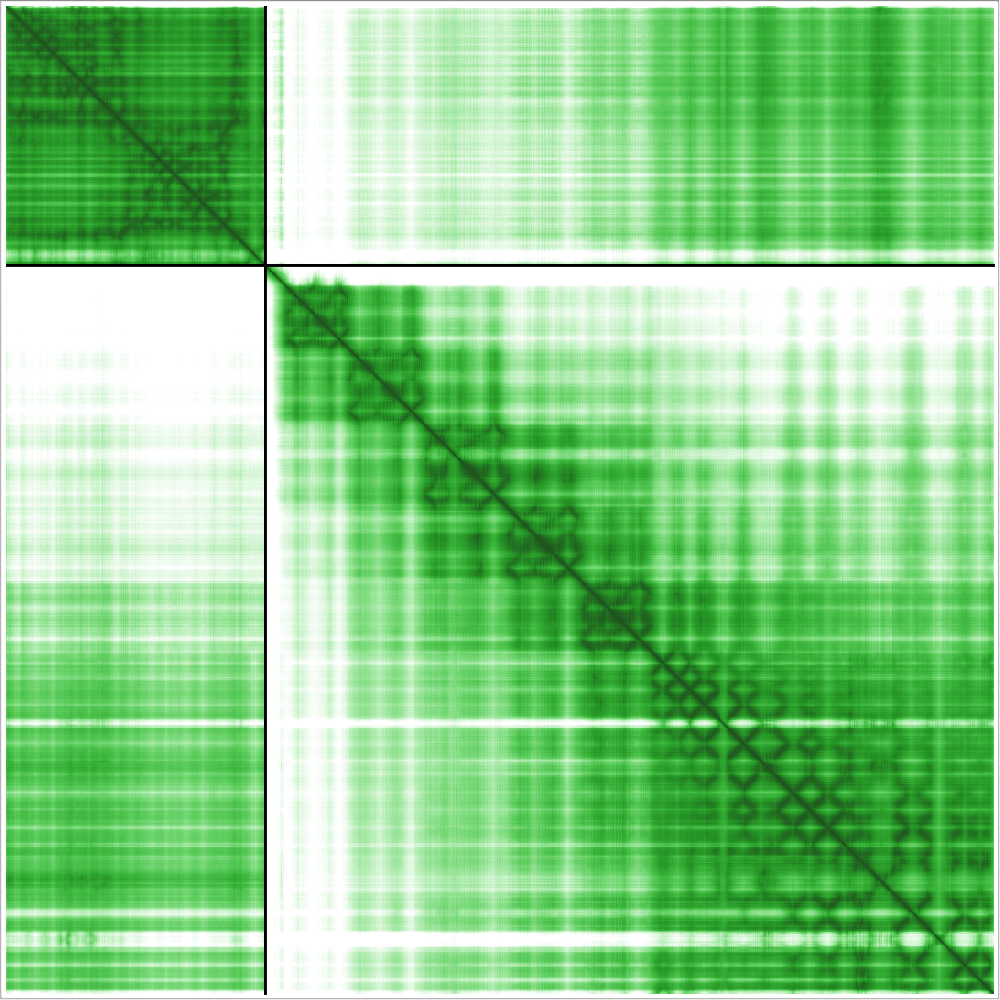

### fold_bama_77212_llpa_c27_full_data_0.png

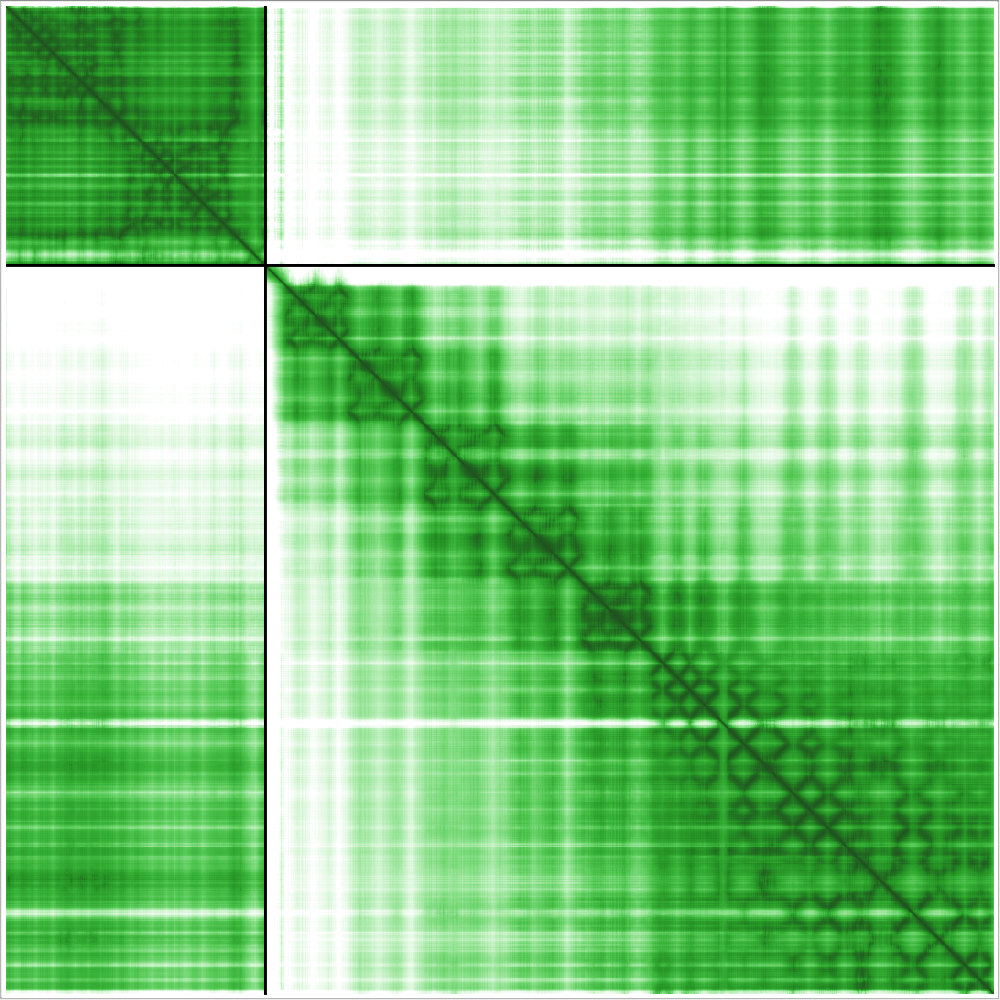
